# Sexual reproduction is controlled by successive transcriptomic waves in *Podospora anserina*

**DOI:** 10.1101/2024.12.09.627484

**Authors:** Frédérique Bidard, Pierre Grognet, Gaëlle Lelandais, Sandrine Imbeaud, Marie-Hélène Mucchielli-Giorgi, Robert Debuchy, Véronique Berteaux-Lecellier, Fabienne Malagnac

## Abstract

Despite the inherent challenge of finding suitable mating partners, most eukaryotes use sexual reproduction to produce offspring endowed with increase genetic diversity and fitness. The persistence of this mode of reproduction is a key question in evolutionary biology. Fungi offer valuable insights into this question, due to their diversity and short lifecycle. This study focuses on *Podospora anserina*, a pseudohomothallic ascomycete fungus that bypasses self-sterility by maintaining two compatible nuclei in one mycelium. We performed genome-wide gene expression profiling during ten stages of *P. anserina* sexual reproduction and identified five major expression patterns. Our expert annotation approaches identified differentially expressed genes related to secondary metabolite production, fungal vegetative incompatibility, programmed cell death, and epigenetic regulation. In addition, master transcriptional regulators and their target networks were uncovered. This study provides a comprehensive database for future functional genomics experiments and novel pathway characterization during sexual reproduction.

## Introduction

Most eukaryotes rely on sexual reproduction to produce offspring. However, the maintenance of sex remains a fundamental question in evolutionary biology. The amazing diversity of fungi makes them of particular interest for addressing this question. Fungi are found in almost every terrestrial and marine ecosystems (Stajich *et al*. 2009; Breyer and Baltar 2023). Some play key roles in biogeochemical cycles, while others possess traits that enable them to be pathogenic to plants or animals (Iliev *et al*. 2024). Understanding the biology of fungi, and in particular why many of them can produce either asexual or meiotic products depending on environmental conditions (Dyer and Kück 2017; Wang *et al*. 2018), is crucial for fields as diverse as ecology, agronomy and human health.

For fungi, as for all eukaryotes, sexual reproduction has well-documented advantages. In particular, meiosis, the sex-specific division, reduces the accumulation of deleterious mutations and can link beneficial alleles that are subsequently fixed in populations by natural selection (Otto 2009). In harsh environments, sexual reproduction, as opposed to asexual reproduction, can lead to improved fitness of the offspring.

Fungal sexual reproduction relies on a wide range of developmental strategies and mating systems. These include both inbreeding, in which a single individual can self-fertilize, and obligate outbreeding, in which only genetically distinct compatible partners can mate. Some fungi, referred to as pseudohomothallic, have managed to retain the best of each strategy. Pseudohomothallism extends the diversity of reproductive systems evolved in fungi by selecting breeding systems that allow self-sterility to be bypassed by maintaining two populations of compatible nuclei in the same mycelium (Peraza-Reyes and Malagnac 2016). Early forward genetic screens and more recent transcriptomic studies in Sordariomycetes (Wang *et al*. 2014; Lütkenhaus *et al*. 2019; Kim *et al*. 2022) have led to the identification of large sets of genes involved in specialized developmental pathways that control the fertilization and the subsequent developmental stages (Wilson *et al*. 2019). Some of these species were heterothallic (self-incompatible, self-sterile (Wang *et al*. 2014; Lütkenhaus *et al*. 2019)), while others were homothallic (self-compatible, self-fertile (Teichert *et al*. 2012; Wang *et al*. 2019)). However, how the development of meiotic tissues within the fruiting body is controlled and which genes are critical in this process remains unknown.

In this study, we propose to approach this question from a different angle using the pseudohomothallic species *Podospora anserina*. This species is used as a model system for the study of the fruiting body development (Peraza-Reyes and Malagnac 2016) but a detailed analysis of the cellular programs allowing its formation from fertilization to ascospore discharge was lacking. In this context, we performed a genome-wide gene expression profiling during the sexual reproduction of *P. anserina* at ten time points representing key developmental steps. To characterize the topology of differential gene expression profiles across sexual reproduction, we analyzed our transcriptomic data using two complementary approaches: *i)* a data-driven approach using k-means clustering and *ii)* a biological expertise-driven approach to mine the gene functions. The data-driven approach revealed five major patterns (waves) of differential expression during sexual development. The expert annotation approach has led to the identification of a number of interesting gene categories, including secondary metabolites and toxins encoding genes, genes responsible for fungal vegetative incompatibility programmed cell death, and gene related to epigenetic effectors that interfere with messenger stability (RNAi/MSUD genes. Finally, we identified several master transcriptional regulators and draw their potential target gene network. Overall, our study has produced a comprehensive database that offers numerous candidates for functional genomics experiments. This database also paves the way for the characterization of new pathways during sexual reproduction.

## Materials and methods

### *Podospora anserina* strains and methods

*P. anserina* is a filamentous ascomycete whose life cycle and general methods have already been described (Esser 1974; Rizet and Engelmann, 1949; Zickler *et al*., 1995). All strains used in this study were derived from the *S* strain (Rizet and Engelmann 1949; Espagne *et al*. 2008). Growth and crosses of *S mat+* and *S mat-* strains were carried out on a minimal synthetic medium (M2 medium) at 27°C with constant light (Silar 2020). The *136* strain is described in (Coppin and Silar 2007).

### Cytology

Sexual cells were either fixed in 7.4% paraformaldehyde and processed for microscopy as previously described (Thompson-Coffe and Zickler, 1994) or directly observed in M2 medium. When required, nuclei were stained with DAPI (4’,6-diamidino-2-phenylindole, 1 mg/mL; Boehringer Ingelheim). Observations were performed with a Zeiss Axioplan microscope, and images captured with a CDD Princeton camera system.

### Time-course procedure and RNA isolation

To facilitate biological materials collecting, we used cellophane sheets (cat # 1650193, Bio Rad Laboratories, Hercules) and cheesecloth pieces (Nitex 03-48/31, Sefar AG, Heiden), which were deposited on M2 medium prior to inoculation. Cellophane sheets avoid the contamination of biological material by M2 medium when scraping mycelium. The mesh of cheesecloth accommodates the mycelium, while perithecia develops above the mesh and can be scraped with minimal mycelium contamination. S mat- strain has been used as male gamete donor and *S mat*+ strain as female. The *S mat*+ strain was grown on cellophane sheet for the analysis of mycelium and on cheesecloth for analysis of perithecial development. The biological material was then collected 6, 12, 18, 24, 30, 42, 54 and 96 hours after fertilization and an additional time point 24 hours prior fertilization was taken. Three biological replicates issued from three independent time course have been collected for all time points and two for T42, as we discarded the third and the back-up replicates, which did not meet our quality requirements.

For each ten points of the time course, the number of Petri dishes has been adjusted (five were used for T-24h, T0h, T6h, T18h, T24h, four for T30h, two for T42h, one for T54h and T96h) to collect 20 to 100 mg of fertilization competent mycelium or/and perithecia. One to five Petri dishes *per* time course experiment were thus inoculated simultaneously with a growing *S mat-* strain and spermatia were collected by washing with 1.5 ml of H_2_O six days later. The collected spermatia (around 10^4^ spermatia per ml) were pooled and then used to fertilize a corresponding number of Petri dishes of 4-day-old *S mat+* strain. The moment just prior the fertilization process was considered as the T0 (0 hours).

The biological material collected by scraping gently either cellophane or cheesecloth with a cover glass, was dried out with filter paper, weighted, frozen in liquid nitrogen and stored at −80°C. Total RNAs of *P. anserina* were extracted using RNeasy Plant Mini Kit (Qiagen), including a grinding process using a Mikro-Dismembrator (Sartorius) and a DNase treatment. The quality and quantity of the total RNAs was determined by using a Nanodrop spectrophotometer (Nanodrop Technologies) and the Bioanalyzer 2100 system (Agilent Technologies) as previously described in (Imbeaud *et al*. 2005).

### Gene expression array-based hybridization

Transcriptome microarray experimental procedure, *i.e.* targets preparation, hybridization and washing, was done following the two-color microarray-based gene expression analysis instructions (version 5.0, February 2007) as described by the manufacturer (Agilent Technologies). One-μg aliquots of total RNA were labelled using the Low RNA input fluorescent linear amplification (LRILAK) PLUS kit (Agilent Technologies); internal standards came from the Two-color RNA spike kit (Agilent technologies). The labelling efficiency and the product integrity were checked as described by (Graudens *et al*., 2006). The reference cRNA was labelled with Cy5 and sample probes with Cy3. Briefly, the reference RNA pool was prepared by mixing equal amounts of RNA extracted from mycelium at 48 h and 96 h post-inoculation, and from perithecia at 24 h, 48 h and 96 h post-fertilization (Bidard *et al*. 2010). This composition allows a detectable and stable expression signal on the majority of CDS (Bidard *et al*. 2010). Then, Cy3- and Cy5-labeled targets were mixed and incubated on an Agilent microarray slide for 17 hours at 65°C, in a rotating oven (6 rpm for a 1×44K and 10 rpm for a 4×44K array format), using an Agilent *in situ* hybridization kit. The slides were washed and then any traces of water were removed by centrifugation at 800 rpm for 1 min (Bidard *et al*. 2010; Bidard *et al*. 2011).

### Data acquisition from microarray experiments

Microarrays were scanned using an Agilent dual laser DNA microarray scanner (Agilent Technologies), model G2567AA, with 5-µm resolution. The resulting 16-bit image files were analyzed in the *Feature Extraction* processing system (FE, v9.5.3). Spot and background intensities were extracted and normalized with the Feature Extraction software using the GE2-v4_95_Feb07 default protocol (Local background subtracted and LOWESS normalization). Preliminary array quality was assessed using Agilent control features as well as spike-in controls (Agilent 2-Color Spike-in Kit for RNA experiment). Subsequent flagging was done according to the GenePix Pro software (Molecular Devices Syunnyvale, CA, USA) nomenclature, including four levels of flags (good (100), bad (−100), not found (−50), moderate (0)) and raw data normalization was performed as previously described in articles such as (Bidard *et al*. 2010; Bidard *et al*. 2011). Briefly, it consisted in using standard bioinformatics protocols to first filter probes with good quality flags, then correct signal intensities considering global and local background noises, and finally correct for systematic bias between Cy3 and Cy5 signals.

### Graphical summary of data analysis

A schematic representation of the obtained data set is shown in Supplementary Figure 1 (see number #1). Gene expression measurements were thus analyzed according to two different strategies, allowing to study either the co-expression between any pairs of annotated genes in the *P. anserina* genome (see number #2) or the differential expression for all annotated genes in the *P. anserina* genome between any pairs of time points (see number #3). For this, gene expression profiles were calculated and statistical analyses to search for differentially expressed genes were performed (see the following sections for more details).

### Calculation of gene expression profiles

For each gene, “A values” were computed at each time point and for each replicate, with the following formula: 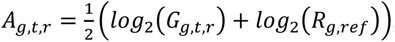, where *G_g,t,r_* represents the normalized Cy3 intensity value measured for gene *g* at time point *t* in biological replicate *r*, and *R_g,ref_* represents the normalized Cy5 intensity value measured for gene *g* in the reference RNA pool (see previous sections). At each time point, triplicated values were next averaged to obtain one value per time point, noted *A_g,t_*. The mean and the median values, respectively noted *A_mean_* and *M_g_*, and associated with *A_g,t_* values over all time points, were finally calculated and used to adjust expression measurements for each gene around a new median value of 0, *i.e.* 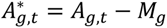. An expression profile for a gene *g* is therefore made with ten *A*\* values which are: 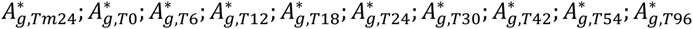. Notably, it represents the expression fluctuations of the gene over the time course.

### Statistical analyses to search for differentially expressed genes

Gene expression measurements between pairwise time points were compared using the MAANOVA (MicroArray ANalysis Of Variance) package (version 1.14.0) (Wu *et al*., 2003). It allows fitting an ANOVA model as proposed by (Kerr and Churchill 2001) and provides a way to consider multiple sources of variation in the microarray experiments. P-values were computed by permutation analysis and were adjusted by the false discovery rate step-up method of Benjamini and Hochberg to account for multiple testing (Benjamini and Hochberg, 1995). From all comparisons, lists of differentially expressed genes (Table S1) were defined using several filtering strategies:

- <u>Filter #1</u> to select the genes with *(i)* an adjusted p-value < 0.05, *(ii)* a fold change ≥ 4 or ≤ −4 with a *A_mean_* ≥ 4.5,
- <u>Filter #2</u> to select the genes with *(i)* an adjusted p-value < 0.001 and *(ii)* a fold change ≥ 2 or ≤ −2 (*i.e.* absolute value of log2(fold change) ≥1).

The two lists of genes were used respectively for unsupervised clustering (see below) and biological interpretations (see the Results section).

### PCA analysis and unsupervised clustering based on gene expression profiles

Expression profiles of all genes were used for principal component analysis using the function “pca” available in the R package mixOmics (Table S2) and expression profiles of differentially expressed genes (selected based on Filter #1, see previous section) were used for cluster analyses. To this list, the expression profiles of genes coding for mating-type transcription factor SMR1, SMR2 and FMR1 were added. A hierarchical clustering (UPGMA method) and several K-means clustering were computed independently using Spotfire DecisionSite software package (Spotfire, Somerville, Mass.) with correlation coefficient as similarity measure between *A*\* values in gene expression profiles. Intersection of clusters obtained by the two-clustering method results in robust clusters (Table S1) from which the temporal waves (see Result section) were derived.

### Sequence comparisons to search for orthologous genes with other species

Orthologous genes were searched between *P. anserina* and the other species *Neurospora crassa*, *Sordaria macrospora*, *Chaetomium globosom* and *Trichoderma reesei*. For this purpose, protein sequences were downloaded from Uniprot (https://www.uniprot.org/) under the accession numbers: UP000001805_367110 (*N. crassa*), UP000001881_771870 (*S. macrospora*), UP000001056_306901 (*C. globosum*) and UP000008984_431241 (*T. reesei*). The OrthoFinder algorithm (Emms and Kelly 2019) was used with default parameters to obtain “orthogroups” (Table S3), *i.e*. groups of genes that are good candidates to share a common ancestor between the compared species.

### Functional enrichment of terms associated with lists of genes

Gene Ontology (Ashburner *et al*. 2000; The Gene Ontology Consortium *et al*. 2023) and Pfam annotations (Mistry *et al*. 2021) were used to explore the biological relevance of the lists of genes that composed the temporal waves (Table S4 and S5). P-values were calculated using Fisher’s exact test and following the methodology described in (Boyle *et al*. 2004).

### Expert annotation of *P. anserina* genes

Expert annotation (Table S6) was performed using either BLASTP reciprocal best-hit and/or Pfam conserved motif analyses. Output results were all validated after careful manual inspection including the use of the FungiBD database for orthologs verifications (Basenko *et al*. 2018).

### Co-expression network based on expression profiles of genes coding for transcription factors

Expression profiles of differentially expressed genes (selected based on Filter #2, see previous section) for which the expert annotation was associated with “transcription factors” (Table S7) were used to search for significant correlation (> 0.95) with the expression profiles of all genes previously classified into transcriptional waves. Selected gene pairs were used to represent a co-expression network as defined in (Wolfe, Kohane, and Butte 2005), in which all potential links of transcriptional control, between the TF and another gene (referred to as “target gene”) are shown.

### Availability of data and source code, website to explore the results

The raw data have been deposited in the MIAME compliant Gene Expression Omnibus database (Edgar, Domrachev, and Lash 2002) and is accessible through the GEO Series accession number GSE21659 https://www.ncbi.nlm.nih.gov/geo/query/acc.cgi?acc=GSE93094. The source code has been deposited at: https://github.com/Podospora-anserina/transcriptome_kinetics_development/). To explore the results for a particular gene or for a set of genes, we have created a WEB application which is available at: https://pierregrognet.shinyapps.io/cinetique_app/.

## Results

### High-resolution kinetics of transcriptional fluctuations during sexual reproduction

To ensure an accurate, high-density representation of transcriptional variation, we collected RNA samples at ten key time points during the sexual reproductive cycle of *P. anserina* (Fig. 1).

**Figure 1 -.**
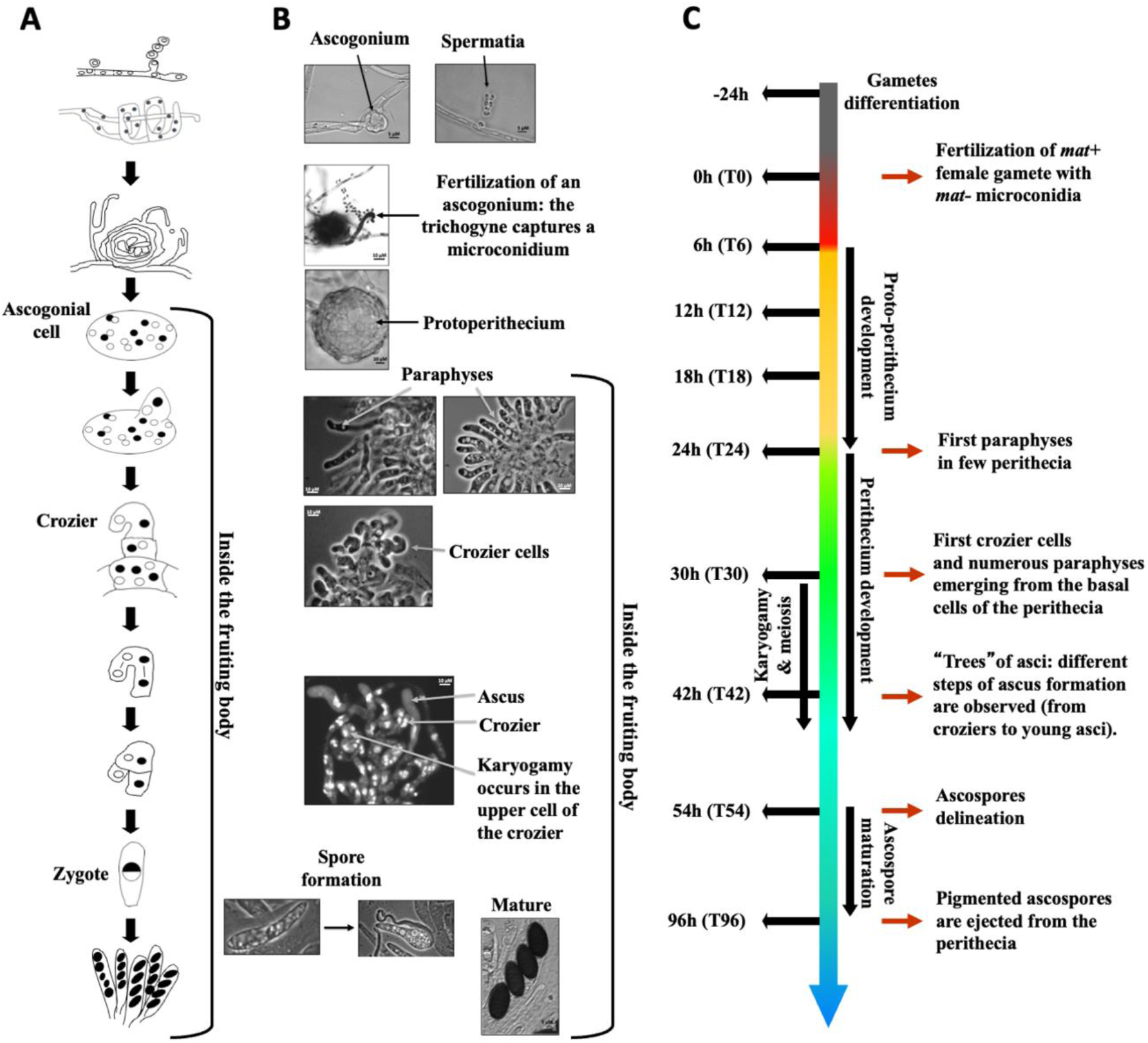
*P. anserina* developmental time course from fertilization to ascospore maturation. **A.** Schematic representation of the fertilization and the main steps of zygotic development. **B.** Pictures of the main steps of sexual development. Sexual reproduction began with the production of male (spermatium) and female (ascogonium) gametes. A pheromone/receptor signaling system allows the female gametes to attract specialized hyphae (trichogynes) that recognize and fuse with male gametes of compatible mating type. This event initiates the multi-step process of sexual development that begins with the formation of fructifications and ends with the discharge of ascospores. The nucleus of the male gamete enters the trichogyne, migrates along this specialized hypha, reaches the ascogonium and undergoes several mitotic divisions. Fertilization (T0) triggers the development of protoperithecia (T6 to T24), which recruit multiple layers of protective maternal hyphae to grow into perithecia (T24 to T45). The developing fructifications shelter the fertilized ascogonial cells (T24 to T30), which contain multiple haploid parental nuclei of either *mat*+ or *mat*- genotype. Differentiation of paraphyses begins (T24) just before three-celled hook-shaped structures called croziers emerge from the ascogonial cells (T30 to T42). Croziers are dikaryotic because they contain two haploid nuclei derived from both parents. Karyogamy (T30 to T44) occurs in the upper cells of croziers. The resulting diploid zygote immediately undergoes meiosis (T30 to T44), producing four haploid nuclei, which then undergo mitosis prior to ascospore delineation (T54 to T96). Ascospores are formed around two non-sister nuclei within the developing asci. In rare cases, two ascospores form around a single haploid nucleus, resulting in five-spored asci. **C.** Time line of the sexual cycle of *P. anserina* with the steps from which RNAs have been extracted on the left.

*P. anserina* is a pseudohomothallic fungus, namely natural isolates are self-fertile because they contain nuclei of each mating-type. Self-fertilization does not start at a specific time point and therefore natural isolates cannot be used for a time course analysis of sexual development. However, the genetic basis of the *P. anserina* mating system is heterothallism with two mating-type idiomorphs called *mat+* and *mat-* (Turgeon and Debuchy 2007). Typical ascospores contain *mat+* and *mat-* nuclei, but a few asci contain monokaryotic ascospores which produce self-sterile individuals. Those latter individuals were used in the experiments presented here. Fertilization was performed by spreading *mat-* male gametes on a *mat+* strain. This experimental design allows the time course to start at a well determined time point for synchronization purpose. Under this experimental condition almost all fertilization events occurred within the first six hours (Fig. S2A and S2B). However, despite this experimental design, RNA samples were extracted from biologically non-homogeneous tissues because *P. anserina* fructifications are composed of different cell types (Fig. 1A and 1B).

We measured the expression of 10,507 nuclear genes at ten time points which are designated as Tm24 for mycelium 24 h before fertilization and T0, T6, T12, T18, T24, T30, T42, T54 and T96 for time 0 to 96 h postfertilization, respectively (Fig. 1C, Table S1). We performed a Principal Component Analysis (PCA) on the complete dataset. More than 75% of the total variance was captured by only three principal components (PCs, Fig. 2A and 2B, Table S2).

**Figure 2 -.**
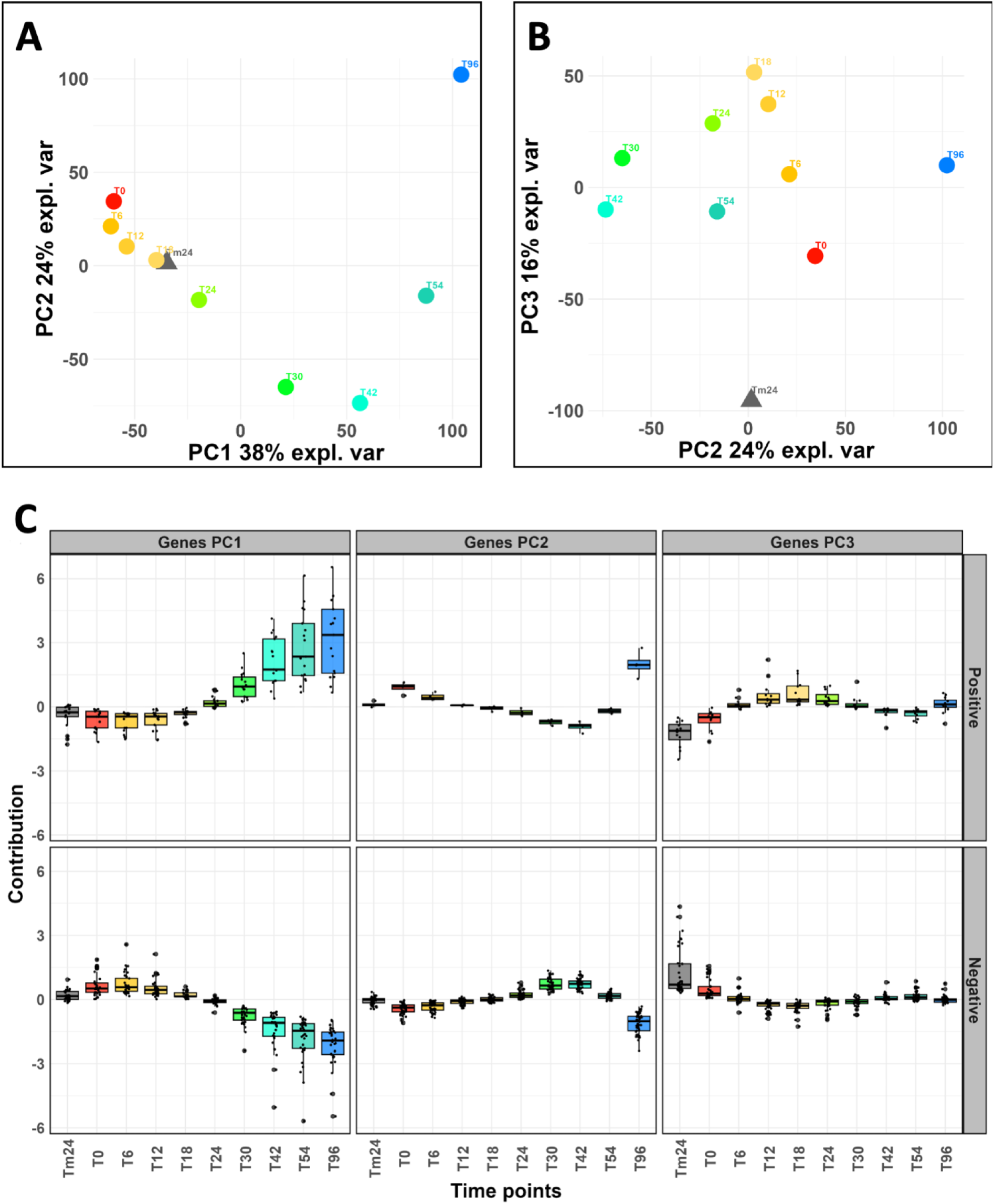
PCA analysis reveals a strong temporal effect on gene expression variations relevant to the key developmental steps in *P. anserina*. Principal Component Analysis (PCA) on the complete dataset (*i.e*. all gene expression profiles, see Methods). **A.** Axis 1 and 2 of the PCA explain 38% + 24 % = 62% of the variance. **B.** Axis 2 and 3 of the PCA explain 24% + 16 % = 40 % of the variance. Altogether, these first three axes of the PCA explain 78 % of the variance. **C.** Gene expression profiles for the 50 genes, which contribute the most to the definition of the first (PCA1), second (PCA2) and third (PCA3) principal components, as shown in A and B. Detailed lists of these genes are given in Table S2. They were separated according to the type of regulation they experience during sexual development (positive or negative). Tm24, T0, T6, T12, T18, T24, T30, T42, T54 and T96 : time points of the kinetic were the biological samples were collected, described in Fig. 1.

With the exception of Tm24, we observed that all samples were distributed along the PC1 axis (38%) according to the sexual development time line (Fig. 2A). On PC2 axis, we found T30 and T42 samples clustered apart from the others (24%). Interestingly, this period corresponds to the formation of the dikaryotic cells, which are the progenitors of fungal zygotic lineage (Fig. 1). The peripheral position of the Tm24 sample (Fig. 2B) which appeared on the PC3 axis (16%), indicated that the gene expression profiles derived from vegetative mycelium (Tm24) were clearly different from those of all samples collected after fertilization. Overall, this unsupervised exploratory data analysis validated both the relevance of our experimental design and the robustness of our differential analysis (see below). The expression profiles for the 50 genes whose expression measurements contributed the most to the calculation of the principal components 1, 2 and 3 are shown Figure 2C (and detailed Table S2). The data indicated that the greatest fold changes, both positive and negative, occurred after the formation of dikaryotic cells (>T30). This was also the case for PC2, but only after the formation of the ascospores (>T54). We observed that the positive fold change values on PC1 exhibited a higher dispersion after T42.

### Topology of global expression variation reveals five major transcriptional waves during sexual development

Using k-means clustering, we defined 13 representative expression patterns (Fig. S1 and Fig. S3), encompassing 1,186 genes. Further data integration outlined five successive transcriptional waves (Fig. 3, Fig S1 and Table S1). Notably, they all shared a common pivot point (Fig. S4), corresponding to the mid-course of the kinetic (T30-T42). This observation indicated a pause in the regulation of transcriptional activity at the time of dikaryon formation. The 1,186 genes were not evenly distributed across the five waves (Fig. 3 and Table S1), resulting in two periods of intense and equal transcriptional fluctuation (waves II and IV) interspersed with three periods of moderate transcriptional fluctuation (waves I, III, and V).

**Figure 3 -.**
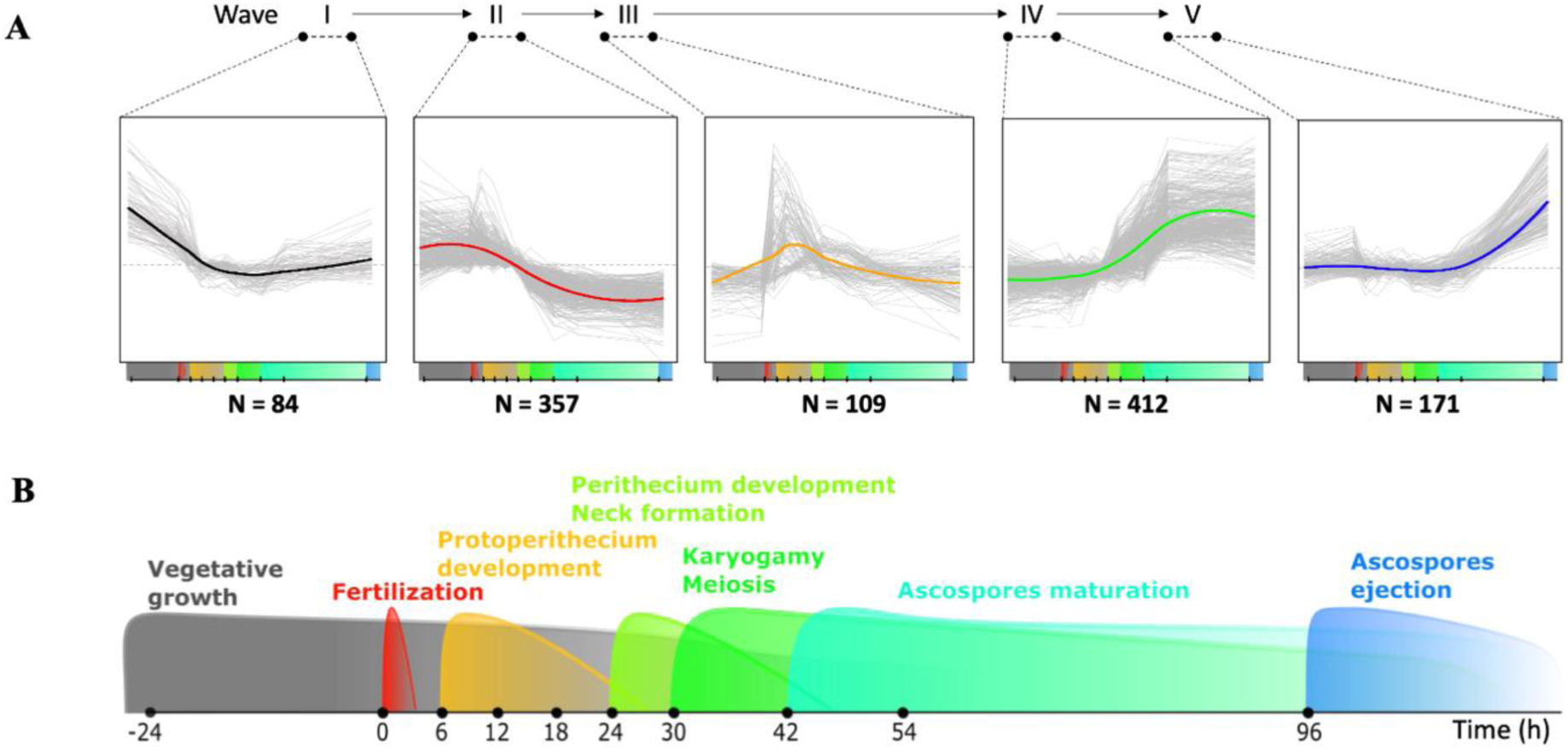
Temporal transcriptional waves over the course of sexual development. **A.** Expression profiles of the five major successive transcriptional waves (numbered in Roman figures) that take place during *P. anserina* sexual reproduction (Bottom timeline, see panel B for zoom). Number of differentially expressed (DE) genes present in each wave is indicated below the corresponding graph. Waves II (early up/late down pattern, 31.51%) and IV (early down/late up pattern, 36.37%) accounted for 67.87% of them (33.98% and 37.42%, respectively), while wave I (early down pattern, 7.41%), wave III (early-mid up pattern, 9.62%) and wave V (late up pattern, 15.02%) contained only 32.13% of them, respectively. **B.** Major developmental events according to the sexual reproduction timeline described in Figure 1.

The topology of the transcriptional waves therefore suggests that the genes of waves II and III are down-regulated after a peak of expression during protoperithecium development, while the genes of waves IV and V are up-regulated after perithecium development. These patterns are in accordance with the results obtained in *N. crassa*, during fruiting body development (Wang *et al*. 2014).

### Evolutive conservation and functional annotation of differentially expressed genes during sexual reproduction

Across the entire time-course experiment, we found 3,466 genes that were differentially expressed (DE, see Methods). They represent 33% of the total gene set for which gene expression measurements were monitored (Table S1, Fig. 4A), including 1,133 of the genes clustered in the five transcriptional waves (more than 95% of the clustered genes, Table S1 and Fig. 3 and Fig. S1). Overall, this analysis revealed that in *P. anserina*, one-third of the complete gene set is transcriptionally regulated during sexual reproduction. Of this, one-third follows one of the five transcriptional waves disclosed by the clustering analysis. To test for evolutive conservation of these DE genes, we built group of orthologs (see Methods), using four species of filamentous fungi, *i.e*. the homothallic species *Chaetomium globosum* and *Sordaria macrospora* and the heterothallic species *N. crassa* and *Trichoderma reesei*. A total of 2,957 DE genes (more than 85%) were identified as belonging to an ortholog group shared with at least one of the four species (Table S3). A core set of 1,496 DE genes is conserved in all four fungi (Fig. 4, Table S3).

**Figure 4 -.**
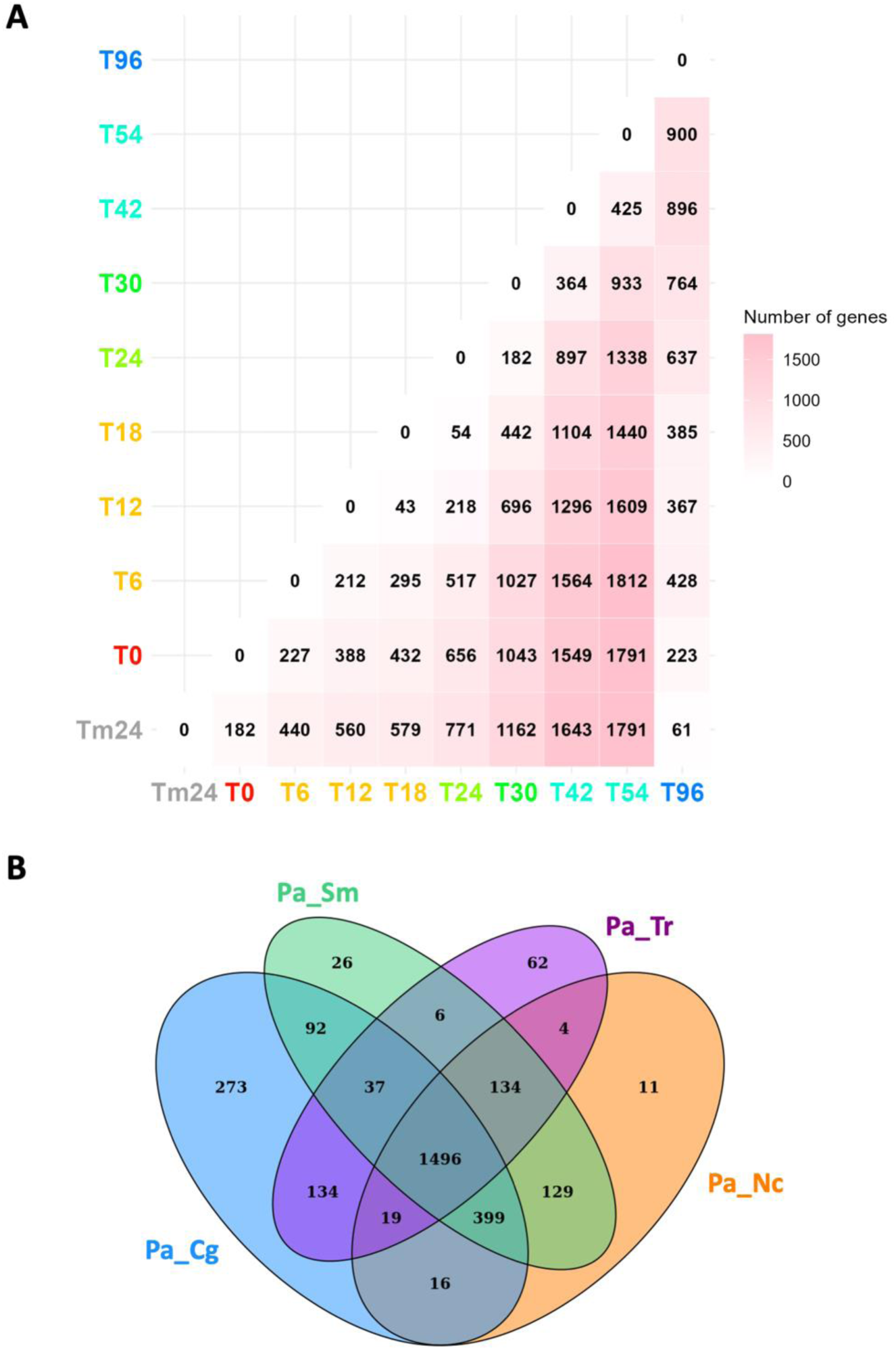
Distribution of differentially expressed genes along the kinetics of sexual reproduction and their evolutive conservation. **A.** Heatmap of pairwise comparisons of differentially expressed (DE) genes across the complete time-course experiment. Numbers correspond to the numbers of genes with an adjusted p-value lower than 0.001 and an absolute fold change value between the two compared time points, higher than 2 (see Methods). **B.** Venn diagram showing the conserved orthologs between the indicated fungal species. Cg: *C. globosum* (homothallic), Sm: *S. macrospora* (homothallic), Nc: *N. crassa* (heterothallic) and Tr: *T. reesei* (heterothallic).

A pairwise comparison of the DE gene set of *P. anserina* with each gene set of the four fungi, showed that the number of conserved DE genes is consistent with fungal evolutionary history. Closely related species (*C. globosum*) share more DE genes than the more distantly related species (*N. crassa*, *S. macrospora* and *T. reesei*). The remaining 506 genes (14.78%) were classified as orphan genes, with 247 (21.84%) belonging to the five transcriptional waves. Notably, the set of DE genes, including that of the five waves, were found to be enriched with orphan genes (p-values: 2.55×10^−9^ and 4.60×10^−20^ respectively).

We then search for functional annotation of the proteins encoded by the DE genes (see Methods). We found that 1,279 (36.95%) of the 3,466 putative proteins encoded by the DE genes lacked Pfam annotation (Table S1, (Mistry *et al*. 2021)), which is significantly more (p-value: 8.55×10^−9^) than in the complete list of *P. anserina*’s proteins. The lack of annotated conserved domains was further enriched (p-value: 1.48×10^−13^) in wave-embedded DE genes, as 515 out of these 1,133 putative proteins (45.44%) lacked Pfam annotation. We also performed Gene Ontology (GO, (Ashburner *et al*. 2000; The Gene Ontology Consortium *et al*. 2023)) analysis on the DE genes (Table S4, Fig. 5A).

**Figure 5 -.**
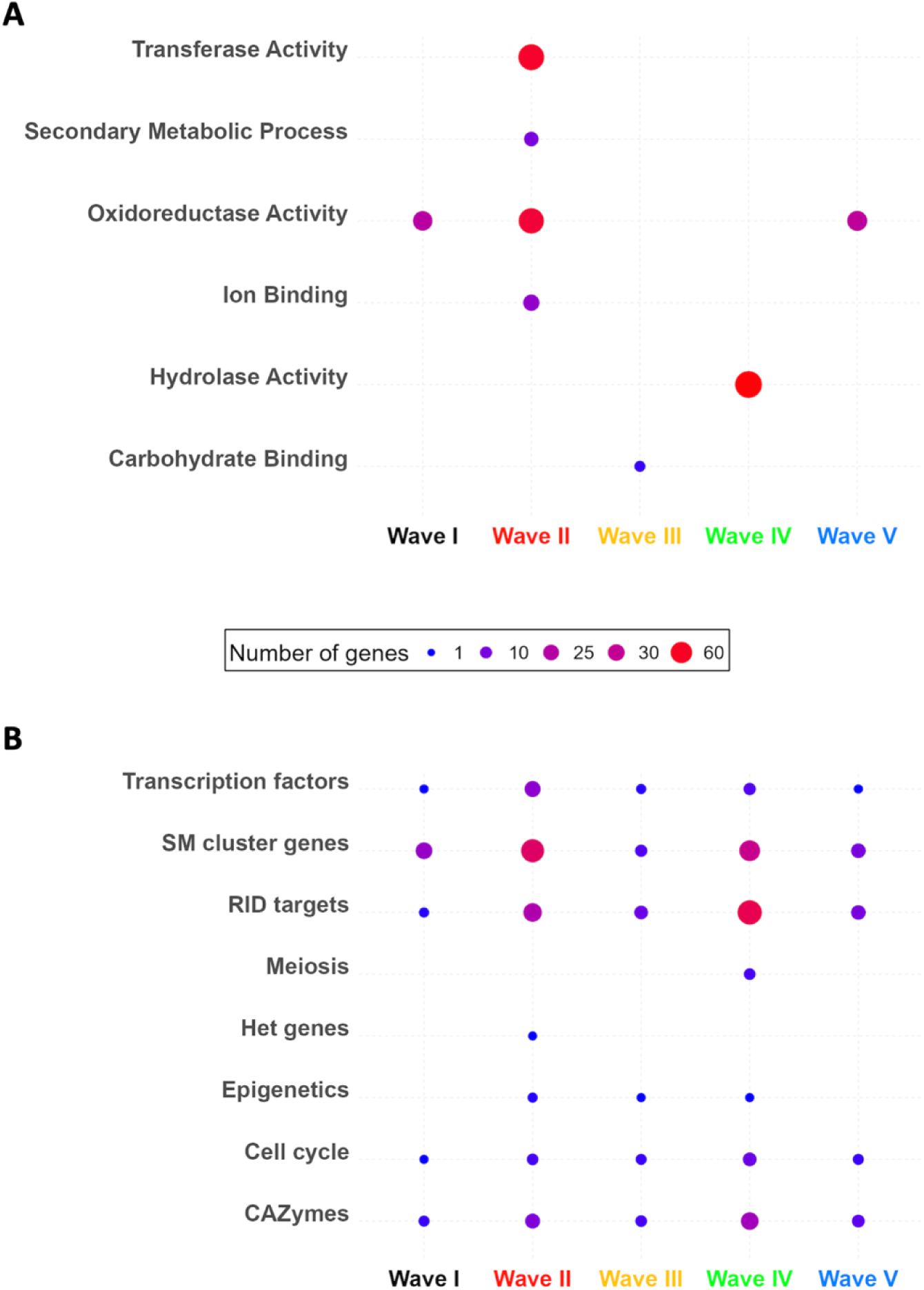
GO term and expert annotation categories enriched in the transcriptional waves. **A.** Descriptions of GO terms significantly enriched (p-value ≤ 0.05) in at least one wave of the transcriptional waves. **B.** Enrichments of genes of interest in at least one wave of the transcriptional waves. TF: transcription factors, SM: secondary metabolites, HET: fungal specific vegetative incompatibility HET domain. Points are scaled and colored according to the number of genes in each category for each wave. No score is shown for zero genes.

In general, the enriched GO terms and Pfam domains (Table S4 and S5) described intense catalytic (GO:0003824) and metabolic (GO:0008152) activities, associated with binding (GO:0005488, PF00187). Accordingly, we observed enrichments in genes involved in redox catalytic activities (GO:0003824, PF00067, PF03055, PF01565, PF00106, PF01494, PF00199, PF08240) and transporter activities (GO:0005215, Pfam PF07690.19). We also uncovered enrichments in DE genes involved in cellular differentiation programs more specifically related to sexual reproduction. They either deal with cellular anatomical entity (GO:0110165), localization (GO:0051179) or encode zinc-binding domain transcription factors (PF13695), necrosis inducing factor (PF14856) and TUDOR domain containing protein (PF11160). However, if the repertoire of GO terms (Table S4) and Pfam domains (Table S1 and S5) generated by this study should provide some clues to select candidate genes for *in vivo* functional characterization, our analysis essentially showed that more than one third of the DE genes have no associated function yet.

### Searching for genes of interest, the expertise-driven approach

We then performed a biological expertise-driven approach to mine the functions of the full set of DE genes (Table S6, Fig. 5B).

#### Developmental processes at work

Fertilization of *P. anserina* requires exposure to blue light, but the molecular basis of this signal transduction is still unknown. In *N. crassa*, the blue light signal is transduced by the photoreceptors WC-1 and WC-2 (Froehlich *et al*. 2002; He *et al*. 2002). In this study, we found five blue light responsive genes that were differentially expressed (Table S6A, Fig. S5A-B). Consistent with a role in fertilization three of them (including *Pa_ 5_6130/WC-1*) were expressed before meiosis, while the other two were expressed after meiosis, suggesting that blue light may also play a role during ascospore formation.

Orthologs of genes involved in plasmogamy in yeast models (*Pa_3_11320/Prm1* (Curto *et al*. 2014), *Pa_3_4770/CDC3* (Kurahashi, Imai, and Yamamoto 2002), *Pa_5_9510/Fig1* (Erdman *et al*. 1998), *Pa_1_1930/KEX2* (Julius *et al*. 1984), *Pa_4_6420/KAR2* (Rose, Misra, and Vogel 1989), *Pa_7_10810/SEC63* (Young *et al*. 2001) were found to be up-regulated until T42 and then downregulated (Table S6A, Fig. S5C). Development of the zygotic lineage likely involves compartmentalization through active cell wall remodeling, as we found several DE genes encoding different types of cell wall proteins and growth factor receptors (Table S6A). One of them was the expansin-like EEL2, involved in cell wall expansion, whose gene *Pa_2_310* was downregulated from karyogamy to ascospore maturation (T30-T96). This expression pattern was similar to that of the *C. globosum* homolog (*CHGG_00523*) but divergent from that of the *N. crassa* homolog (*NCU04603*) (Hutchison and Glass 2010; Wang *et al*. 2019). Cell-cell interactions may also be tightly regulated, as exemplified by the strict downregulation (T0-T96) of *Pa_1_17480,* which encodes the adhesin protein MAD1 (Wang and St Leger 2007).

Since sexual development is rich in cell divisions, we searched for DE cell cycle related genes and found this category slightly enriched (p-value = 2.9×10^−2^, Table S5B). In addition to the TOR pathway inhibitor *Pa_3_3700/FgFkbp12*, it included homologs of genes required for proper fructification formation (*Pa_1_17170/KIN3*, *Pa_1_23300/cel-2*, *Pa_2_6460/Pah1*, *Pa_2_890/asm-1*), axial budding (*Pa_2_290*/*axl2*) and genes encoding G-coupled receptors (*Pa_4_5350*, *Pa_1_590*) and ras-like protein (*Pa_1_19030*/*krev-1*). The expression profile of the *Pa_2_290*/*axl2* gene, encoding an integral plasma membrane protein, was of particular interest: it was down-regulated when the two haploid parental nuclei actively divide into multinucleated syncytial ascogonial cells (Fig. S5D).

As expected, we also found DE genes encoding cytoskeletal elements (Fig. S5E). Interestingly, *Pa_5_5720*, encoding a pyruvate decarboxylase (PDC) was found strongly up-regulated during ascospore maturation (wave V). This observation further enforces the hypothesis that pyruvate decarboxylase is involved as a structural protein (PDC-filaments) associated with the cytoskeleton during the sexual development of filamentous fungi (Thompson-Coffe *et al*. 1999). Conversely, *Pa_4_7260*, encoding the nucleoprotein Cik1, required for both karyogamy and mitotic spindle organization (Cottingham *et al*. 1999), was found down-regulated at the same time. Some of the cytoskeleton DE genes showed co-expression with the plasmogamy DE genes (Fig. S5F), making them good candidates to be involved in fusion morphogenesis processes.

Three genes essential for ascus development were also found DE (Fig. S5G, Table S6A): *P. anserina*’s orthologs of *Pa_4_9730/asd-1* (wave IV (Nelson, Merino, and Metzenberg 1997)), *Pa_4_9750/asd-3* (Galagan *et al*. 2003) and *Pa_2_13270/con-6* (wave V (White and Yanofsky 1993)). *P. anserina* does not form clonal propagules. However, the orthologs of genes required for conidiation in *N. crassa* and *Aspergillus nidulans* such as *Pa_2_13270/con-6*, *Pa_4_9730/asd-1*, *Pa_4_9750/asd-3, Pa_1_12720/Pyg1*, *Pa_1_30/TmpA* and *Pa_0_1540* were found to be up-regulated after meiosis, suggesting that genes primarily involved in conidiation, *i.e.* asexual reproduction, may have been recruited to participate in ascospore formation. On the contrary, early conidial development-2 *Pa_7_780/ecd-2* was down regulated during the first half of the kinetic (Sun *et al*. 2012).

#### Producing energy

The reproductive process is energy consuming and therefore the mobilization of carbon resources is one of the keys to its success. The genome of *P. anserina* contains 276 genes encoding CAZymes ((Espagne *et al*. 2008) and Table S6D). Among them, 141 were found DE during sexual development, showing an enrichment of this metabolic activity (p-value = 4.36×10^−5^). The expression profile of 42 CAZyme genes corresponded to one of the five transcriptional waves, especially waves II and IV. The remaining 99 showed contrasting expression patterns (Fig. S5 H-T), but up-regulation was enriched over down-regulation, as 62 of them were up-regulated during the kinetic (Chi², p-val < 0.001). We also found that several CAZymes genes had some of the highest positive fold changes measured in this study (Table S1). In addition, three glycolysis enzymes genes (6-phosphofructo-2-kinase *Pa_3_6610*, fructose-bisphosphate aldolase *Pa_0_710*, and phosphoglycerate kinase *Pa_6_10610*) were up-regulated with a similar pattern of overexpression when asci begin to differentiate (Fig. S5U, Table S6E).

In *P. anserina*, long-chain fatty acids are shortened in peroxisomes then degraded in mitochondria. The *Pa_6_3500*/*PaPex7* gene, which encodes the peroxisomal-targeting-signal-2 receptor, is found to be up-regulated during ascospore maturation (T54-T96, Table S1 and Fig. S5V). This is in line with the defects in nuclear positioning of the post-meiotic asci observed in *pex7* knock-out mutants, which can result in either abnormal ascospore formation or to ascospores containing an abnormal number of nuclei (Bonnet *et al*. 2006). The *Pa_1_23790*/*PaFox2* gene (Table S1 and Fig. S5V), which encodes the peroxisomal multifunctional protein 2, a key enzyme in the fatty acid ß-oxidation pathway, was found to be down-regulated during the early stages of sexual development (T12-T54) and up-regulated during ascospore maturation (T54-T96). This expression profile may explain the reduced acetyl-CoA availability within the ascospores that results in the pigmentation defect observed in *PaFox2* knock-out mutants (Boisnard *et al*. 2009).

#### Secondary metabolites: major actors of sexual development?

Prokaryotic and eukaryotic micro-organisms produce a wide variety of secondary metabolites (SMs). These molecules are not intended to perform housekeeping functions, but rather are used as adaptive tools to cope with environmental fluctuations or to compete with other living organisms. The *P. anserina* genome contains 35 biosynthetic gene clusters (BGCs) (Lamacchia *et al*. 2016), Table S6F, Fig. S5W), containing 474 genes. SM encoding genes were enriched among the complete set of DE genes (p-value = 8.31×10^−10^) as 258 of them were found to be DE (33 out of 35 BGCs contained at least two DE genes). These genes were also significantly up-regulated (Chi^2^, p-val < 0.001). BGCs N°1, 11, 18, 19, 24, 27 and 34 showed an expression profile topology (Fig. S5W) similar to the general pattern described in *N. crassa* (Wang *et al*. 2022), which was a coordinated downregulation from fertilization to meiosis followed by a coordinated upregulation during ascospore formation and maturation.

We also examined whether co-expression was shared by multiple SM genes within the 34 BGCs. This was the case for the sterigmatocystin BGC, where 17 of the 23 genes were differentially expressed during sexual reproduction (Table S6F, cluster 32, Fig. S5X, (Slot and Rokas 2011)), including the transcription factor *PaAflr* (Shen *et al*. 2019). In addition, we identified four positive regulators of the sterigmatocystin BGC (Table S6F, Fig. S5Y, *Pa_2_7220, Pa_2_7240, Pa_2_7250, Pa_2_7260* (Bhetariya *et al*. 2011)), that showed the same expression pattern across sexual reproduction. Similarly, 13 of the 20 genes in the terrequinone BGC were differentially expressed (Fig. S5Z).

In addition, we identified several isolated genes from different BGCs that showed similar peak expression to each other but no co-expression with other genes from their respective BCG (Fig. S5AA-AB). These expression peaks of genes encoding mostly proteins of unknown function corresponded to two key moments in early development (T6 and T12), just before and during dikaryon formation. Taken together, our analyses further support the key role of SM in sexual reproduction and highlight the role of previously uncharacterized clusters and isolated SM genes.

#### From regulation of biotic interactions to that of sexual development

Genes encoding STAND proteins (Signal-transducing ATPases with Numerous Domains, (Leipe, Koonin, and Aravind 2004) have been described as involved in fungal biotic interactions with living organisms, either from the same species or from different species but mostly during vegetative growth. The *P. anserina* genome contains 34 genes encoding STAND proteins with a NACHT domain (PF05729), eight of which were found to be down-regulated either throughout the sexual cycle kinetics or prior to perithecium development (Table S6G, Fig. S5AC). This group includes *Pa_3_10930*/*nwd1* (Chevanne *et al*. 2010), *Pa_7_10370*/*sesB-like*, *Pa_2_7940*/*het-d* (Espagne *et al*. 2002) and *Pa_1_11380* (Bidard, Clavé, and Saupe 2013). Fungal interactions are also under the control of a programmed cell death called vegetative incompatibility that prevents hyphal fusion of two genetically distinct wild-type strains. Most of the effectors of this incompatibility reaction (IR) contain HET domains (PF06985). Strikingly, 59 of the 128 *het* genes annotated in the *P. anserina*’s genome were found to be DE, including *Pa_4_1190*/*hnwd1* and *Pa_7_3460*/*Pin-C* (Chevanne *et al*. 2010; Kaneko *et al*. 2006; Bidard, Clavé, and Saupe 2013). The majority of the DE *het* genes were down-regulated either throughout the complete sexual development or from dikaryon formation (Fig. S5AD), but seven of them were up-regulated during some steps of the kinetic, mostly after karyogamy (Fig. S5AE). In addition, three *het* genes showed a peculiar behavior (Fig. S5AF): *Pa_5_2540* expression oscillated during sexual reproduction, *Pa_3_5550* expression peaked during protoperithecium development and *Pa_4_1800* expression was down-regulated until ascospore maturation and over-expressed during this process.

In the DE gene set, we also found gene families encoding toxins that can induced host cell death or effector-like proteins. All of the six genes encoding the NEP1-like proteins pore-forming toxin NPP1 (necrosis-inducing *Phytophthora* protein, (Ottmann *et al*. 2009) were down-regulated at least during the second half of the sexual development, i.e. before karyogamy (Fig. S5AG). Six out of the eight genes encoding Hce2 pathogen effectors (Stergiopoulos *et al*. 2013) and one gene annotated as Ptu bug toxin encoding gene (*Pa_6_9450*) were also DE. In contrast to the *NPP1* genes, all of them showed up-regulation during the course of sexual development, five of them belonging to the wave IV pattern of expression (Fig S5AH).

#### Epigenetic effectors at work in sexual reproduction

In recent years, epigenetic effectors have been identified as key players in fructification morphogenesis (Carlier *et al*. 2021; Nowrousian 2022). Therefore, we investigated the expression patterns of genes encoding proteins that can either methylate cytosines, write or erase histone post-translational modifications or interfere with messenger RNA synthesis and stability.

*PaRid* and *PaDnmt5*, two of the three DNA methyltransferase encoding genes present in the *P. anserina* genome, were found to be DE during this time-course experiment. *PaRid* expression, which is essential for dikaryon formation (Grognet *et al*. 2019) peaked at T12, while *PaDnmt5* expression was increasingly up-regulated from T12 to T42 (Fig. S5AI). *Pa_5_2900*, which encodes a RING-type E3 ubiquitin transferase URF1-like, was found to be strongly up-regulated from T42. In mammals, this protein binds both hemi-methylated DNA and tri-methylated lysine 9 of histone H3 (H3K9me3), thereby bridging DNA methylation and chromatin modification (Foster *et al*. 2018).

*Pa_7_10020*/*KAT1* and *Pa_1_7460*/*Hos-2* encodes two chromatin modifiers with antagonistic enzymatic activities, respectively a histone acetyltransferase encoding gene (Tong *et al*. 2012) and a histone deacetylase (Wang, Kurdistani, and Grunstein 2002) respectively. However, these two genes showed co-expression over the entire kinetic, *i.e*. they were up-regulated from fertilization to karyogamy and then down-regulated (Fig. S5AJ). This pattern was in contrast to that of *Pa_7_5890* encoding the Sin3-associated polypeptide SAP18, which enhances transcriptional repression through histone deacetylation (Cheng and Bishop 2002). Two genes encoding sirtuins NAD(+)-dependent histone deacetylases were also found to be DE: *Pa_4_8570*/*PaSir2* showed a wave III expression pattern that resembled that of *Pa_1_19440*/*PaRid* (Fig. S5AI), while the gene encoding *Pa_4_7390*/*SIRT5* was down-regulated throughout sexual reproduction. Of the 32 genes encoding SET domain proteins, seven were found to be DE (Fig. S5AK), although none has yet been shown to have histone lysine methyltransferase activity. Four genes encoding putative histone demethylase containing JmJC domains were also found DE (Fig. S5AL).

Promoter accessibility also depends on nucleosome positioning. The ATP-dependent chromatin remodeling protein SWI/SNF *Pa_5_4310/SMARCA3* (Oh *et al*. 2013) was one of the most highly up-regulated wave III genes (Fig. S5AM). Its transcription pattern is similar to that of *Pa_1_19440*/*PaRid* (see Fig. S5AI) and *Pa_4_8570*/*PaSir2* (Fig. S5AJ). Transcription requires a large number of proteins assembled into conserved complexes. Two genes (*Pa_1_17860* and *Pa_6_1460*) encoding components of the SAGA complex were found to be up-regulated from T6 to T54 with very similar expression patterns (Fig. S5AM). However, we also found similarly co-regulated genes although encoding factors with antagonistic effects on transcription, such as the SWD2 subunit of the COMPASS complex (*Pa_7_11070*) and the extra sexcombs subunit of the Polycomb Repressive Complex 2 (*Pa_3_4080*).

There is no experimental evidence that post-transcriptional RNAi mechanisms Quelling (Romano and Macino 1992) and MSUD (Shiu *et al*. 2001) are functional in *P. anserina*. However, genes encoding essential components of Quelling (Fig. S5AN), the RNA-dependant RNA polymerase (RdRP) *Pa_7_4790/Qde1* (Cogoni and Macino 1999) and the dicer *Pa_6_6150/Dcl-1* (Yang, Ye, and Liu 2015), were downregulated from T18 and T30, respectively. On the contrary, expression of *Pa_7_9210* encoding the Qde-2 interacting exonuclease QIP was up-regulated from T6 to T30. *N. crassa* Δ*qde-1* quelling mutant strains produced normal-sized perithecia but no ascospore (Wang *et al*. 2014). Four genes involved in MSUD were also found to be DE (Fig. S5AO). *Pa_2_9830*/*Sad-2* and *Pa_5_11380*/*Sad-5* were up-regulated whereas *Pa_1_19900*/*Sad-1*, encoding the RdRP and *Pa_1_16040*/*Sad-3*, encoding the helicase, were down-regulated. Interestingly, *Pa_6_11690*/*rrp-3*, which encodes an RdRP that could not be clearly assigned to Quelling or MSUD pathways in *N. crassa* (Borkovich *et al*. 2004), was co-expressed with the Quelling genes *Pa_1_19900*/*Sad-1* and *Pa_1_16040*/*Sad-3* in *P. anserina*. Knockout of the *N. crassa Sad-1* and *Sad-3* resulted in arrest of ascus development in meiotic prophase and production of normal-sized perithecia and young asci without ascospores, respectively (Wang *et al*. 2014). *Sad-2* is essential to *N. crassa* fructification development (Wang *et al*. 2014). These results suggest that RNAi genes may be silenced during meiosis and sporulation, which may explain the apparent lack of MSUD in *P. anserina* but also that *Pa_2_9830*/*Sad-2* and *Pa_5_11380*/*Sad-5* might have been recruited to play a role during ascospore dormancy.

Repeated sequences present in both haploid parental genomes of *P. anserina*’s dikaryotic cells are subjected to cytosine-to-thymine mutations by repeat-induced point mutation (RIP, (Bouhouche *et al*. 2004)). Modified 5-methylcytosines are known to be hotspots for cytosine-to-thymine transitions, because, unlike unmethylated cytosines, spontaneous deamination of 5-methylcytosines cannot be repaired by uracil excision (Pearl 2000). To counteract this genetic threat, members of the AlkB alpha-ketoglutarate-dependent dioxygenase (PF13532) family oxidize 5-methylcytosines to restore undamaged bases (Bian *et al*. 2019). Two genes encoding AlkB-like enzymes are present in the *P. anserina* genome. The expression of *Pa_1_12280* was down-regulated throughout the kinetic, while *Pa_1_17260* was up-regulated during two segments of the kinetic: i) from Tm24 to T12, corresponding to the peak of expression of *PaRid* and ii) from T24 to T54, when *PaRid* is no longer expressed (Fig. S5AP). Notably, both genes showed a superimposable flat expression profile at the time of RIP.

In filamentous fungi, a conserved post-transcriptional A-to-I mRNA editing process recodes transcription units (Teichert *et al*. 2017; Blank-Landeshammer *et al*. 2019), mainly during fructification and ascospore morphogenesis. We identified seven DE genes encoding adenosine deaminase acting on tRNA (ADAT), a class of enzymes which are responsible for A-to-I mRNA editing. *Pa_1_21170*, a wave III gene, showed a peak of expression similar to that of *PaRid*, while *Pa_3_6600* and *Pa_6_7750* were up-regulated from perithecium development to ascospore discharge (Fig. S5AQ).

### Co-expression Network: In Search of Potential Transcription Factor Targets During Sexual Reproduction

We then focused on the 96 TFs encoding genes that were found to be DE (Table S6I). The expression profile of each TF gene was used to search for significant correlations with the expression profiles of all genes previously classified as transcriptional waves I, II, III, IV or V (see Methods). Using this strategy, we build a network of 2002 potential transcriptional links between 76 TFs and 683 target genes (Fig. 6 and Table S7A-C).

**Figure 6 -.**
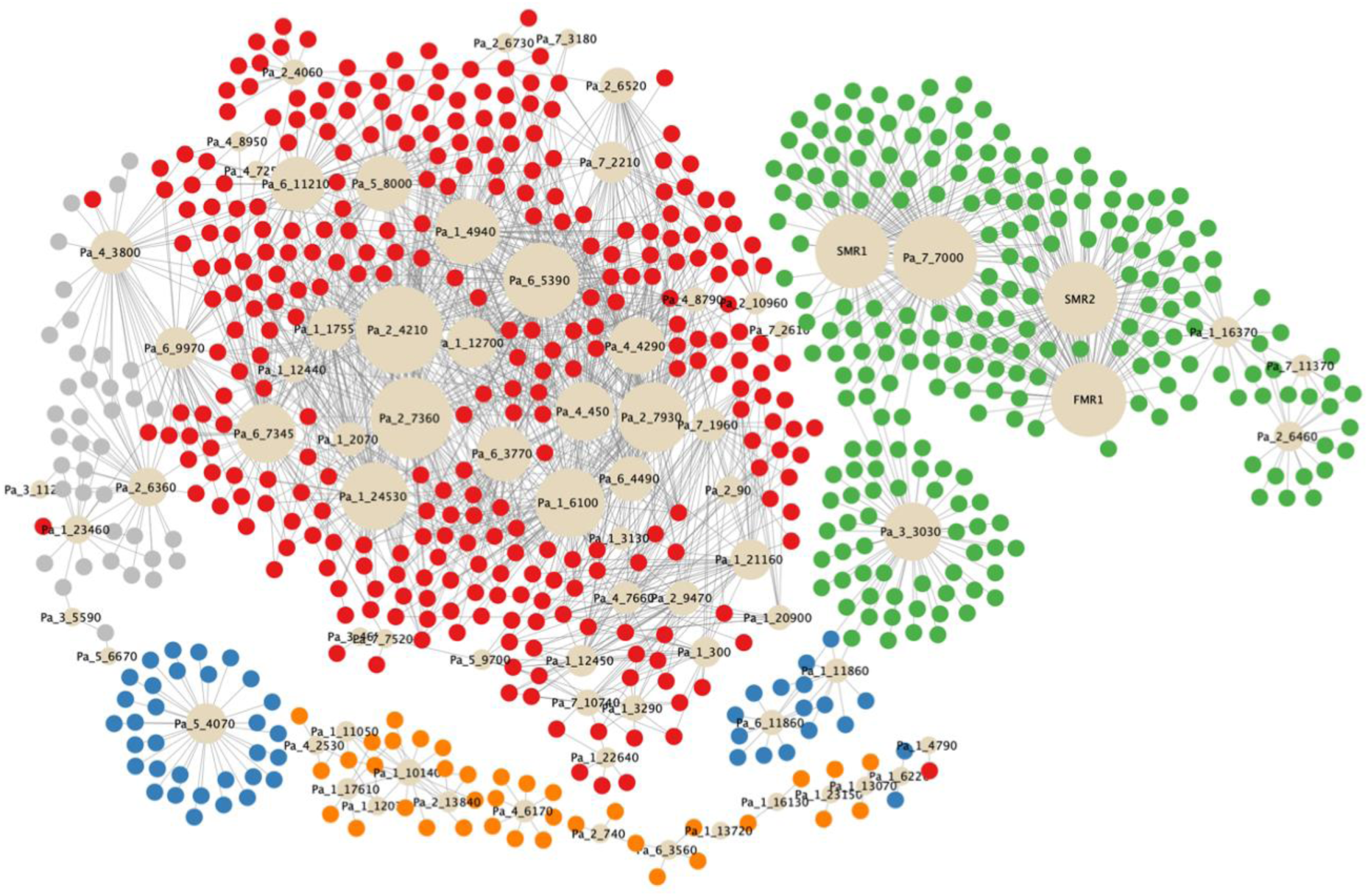
Co-expression network based on expression profiles of differentially expressed (DE) genes coding for transcription factors. Genes encoding transcription factors (TF) are represented by beige circles whose size represents the number of co-regulated genes. These coregulated genes, which can be considered as potential TF target genes, were classified in transcriptional waves as such: wave I: grey color (61 genes), wave II: red color (1402 genes), wave III: orange color (58 genes), wave IV: green color (426 genes) and wave V: blue color (55 genes).

Some of the prominent nodes of this network were expected TFs (FMR1, SMR1, SMR2, Pa_1_10140/Pro1, see below), confirming its relevance. However, most of the 76 TFs (72%) had no characterized biological function, including those with the broader set of target genes. The distribution of the TFs encoding genes in the five transcriptional waves differed from that of the total set of DE genes (Fig. 6), with the majority belonging to wave II. As with TFs, most of the target genes were of unknown function. Importantly, some TFs were also target genes, e.g. *Pa_1_10140/Pro1*is a target of Pa_1_12070/HOM3Pa_1_11050/HMG4, Pa_1_17610, and Pa_2_13840 (Table S7A).

Among the master regulators of this network, we identified HMG-domain and homeodomain TFs (Espagne *et al*. 2008; Ait Benkhali *et al*. 2013). The HMG-domain TFs include the mating-type proteins FMR1 and SMR2, which are both required for the development of the perithecium after fertilization (Zickler *et al*. 1995; Arnaise *et al*. 2001). Physical interaction between FMR1 and SMR2 was demonstrated and proposed to be required for their post-fertilization functions (Arnaise *et al*. 1995; Turgeon and Debuchy 2007; Bidard *et al*. 2011). Our study shows that FMR1 and SMR2 have a similar expression profile throughout sexual reproduction (Fig. S5AR). Their co-expression networks revealed also an almost identical number of potential target genes (FMR1, n = 79; SMR2, n = 78) (Table S7). They share 62 of their 79 potential target genes (Table S7A). These observations support the idea that they act together in identical regulatory pathways by forming a heterodimer (Arnaise, Debuchy, and Picard 1997). The third and last *mat-* mating-type gene, *SMR1*, does not belong to the HMG-box family. It encodes a HPG protein (Turgeon and Debuchy 2007), which has been proposed to provide a scaffold for physical interaction of mating-type proteins (Zheng *et al*. 2013). This role is consistent with its presence in wave IV, along with *FMR1* and *SMR2* (Fig. S5AR). Critical genes for the development of ascogenous hyphae are likely to be found among target genes of FMR1, SMR2 and SMR1 (Fig. 6, Table S7A). The only non-mating-type HMG protein present in the network is Pa_1_11050/PaHMG4, related to Nhp6p and Hmo1p from *S*. *cerevisiae* (Ait Benkhali *et al*. 2013). Its deletion in *P*. *anserina* resulted in an increased number of male organs. The lack of effect on fructification development is consistent with the down-regulation of *Pa_1_11050*/*PaHMG4* at T42-T96 (Fig. S5AS). Proper shaping of the developing fructifications requires the homeobox protein *Pa_2_6460/Pah1* (Coppin *et al*. 2012). The expression of the corresponding gene was accordingly up-regulated from T0 to T42 (Fig. S5AS). In contrast, *Pa_3_4650/Pah6* was down-regulated between T0 and T54 (Fig. S5AS), although no fructification defect was associated with its deletion.

The Zn(II)_2_Cys_6_ class of regulators is one of the largest families found in fungi (MacPherson, Larochelle, and Turcotte 2006) and consequently they represented almost half of the TFs present in the network (Table S7A). Three of them are of particular interest and have been extensively studied because of their properties. *Pa_1_10140*/*PaPro1* is a regulator of the expression of the mating-type and *IDC* genes, both required for female fertility (Gautier *et al*. 2018). In *S. macrospora*, *Pro1* has been identified as a master regulator of fructification formation (Steffens *et al*. 2016; Teichert, Pöggeler, and Nowrousian 2020). Its temporal and spatial expression is consistent with the presence of *Pa_1_10140/PaPro1* in wave III (Fig. S5AS). *Pa_7_7000* was characterized in wave IV as a coregulator of FMR1, SMR2 and SMR1 target genes (Fig. 6). *Pa_7_7000* is revealed as one of the most prominent nodes in the TF network (Fig. 6 and Fig. S5AT), but has not yet been characterized in *P. anserina*. *Pa_7_7000* has orthologs in the seven species of the *Podospora* complex (Boucher, Nguyen, and Silar 2017) and in some fungi closely related to this species complex (*Cercophora samala*, *Triangularia verruculosa* and *Apiosordaria backusii*). However, more distantly related species (*Podospora setosa*, *Podospora australis*, *Podospora appendiculata*, *Thermothelomyces thermophilus*, *Neurospora tetrasperma* and *Neurospora discreta*) contain a protein that shares some identity with Pa_7_7000, but they all lack the Zn(II)2Cys6 domain and are therefore unlikely to be functional orthologs. Wave V contains *Pa_6_11860*/*Fluffy* (Bailey and Ebbole 1998) (Fig. S5AT), which is required in *N. crassa* for the formation of conidia. These vegetative spores are absent in *P. anserina*, suggesting that this gene has been recruited for ascospore formation in this fungus, as in others members of the *P. anserina / P. pauciseta / P. comata* species complex (Boucher, Nguyen, and Silar 2017).

### Comparison of mating-type target genes at different developmental stages

A transcriptomic analysis of *P. anserina* was performed previously to determine the FMR1 and FPR1 target genes required for fertilization (Bidard *et al*. 2011). The analysis focused on the comparison of *mat-* and *mat+* mycelia competent for fertilization, namely the stage preceding the development of the perithecium. It was performed using the same methods as described here. The analysis identified 69 genes upregulated in a *mat*- mycelium and 88 genes upregulated in a *mat*+ mycelium. Among these genes, we search for the DE genes identified in this study. A total of 32 DE genes (46 %) were identified among the 69 genes which were found to be upregulated in the *mat-* mycelium (Table S8) and 45 DE genes (51%) are found among the *mat*+ upregulated genes (Table S9). A statistical analysis indicates that the DE genes are over-represented in both categories, when compared with the 3,466 DE genes in the 10,507 *P. anserina*’s gene models. This overrepresentation was higher for the *mat*+ upregulated genes (Chi^2^, p-value = 0.0003) than for the *mat*- upregulated genes (Chi^2^, p-value = 0.0183). This result is likely due to the use of a *mat+* maternal strain for the fruiting body development presented here and indicates that half of the genes upregulated in fertilization competent mycelium are also up-regulated in maternal tissues during perithecial development.

## Discussion

### Mating-type gene expression

In this study, we analyzed on a genome-wide scale the transcriptional changes associated with sexual development in *P. anserina* over a 10-time points course, from fertilization to ascospore maturation. We found that more than 32% of the *P. anserina*’s genes are differentially expressed during at least one interval covered by this kinetic. Interestingly, more than 36% of the proteins encoded by these genes do not have a Pfam domain, suggesting that a significant proportion of them remain functionally uncharacterized. Overall, the data generated by our study are rich in novel players involved in fungal sexual reproduction and are likely to provide an exploratory framework to analyze the pathways that are required for the differentiation of the 15 different tissues identified in a perithecium (Bistis, Perkins, and Read 2003).

Among these 15 tissues, the ascogenous hyphae plays a central role in the sexual cycle, since it is where karyogamy, meiosis and ascospore formation take place. Its formation starts at T24 (Fig. 1) and requires the expression of the four mating-type genes: *FMR1*, *SMR1*, *SMR2* and *FPR1* (Zickler *et al*. 1995; Arnaise *et al*. 2001). The ratio of ascogenous tissue to the other tissues of the perithecium is low, potentially masking the zygotic expression profile. This detection limit of our experiments can be assessed by searching the transcripts of *FMR1*, *SMR1* and *SMR2*. These three mating-type genes are specific to the *mat-* nuclei, which come from the male organs and are present only in the ascogonial cell and in the ascogenous hyphae. These three mating-type genes are listed in the DE genes (Table S1), indicating unambiguously that transcripts specific to the ascogenous hyphae are detected in our experiments, in spite of its scarcity. Moreover, upregulations of *FMR1* and *SMR2* transcripts are detected as soon as T24, correlating with the initial development of ascogenous hyphae (Table S1). In addition, 11 genes upregulated in *mat-* fertilization competent mycelium are among the DE genes (Table S8), confirming the detection of transcript from *mat-* male nuclei in the female organ. However, no significant upregulation of the *mat-* mating-type transcripts is detected in the early steps of perithecial development, *i.e*. from T0 to the formation of the ascogonial cell at T18 (Fig. 1). Correspondingly, no wave I DE genes are identified in genes that accumulate transcripts in the *mat-* mycelium (Table S8). The earliest DE genes upregulated in *mat-* competent mycelium belong to wave II (n = 4). Wave IV, which corresponds to upregulation of *mat-* mating-type genes and their correlated genes (Fig. 6), includes 412 genes (Fig. 3), among which six belong to genes upregulated in *mat-* mycelium (Table S8). Overall, these data suggests that critical target genes required for the development from fertilization to the formation of ascogonial cells are not included in the DE genes, because their expression level is below the detection threshold of microarrays. In contrast, the expression of genes required for the subsequent developmental stages (differentiation of ascogenous hyphae, meiosis and ascospore formation) are indeed detected in our experiments.

Surprisingly, *FPR1*/*Pa_1_20590*, the *mat+* specific mating type gene, does not belong to the DE genes, although this gene is also required for the formation of ascogenous hyphae (Zickler *et al*. 1995; Arnaise *et al*. 2001). It is likely that any changes in expression of this gene in zygotic tissues is hidden by its transcription in the maternal tissues, which are predominant in the fruiting body. We have however found 45 DE genes that are specifically upregulated in *mat+* mycelium competent for fertilization (Table S9). Among these DE genes, 32 are direct or indirect target genes of FPR1 (Bidard *et al*. 2011). Their detection proved that FPR1 target genes do not escape our method, although FPR1 itself is not present in DE genes. Moreover, detection starts in the early stages of perithecium development as suggested by FPR1 target genes belonging to wave II (n = 9). Later stages are represented by FPR1 target genes belonging to wave IV (n = 3) and V (n = 2). However, our experiments do not allow us to determine unambiguously whether the differential expression of a gene occurs in the maternal or in the zygotic tissues.

The analysis presented above indicates that our data are pertinent to reveal new pathways in the development of meiotic tissues of P. anserina. Fertilization, which initiates the development of meiotic tissues, is the best-known step of this process. Its molecular description has beneficiated from the yeast model and has revealed that pheromone/pheromone receptor interactions and downstream cascade signal were both conserved in Pezizomycotina and Saccharomycotina (Wilson *et al*. 2021). Subsequent steps have been cytologically well described in numerous fungi and appear specific to Pezizomycotina. In contrast to fertilization, the molecular mechanisms underlying these various cell programmes are unknown. These blind spots may be exemplified by two steps, which are critical in Pezizomycotina to produce numerous genetically different asci from a single fertilization event : i) several models were proposed for the internuclear recognition following mitotic divisions after fertilization, but no one brings a decisive understanding of this phenomenon (Turgeon and Debuchy 2007), ii) internuclear recognition is followed by the formation of dikaryotic cells and croziers, but the molecular mechanisms controlling their formation are unknown. The similarity of dikaryotic cells and croziers to the basidiomycete clamp cells, which require homeodomain transcription factors for their formation, might suggest similar underlying regulatory circuits, but any functions of homeoboxes in dikaryotic cell and crozier formation were dismissed (Coppin *et al*. 2012). A third instance of blind spot is the differentiation of croziers into meiocytes. Although the role of peroxisomes in this event is well documented, the signal triggering the change in cellular programme is unknown (Peraza Reyes and Berteaux-Lecellier 2013). Our analyses revealed co-expression networks, which provide candidates for regulatory circuits of perithecium development and may be involved in the processes presented above. Some particular pathways are discussed below.

### Secondary metabolite gene are actively regulated during sexual development

As in *N. crassa* (Wang *et al*. 2022), *C. globosum* (Wang *et al*. 2019) and *A. nidulans* (Liu *et al*. 2021), our results further support the active role of SMs in fungal sexual reproduction. The evolutionary conservation of their expression profile, *i.e*. downregulation from fertilization to meiosis followed by a coordinated upregulation during ascospore formation and maturation suggests that SMs protect fungal progeny, *i.e*. ascospores, from animal predation. However, we identified several genes of unknown function that were overexpressed during early phases. It is therefore possible that these metabolites protect newly-formed fructification but also that they are recruited to mediate developmental processes. This is reminiscent of the *Pa_5_2090/pks-6* gene, which encodes the insecticidal polyketide neurosporin A produced at a similar developmental stage by *S. macrospora* (Nowrousian 2009) and *N. crassa* (Zhao *et al*. 2017) to protect fructifications from arthropod attack. Whether this gene product is also involved in the process of morphogenesis is still an open question (Wang *et al*. 2022).

Our data reveals a relationship between sterigmatocystin synthesis and mating-type genes by correlating expression of *PaAflR/Pa_2_7360*, *HMG5/Pa_1_13940* and *HMG8/Pa_6_4110* (Table 6 and Fig. 6). PaAflR/Pa_2_7360 is a regulator of the sterigmatocystin synthesis cluster (Shen *et al*. 2019), while HMG5/Pa_1_13940 and HMG8/Pa_6_4110 are HMG-domain TFs acting as hubs of regulation for the mating-type genes and mating-type target genes (Ait Benkhali *et al*. 2013; Gautier *et al*. 2018). Our results suggest that PaAflR/Pa_2_7360 controls positively the transcription of the two other genes. Over expression of PaAflR/Pa_2_7360 results in sterility, as does the over-expression of HMG5/Pa_1_13940 (Shen *et al*. 2019; Ait Benkhali *et al*. 2013). This result is in agreement with the positive transcriptional control of *HMG5/Pa_1_13940* by PaAflR/Pa_2_7360. To date, *PaAflR*/*Pa_2_7360* has not been deleted in *P. anserina*, and although its orthologue in *A. nidulans* has been knocked out, the effect on the sexual cycle remains poorly understood (Wang *et al*. 2022).

### Het genes operates important but yet unknown roles in sexual reproduction

This study unveiled an interesting regulation of the *het* genes, which are involved in allorecognition and thus control the viability of heterokaryons formed after hyphal fusion during vegetative growth (Daskalov 2023). However, heterokaryotic syncytia are not restricted to vegetative thalli, as fertilization also leads to the formation of such multinucleate cells, and accordingly, several *het* loci have been shown to control hybrid fertility in *P. anserina* (Ament-Velásquez *et al*. 2020). When we compared the *het* genes expression profiles during sexual reproduction with those assayed during IR (Bidard, Clavé, and Saupe 2013) and bacterial defense response (Lamacchia *et al*. 2016) (Table S6 G), we found that they fall into two categories and four classes.

The first category consists of het genes that were always down-regulated during sexual reproduction but were either up-regulated during IR and/or bacterial confrontation (class 1) or down-regulated during IR and up-regulated during bacterial confrontation (class 4). This category therefore contains *het* genes that play a role only in vegetative growth and/or defense against pathogens, but must otherwise be silenced.

The second category consisted of *het* genes that were either up-regulated during IR but also at some time points during sexual reproduction (class 2) or were not DE during IR and bacterial confrontation but during sexual reproduction (class 3). Class 2 genes may be recruited to control cell fusion during morphogenesis of fructifications and/or of early zygotic lineage for wave II genes, or of the developing asci and ascospores for waves IV and V genes. Class 3 genes may have been recruited to carry out specific programmed cell death events, not related to hyphal fusion but necessary for sexual morphogenesis. In this context, *Pa_3_5550* is of particular interest since it shows a peak of expression during dikaryon formation. Hce2 proteins also belong to this category, suggesting a specific role during both the pre-zygotic (*Pa_2_4765*) and post-zygotic (*Pa_1_17065*) stages.

The *het-6* and *het-13* genes were found to be DE during sexual reproduction in the heterothallic fungus *N. crassa*, whereas no such observation was made in the homothallic fungus *C. globosum* (Wang *et al*. 2019). In *P. anserina*, both orthologs *Pa_6_6160/het-6* and *Pa_0_150/het-13* were also DE but showed opposite expression patterns compared to *N. crassa*. This suggests that the roles of *het* genes, although central in the sexual developmental network of fungi endowed with two different mating types, are polymorphic.

### Rewiring genes in the RNAi pathway?

In *P. anserina* (Grognet *et al*. 2019), as in *T. reesei* (Li, Chen, and Wang 2018) and *A. immersus* (Malagnac *et al*. 1997), the *rid* (*RIP defective*) ortholog is required for proper sexual reproduction, which is not the case in *N. crassa*. *Pa_1_19440*/*PaRid* showed a short peak of expression at T12 (Fig. S5AI), which is consistent with its requirement for the individualization of the dikaryotic cells leading to meiocytes (Grognet *et al*. 2019). In contrast, *N. crassa rid* up-regulation started at dikaryon formation and increased continuously throughout the second half of sexual reproduction (Wang *et al*. 2014). Interestingly, the *N. crassa rid* expression pattern was similar to that of the *P. anserina* DNA methyltransferase encoding gene *PaDnmt5*, which is absent in *N. crassa*. These results were in agreement with our previous data (Grognet *et al*. 2019) and strengthened the body of evidence pointing to a role for DNA methylation during sexual development.

Although there is no experimental proof that either somatic or meiotic RNAi pathways (Quelling or MSUD, respectively) works efficiently in *P. anserina*, some of the corresponding genes are found to be DE during sexual reproduction. Globally the expression patterns of *P. anserina* genes involved in the RNAi pathways are opposite to those of *N. crassa*, except for *Pa_2_9830/Sad-2* and *Pa_5_11380/Sad-5*. During MSUD, Sad-2 is thought to interact with the RdRP Sad-1 as a scaffold. Yet, *Sad-1* and *Sad-2* showed opposite patterns of expression during *N. crassa* sexual development, which likely discards this hypothesis (Wang *et al*. 2014). However, *Sad-5*, which role is yet unknown, has an expression profile that is similar to that of *Sad-2*. The evolutive conservation of *Pa_2_9830/Sad-2* and *Pa_5_11380/Sad-5* profiles of expression in both *N. crassa* and *P. anserina*, suggests that they have been recruited together to play a role in sexual reproduction. In the homothallic ascomycete *S. macrospora*, *Dcl1*, *Dcl2*, *Sms2* and *Qde2* are involved in meiocyte formation (Girard *et al*. 2021), which is not consistent with the expression pattern of the *Pa_6_6150/Dcl-1* but rather with that of *Pa_5_8910/Sms-2*. Altogether, these results suggest a complex role for the RNAi machinery during fungal sexual reproduction.

## Conclusion

Our work reveals 3,466 DE genes during the differentiation of fruiting bodies of *P. anserina*. Only 86 of these DE genes have been previously functionally analyzed (Table S10), leaving a large number of genes for future investigations. Although our analysis does not allow us to determine the DE genes specifically expressed in each tissue of the fruiting body, our method is sensitive enough to detect DE genes in ascogenous hyphae, at a stage where this tissue is represented by few cells in the mass of the fruiting body. The sensitivity of our method is emphasized by the detection of a peak of transcription of *Pa_1_19440*/*PaRid* at a time of the early perithecial development corresponding to its expected function. Among the DE genes, we have identified at least two pathways that may control mating-type genes. The first one involves *Pa_7_7000*, possibly revealing a new regulatory pathway upstream of mating-type genes. The second one is the relationship between *PaAflR*/*Pa_2_7360*, *HMG5*/*Pa_1_13940* and *HMG8*/*Pa_6_4110*. The latter two TFs control mating-type genes, thus revealing an additional putative pathway regulating indirectly these genes. Our analysis reveals also the intriguing expression patterns of the genes from the MSUD and RNAi pathways, although MSUD has not been found in *P. anserina* and RNAi not investigated. Further research should be focused in *P. anserina* on the role of the genes involved in these pathways in other species.

## Acknowledgements

We are grateful to S. Saupe for the exchange of data and ideas.

## Data, scripts, code, and supplementary information availability

Data are available online: Go to https://www.ncbi.nlm.nih.gov/geo/query/acc.cgi?acc=GSE93094. Enter token whcrwemexfkfhqr into the box.

## References

Ait Benkhali, Jinane, Evelyne Coppin, Sylvain Brun, Leonardo Peraza-Reyes, Tom Martin, Christina Dixelius, Noureddine Lazar, Herman van Tilbeurgh, and Robert Debuchy. 2013. ‘A Network of HMG-Box Transcription Factors Regulates Sexual Cycle in the Fungus *Podospora anserina*’. PLoS Genetics 9 (7): e1003642. 10.1371/journal.pgen.1003642.

Ament-Velásquez, S. Lorena, Hanna Johannesson, Tatiana Giraud, Robert Debuchy, Sven J. Saupe, Alfons J.M. Debets, Eric Bastiaans, et al. 2020. ‘The Taxonomy of the Model Filamentous Fungus *Podospora anserina*’. MycoKeys 75 (November):51–69. 10.3897/mycokeys.75.55968.

Arnaise, S., R. Debuchy, and M. Picard. 1997. ‘What Is a Bona Fide Mating-Type Gene? Internuclear Complementation of Mat Mutants in *Podospora anserina*’. Molecular & General Genetics: MGG 256 (2): 169–78.

Arnaise, S., D. Zickler, S. Le Bilcot, C. Poisier, and R. Debuchy. 2001. ‘Mutations in Mating-Type Genes of the Heterothallic Fungus *Podospora anserina* Lead to Self-Fertility’. Genetics 159 (2): 545–56.

Arnaise, Sylvie, Evelyne Coppin, Robert Debuchy, Denise Zickler, and Marguerite Picard. 1995. ‘Models for Mating Type Gene Functions in *Podospora anserina*’. Fungal Genetics Newsletter, 42:79.

Ashburner, Michael, Catherine A. Ball, Judith A. Blake, David Botstein, Heather Butler, J. Michael Cherry, Allan P. Davis, et al. 2000. ‘Gene Ontology: Tool for the Unification of Biology’. Nature Genetics 25 (1): 25–29. 10.1038/75556.

Bailey, Lori A, and Daniel J Ebbole. 1998. ‘The Fluffy Gene of *Neurospora crassa* Encodes a Gal4p-Type C6 Zinc Cluster Protein Required for Conidial Development’. Genetics 148 (4): 1813–20. 10.1093/genetics/148.4.1813.

Basenko, Evelina Y., Jane A. Pulman, Achchuthan Shanmugasundram, Omar S. Harb, Kathryn Crouch, David Starns, Susanne Warrenfeltz, et al. 2018. ‘FungiDB: An Integrated Bioinformatic Resource for Fungi and Oomycetes’. Journal of Fungi 4 (1): 39. 10.3390/jof4010039.

Bhetariya, Preetida J., Taruna Madan, Seemi Farhat Basir, Anupam Varma, and Sarma P. Usha. 2011. ‘Allergens/Antigens, Toxins and Polyketides of Important Aspergillus Species’. Indian Journal of Clinical Biochemistry 26 (2): 104–19. 10.1007/s12291-011-0131-5.

Bian, K, Sap Lenz, Q Tang, F Chen, R Qi, M Jost, Cl Drennan, Jm Essigmann, Sd Wetmore, and D Li. 2019. ‘DNA Repair Enzymes ALKBH2, ALKBH3, and AlkB Oxidize 5-Methylcytosine to 5-Hydroxymethylcytosine, 5-Formylcytosine and 5-Carboxylcytosine in Vitro’. Nucleic Acids Research 47 (11). 10.1093/nar/gkz395.

Bidard, F, S Imbeaud, N Reymond, O Lespinet, P Silar, C Clave, H Delacroix, V Berteaux-Lecellier, and R Debuchy. 2010. ‘A General Framework for Optimization of Probes for Gene Expression Microarray and Its Application to the Fungus Podospora anserina’ 3 (1): 171.

Bidard, Frédérique, Jinane Aït Benkhali, Evelyne Coppin, Sandrine Imbeaud, Pierre Grognet, Hervé Delacroix, and Robert Debuchy. 2011. ‘Genome-Wide Gene Expression Profiling of Fertilization Competent Mycelium in Opposite Mating Types in the Heterothallic Fungus *Podospora anserina*’. PloS One 6 (6): e21476. 10.1371/journal.pone.0021476.

Bidard, Frédérique, Corinne Clavé, and Sven J. Saupe. 2013. ‘The Transcriptional Response to Nonself in the Fungus *Podospora anserina*’. G3 (Bethesda, Md.) 3 (6): 1015–30. 10.1534/g3.113.006262.

Bistis, George, David Perkins, and Nick Read. 2003. ‘Different Cell Types in *Neurospora crassa*’. Fungal Genetics Reports 50 (1): 17–19. 10.4148/1941-4765.1154.

Blank-Landeshammer, B, I Teichert, R Märker, M Nowrousian, U Kück, and A Sickmann. 2019. ‘Combination of Proteogenomics with Peptide De Novo Sequencing Identifies New Genes and Hidden Posttranscriptional Modifications’. mBio 10 (5). 10.1128/mBio.02367-19.

Boisnard, Stéphanie, Eric Espagne, Denise Zickler, Anne Bourdais, Anne-Laure Riquet, and Véronique Berteaux-Lecellier. 2009. ‘Peroxisomal ABC Transporters and Beta-Oxidation during the Life Cycle of the Filamentous Fungus *Podospora anserina*’. Fungal Genetics and Biology: FG & B 46 (1): 55–66. 10.1016/j.fgb.2008.10.006.

Bonnet, Crystel, Eric Espagne, Denise Zickler, Stéphanie Boisnard, Anne Bourdais, and Véronique Berteaux-Lecellier. 2006. ‘The Peroxisomal Import Proteins PEX2, PEX5 and PEX7 Are Differently Involved in *Podospora anserina* Sexual Cycle’. Molecular Microbiology 62 (1): 157–69. 10.1111/j.1365-2958.2006.05353.x.

Borkovich, Katherine A., Lisa A. Alex, Oded Yarden, Michael Freitag, Gloria E. Turner, Nick D. Read, Stephan Seiler, et al. 2004. ‘Lessons from the Genome Sequence of *Neurospora crassa*: Tracing the Path from Genomic Blueprint to Multicellular Organism’. Microbiology and Molecular Biology Reviews 68 (1): 1–108. 10.1128/mmbr.68.1.1-108.2004.

Boucher, Charlie, Tinh-Suong Nguyen, and Philippe Silar. 2017. ‘Species Delimitation in the *Podospora anserina*/*P. Pauciseta*/*P. Comata* Species Complex (Sordariales)’. *Cryptogamie*, Mycologie 38 (4): 485–506.

Bouhouche, K, D Zickler, R Debuchy, and S Arnaise. 2004. ‘Altering a Gene Involved in Nuclear Distribution Increases the Repeat-Induced Point Mutation Process in the Fungus *Podospora anserina*’. Genetics 167 (1): 151–59.

Boyle, Elizabeth I., Shuai Weng, Jeremy Gollub, Heng Jin, David Botstein, J. Michael Cherry, and Gavin Sherlock. 2004. ‘GO::TermFinder--Open Source Software for Accessing Gene Ontology Information and Finding Significantly Enriched Gene Ontology Terms Associated with a List of Genes’. Bioinformatics (Oxford, England) 20 (18): 3710–15. 10.1093/bioinformatics/bth456.

Breyer, Eva, and Federico Baltar. 2023. ‘The Largely Neglected Ecological Role of Oceanic Pelagic Fungi’. Trends in Ecology & Evolution 38 (9): 870–88. 10.1016/j.tree.2023.05.002.

Carlier, F., M. Li, L. Maroc, R. Debuchy, C. Souaid, D. Noordermeer, P. Grognet, and F. Malagnac. 2021. ‘Loss of EZH2-like or SU(VAR)3-9-like Proteins Causes Simultaneous Perturbations in H3K27 and H3K9 Tri-Methylation and Associated Developmental Defects in the Fungus *Podospora anserina*’. Epigenetics & Chromatin 14 (1): 22. 10.1186/s13072-021-00395-7.

Cheng, Steven Yan, and J. Michael Bishop. 2002. ‘Suppressor of Fused Represses Gli-Mediated Transcription by Recruiting the SAP18-mSin3 Corepressor Complex’. Proceedings of the National Academy of Sciences 99 (8): 5442–47. 10.1073/pnas.082096999.

Chevanne, Damien, Sven J Saupe, Corinne Clavé, and Mathieu Paoletti. 2010. ‘WD-Repeat Instability and Diversification of the *Podospora anserina* Hnwd Non-Self Recognition Gene Family’. BMC Evolutionary Biology 10 (May):134. 10.1186/1471-2148-10-134.

Cogoni, C, and G Macino. 1999. ‘Gene Silencing in *Neurospora crassa* Requires a Protein Homologous to RNA-Dependent RNA Polymerase’. Nature 399 (6732): 166–69. 10.1038/20215.

Coppin, E, and P Silar. 2007. ‘Identification of PaPKS1, a Polyketide Synthase Involved in Melanin Formation and Its Use as a Genetic Tool in *Podospora anserina*’. Mycological Research 111 (Pt 8): 901–8. 10.1016/j.mycres.2007.05.011.

Coppin, Evelyne, Véronique Berteaux-Lecellier, Frédérique Bidard, Sylvain Brun, Gwenaël Ruprich-Robert, Eric Espagne, Jinane Aït-Benkhali, et al. 2012. ‘Systematic Deletion of Homeobox Genes in *Podospora anserina* Uncovers Their Roles in Shaping the Fruiting Body’. PloS One 7 (5): e37488. 10.1371/journal.pone.0037488.

Cottingham, F. R., L. Gheber, D. L. Miller, and M. A. Hoyt. 1999. ‘Novel Roles for *Saccharomyces cerevisiae* Mitotic Spindle Motors’. The Journal of Cell Biology 147 (2): 335–50. 10.1083/jcb.147.2.335.

Curto, M.-Ángeles, Mohammad Reza Sharifmoghadam, Eduardo Calpena, Nagore De León, Marta Hoya, Cristina Doncel, Janet Leatherwood, and M.-Henar Valdivieso. 2014. ‘Membrane Organization and Cell Fusion during Mating in Fission Yeast Requires Multipass Membrane Protein Prm1’. Genetics 196 (4): 1059–76. 10.1534/genetics.113.159558.

Daskalov, Asen. 2023. ‘Emergence of the Fungal Immune System’. iScience 26 (6): 106793. 10.1016/j.isci.2023.106793.

Dyer, Paul S., and Ulrich Kück. 2017. ‘Sex and the Imperfect Fungi’. Microbiology Spectrum 5 (3): 10.1128/microbiolspec.funk-0043-2017. https://doi.org/10.1128/microbiolspec.funk-0043-2017.

Edgar, Ron, Michael Domrachev, and Alex E. Lash. 2002. ‘Gene Expression Omnibus: NCBI Gene Expression and Hybridization Array Data Repository’. Nucleic Acids Research 30 (1): 207–10. 10.1093/nar/30.1.207.

Emms, David M., and Steven Kelly. 2019. ‘OrthoFinder: Phylogenetic Orthology Inference for Comparative Genomics’. Genome Biology 20 (1): 238. 10.1186/s13059-019-1832-y.

Erdman, S., L. Lin, M. Malczynski, and M. Snyder. 1998. ‘Pheromone-Regulated Genes Required for Yeast Mating Differentiation’. The Journal of Cell Biology 140 (3): 461–83. 10.1083/jcb.140.3.461.

Espagne, Eric, Pascale Balhadère, Marie-Louise Penin, Christian Barreau, and Béatrice Turcq. 2002. ‘HET-E and HET-D Belong to a New Subfamily of WD40 Proteins Involved in Vegetative Incompatibility Specificity in the Fungus *Podospora anserina*.’ Genetics 161 (1): 71–81.

Espagne, Eric, Olivier Lespinet, Fabienne Malagnac, Corinne Da Silva, Olivier Jaillon, Betina M Porcel, Arnaud Couloux, et al. 2008. ‘The Genome Sequence of the Model Ascomycete Fungus *Podospora anserina*’. Genome Biology 9 (5): R77. 10.1186/gb-2008-9-5-r77.

Foster, Benjamin M., Paul Stolz, Christopher B. Mulholland, Alex Montoya, Holger Kramer, Sebastian Bultmann, and Till Bartke. 2018. ‘Critical Role of the UBL Domain in Stimulating the E3 Ubiquitin Ligase Activity of UHRF1 toward Chromatin’. Molecular Cell 72 (4): 739–752.e9. 10.1016/j.molcel.2018.09.028.

Froehlich, Allan C., Yi Liu, Jennifer J. Loros, and Jay C. Dunlap. 2002. ‘White Collar-1, a Circadian Blue Light Photoreceptor, Binding to the Frequency Promoter’. Science (New York, N.Y.) 297 (5582): 815–19. 10.1126/science.1073681.

Galagan, James E, Sarah E Calvo, Katherine A Borkovich, Eric U Selker, Nick D Read, David Jaffe, William FitzHugh, et al. 2003. ‘The Genome Sequence of the Filamentous Fungus *Neurospora crassa*’. Nature 422 (6934): 859–68. 10.1038/nature01554.

Gautier, Valérie, Laetitia Chan Ho Tong, Tinh-Suong Nguyen, Robert Debuchy, and Philippe Silar. 2018. ‘PaPro1 and IDC4, Two Genes Controlling Stationary Phase, Sexual Development and Cell Degeneration in *Podospora anserina*’. Journal of Fungi (Basel, Switzerland) 4 (3): 85. 10.3390/jof4030085.

Girard, Chloe, Karine Budin, Stéphanie Boisnard, Liangran Zhang, Robert Debuchy, Denise Zickler, and Eric Espagne. 2021. ‘RNAi-Related Dicer and Argonaute Proteins Play Critical Roles for Meiocyte Formation, Chromosome-Axes Lengths and Crossover Patterning in the Fungus *Sordaria macrospora*’. Frontiers in Cell and Developmental Biology 9 (June). 10.3389/fcell.2021.684108.

Grognet, Pierre, Hélène Timpano, Florian Carlier, Jinane Aït-Benkhali, Véronique Berteaux-Lecellier, Robert Debuchy, Frédérique Bidard, and Fabienne Malagnac. 2019. ‘A RID-like Putative Cytosine Methyltransferase Homologue Controls Sexual Development in the Fungus *Podospora anserina*’. PLoS Genetics 15 (8): e1008086. 10.1371/journal.pgen.1008086.

He, Qiyang, Ping Cheng, Yuhong Yang, Lixing Wang, Kevin H. Gardner, and Yi Liu. 2002. ‘White Collar-1, a DNA Binding Transcription Factor and a Light Sensor’. Science (New York, N.Y.) 297 (5582): 840–43. 10.1126/science.1072795.

Hutchison, Elizabeth A, and N Louise Glass. 2010. ‘Meiotic Regulators Ndt80 and Ime2 Have Different Roles in Saccharomyces and Neurospora’. Genetics 185 (4): 1271–82. 10.1534/genetics.110.117184.

Iliev, Iliyan D., Gordon D. Brown, Petra Bacher, Sarah L. Gaffen, Joseph Heitman, Bruce S. Klein, and Michail S. Lionakis. 2024. ‘Focus on Fungi’. Cell 187 (19): 5121–27. 10.1016/j.cell.2024.08.016.

Imbeaud, Sandrine, Esther Graudens, Virginie Boulanger, Xavier Barlet, Patrick Zaborski, Eric Eveno, Odilo Mueller, Andreas Schroeder, and Charles Auffray. 2005. ‘Towards Standardization of RNA Quality Assessment Using User-Independent Classifiers of Microcapillary Electrophoresis Traces’. Nucleic Acids Research 33 (6): e56. 10.1093/nar/gni054.

Julius, D., A. Brake, L. Blair, R. Kunisawa, and J. Thorner. 1984. ‘Isolation of the Putative Structural Gene for the Lysine-Arginine-Cleaving Endopeptidase Required for Processing of Yeast Prepro-Alpha-Factor’. Cell 37 (3): 1075–89. 10.1016/0092-8674(84)90442-2.

Kaneko, Isao, Karine Dementhon, Qijun Xiang, and N. Louise Glass. 2006. ‘Nonallelic Interactions between Het-c and a Polymorphic Locus, Pin-c, Are Essential for Nonself Recognition and Programmed Cell Death in *Neurospora crassa*’. Genetics 172 (3): 1545–55. 10.1534/genetics.105.051490.

Kerr, M. K., and G. A. Churchill. 2001. ‘Statistical Design and the Analysis of Gene Expression Microarray Data’. Genetical Research 77 (2): 123–28. 10.1017/s0016672301005055.

Kim, Wonyong, Zheng Wang, Hyeonjae Kim, Kasey Pham, Yujia Tu, Jeffrey P. Townsend, and Frances Trail. 2022. ‘Transcriptional Divergence Underpinning Sexual Development in the Fungal Class Sordariomycetes’. mBio, May, e0110022. 10.1128/mbio.01100-22.

Kurahashi, Hiroshi, Yoshiyuki Imai, and Masayuki Yamamoto. 2002. ‘Tropomyosin Is Required for the Cell Fusion Process during Conjugation in Fission Yeast’. Genes to Cells: Devoted to Molecular & Cellular Mechanisms 7 (4): 375–84. 10.1046/j.1365-2443.2002.00526.x.

Lamacchia, Marina, Witold Dyrka, Annick Breton, Sven J. Saupe, and Mathieu Paoletti. 2016. ‘Overlapping *Podospora anserina* Transcriptional Responses to Bacterial and Fungal Non Self Indicate a Multilayered Innate Immune Response’. Frontiers in Microbiology 7:471. 10.3389/fmicb.2016.00471.

Leipe, Detlef D., Eugene V. Koonin, and L. Aravind. 2004. ‘STAND, a Class of P-Loop NTPases Including Animal and Plant Regulators of Programmed Cell Death: Multiple, Complex Domain Architectures, Unusual Phyletic Patterns, and Evolution by Horizontal Gene Transfer’. Journal of Molecular Biology 343 (1): 1–28. 10.1016/j.jmb.2004.08.023.

Li, Wan-Chen, Chia-Ling Chen, and Ting-Fang Wang. 2018. ‘Repeat-Induced Point (RIP) Mutation in the Industrial Workhorse Fungus *Trichoderma reesei*’. Applied Microbiology and Biotechnology, January. 10.1007/s00253-017-8731-5.

Liu, Li, Christoph Sasse, Benedict Dirnberger, Oliver Valerius, Enikő Fekete-Szücs, Rebekka Harting, Daniela E Nordzieke, et al. 2021. ‘Secondary Metabolites of Hülle Cells Mediate Protection of Fungal Reproductive and Overwintering Structures against Fungivorous Animals’. Edited by Antonis Rokas, Detlef Weigel, Milton Drott, and Jae-Hyuk Yu. eLife 10 (October):e68058. 10.7554/eLife.68058.

Lütkenhaus, Ramona, Stefanie Traeger, Jan Breuer, Laia Carreté, Alan Kuo, Anna Lipzen, Jasmyn Pangilinan, et al. 2019. ‘Comparative Genomics and Transcriptomics to Analyze Fruiting Body Development in Filamentous Ascomycetes’. Genetics, October. 10.1534/genetics.119.302749.

MacPherson, Sarah, Marc Larochelle, and Bernard Turcotte. 2006. ‘A Fungal Family of Transcriptional Regulators: The Zinc Cluster Proteins’. Microbiology and Molecular Biology Reviews 70 (3): 583–604. 10.1128/mmbr.00015-06.

Malagnac, F, B Wendel, C Goyon, G Faugeron, D Zickler, J L Rossignol, M Noyer-Weidner, P Vollmayr, T A Trautner, and J Walter. 1997. ‘A Gene Essential for de Novo Methylation and Development in Ascobolus Reveals a Novel Type of Eukaryotic DNA Methyltransferase Structure’. Cell 91 (2): 281–90.

Mistry, Jaina, Sara Chuguransky, Lowri Williams, Matloob Qureshi, Gustavo A. Salazar, Erik L. L. Sonnhammer, Silvio C. E. Tosatto, et al. 2021. ‘Pfam: The Protein Families Database in 2021’. Nucleic Acids Research 49 (D1): D412–19. 10.1093/nar/gkaa913.

Nelson, M. A., S. T. Merino, and R. L. Metzenberg. 1997. ‘A Putative Rhamnogalacturonase Required for Sexual Development of *Neurospora crassa*’. Genetics 146 (2): 531–40. 10.1093/genetics/146.2.531.

Nowrousian, Minou. 2009. ‘A Novel Polyketide Biosynthesis Gene Cluster Is Involved in Fruiting Body Morphogenesis in the Filamentous Fungi *Sordaria macrospora* and *Neurospora crassa*’. Current Genetics 55 (2): 185–98. 10.1007/s00294-009-0236-z.

Nowrousian, Minou. 2022. ‘The Role of Chromatin and Transcriptional Control in the Formation of Sexual Fruiting Bodies in Fungi’. Microbiology and Molecular Biology Reviews: MMBR 86 (4): e0010422. 10.1128/mmbr.00104-22.

Oh, Yong-Seok, Pu Gao, Ko-Woon Lee, Ilaria Ceglia, Ji-Seon Seo, Xiaozhu Zhang, Jung-Hyuck Ahn, et al. 2013. ‘SMARCA3, a Chromatin-Remodeling Factor, Is Required for P11-Dependent Antidepressant Action’. Cell 152 (4): 831–43. 10.1016/j.cell.2013.01.014.

Ottmann, Christian, Borries Luberacki, Isabell Küfner, Wolfgang Koch, Frédéric Brunner, Michael Weyand, Laura Mattinen, et al. 2009. ‘A Common Toxin Fold Mediates Microbial Attack and Plant Defense’. Proceedings of the National Academy of Sciences 106 (25): 10359–64. 10.1073/pnas.0902362106.

Otto, Sarah P. 2009. ‘The Evolutionary Enigma of Sex’. The American Naturalist 174 Suppl 1 (July):S1–14. 10.1086/599084.

Pearl, LH. 2000. ‘Structure and Function in the Uracil-DNA Glycosylase Superfamily’. Mutation Research 460 (3–4). 10.1016/s0921-8777(00)00025-2.

Peraza Reyes, Leonardo, and Véronique Berteaux-Lecellier. 2013. ‘Peroxisomes and Sexual Development in Fungi’. Frontiers in Physiology 4. 10.3389/fphys.2013.00244.

Peraza-Reyes, Leonardo, and Fabienne Malagnac. 2016. Sexual Development in Fungi. The Mycota I, Growth, Differentiation and Sexuality. Springer.

Rizet, G, and C Engelmann. 1949. ‘Contribution à l’étude Génétique d’un Ascomycète Tétrasporé: *Podospora anserina*’ 11:201–304.

Romano, Nicoletta, and Giuseppe Macino. 1992. ‘Quelling: Transient Inactivation of Gene Expression in *Neurospora crassa* by Transformation with Homologous Sequences’. Molecular Microbiology 6 (22): 3343–53. 10.1111/j.1365-2958.1992.tb02202.x.

Rose, M. D., L. M. Misra, and J. P. Vogel. 1989. ‘KAR2, a Karyogamy Gene, Is the Yeast Homolog of the Mammalian BiP/GRP78 Gene’. Cell 57 (7): 1211–21. 10.1016/0092-8674(89)90058-5.

Shen, Ling, François-Hugues Porée, Thomas Gaslonde, Hervé Lalucque, Florence Chapeland-Leclerc, and Gwenaël Ruprich-Robert. 2019. ‘Functional Characterization of the Sterigmatocystin Secondary Metabolite Gene Cluster in the Filamentous Fungus *Podospora anserina*: Involvement in Oxidative Stress Response, Sexual Development, Pigmentation and Interspecific Competitions’. Environmental Microbiology 21 (8): 3011–26. 10.1111/1462-2920.14698.

Shiu, P K, N B Raju, D Zickler, and R L Metzenberg. 2001. ‘Meiotic Silencing by Unpaired DNA’. Cell 107 (7): 905–16.

Silar, Philippe. 2020. *Podospora anserina*. https://hal.archives-ouvertes.fr/hal-02475488.

Slot, Jason C., and Antonis Rokas. 2011. ‘Horizontal Transfer of a Large and Highly Toxic Secondary Metabolic Gene Cluster between Fungi’. Current Biology: CB 21 (2): 134–39. 10.1016/j.cub.2010.12.020.

Stajich, Jason E., Mary L. Berbee, Meredith Blackwell, David S. Hibbett, Timothy Y. James, Joseph W. Spatafora, and John W. Taylor. 2009. ‘The Fungi’. Current Biology 19 (18): R840–45. 10.1016/j.cub.2009.07.004.

Steffens, Eva Katharina, Kordula Becker, Sabine Krevet, Ines Teichert, and Ulrich Kück. 2016. ‘Transcription Factor PRO1 Targets Genes Encoding Conserved Components of Fungal Developmental Signaling Pathways’. Molecular Microbiology 102 (5): 792–809. 10.1111/mmi.13491.

Stergiopoulos, Ioannis, Jérôme Collemare, Rahim Mehrabi, and Pierre J. G. M. De Wit. 2013. ‘Phytotoxic Secondary Metabolites and Peptides Produced by Plant Pathogenic Dothideomycete Fungi’. FEMS Microbiology Reviews 37 (1): 67–93. 10.1111/j.1574-6976.2012.00349.x.

Sun, Xianyun, Luning Yu, Nan Lan, Shiping Wei, Yufei Yu, Hanxing Zhang, Xinyu Zhang, and Shaojie Li. 2012. ‘Analysis of the Role of Transcription Factor VAD-5 in Conidiation of *Neurospora crassa*’. Fungal Genetics and Biology 49 (5): 379–87. 10.1016/j.fgb.2012.03.003.

Teichert, Ines, Tim A. Dahlmann, Ulrich Kück, and Minou Nowrousian. 2017. ‘RNA Editing During Sexual Development Occurs in Distantly Related Filamentous Ascomycetes’. Genome Biology and Evolution 9 (4): 855–68. 10.1093/gbe/evx052.

Teichert, Ines, Stefanie Pöggeler, and Minou Nowrousian. 2020. ‘*Sordaria macrospora*: 25 Years as a Model Organism for Studying the Molecular Mechanisms of Fruiting Body Development’. Applied Microbiology and Biotechnology 104 (9): 3691–3704. 10.1007/s00253-020-10504-3.

Teichert, Ines, Gabriele Wolff, Ulrich Kück, and Minou Nowrousian. 2012. ‘Combining Laser Microdissection and RNA-Seq to Chart the Transcriptional Landscape of Fungal Development’. BMC Genomics 13 (1): 511. 10.1186/1471-2164-13-511.

The Gene Ontology Consortium, Suzi A Aleksander, James Balhoff, Seth Carbon, J Michael Cherry, Harold J Drabkin, Dustin Ebert, et al. 2023. ‘The Gene Ontology Knowledgebase in 2023’. Genetics 224 (1): iyad031. 10.1093/genetics/iyad031.

Thompson-Coffe, C., G. Borioli, D. Zickler, and A. L. Rosa. 1999. ‘Pyruvate Decarboxylase Filaments Are Associated with the Cortical Cytoskeleton of Asci and Spores over the Sexual Cycle of Filamentous Ascomycetes’. Fungal Genetics and Biology: FG & B 26 (1): 71–80. 10.1006/fgbi.1998.1106.

Tong, Kevin, Thomas Keller, Charles S. Hoffman, and Anthony T. Annunziato. 2012. ‘*Schizosaccharomyces pombe* Hat1 (Kat1) Is Associated with Mis16 and Is Required for Telomeric Silencing’. Eukaryotic Cell 11 (9): 1095–1103. 10.1128/EC.00123-12.

Turgeon, B Gillian, and R. Debuchy. 2007. ‘Cochliobolus and Podospora: Mechanism of Sex Determination and the Evolution of Reproductive Lifestyle’. In Sex in Fungi, Molecular Determination and Evolutionary Implications.

Wang, Amy, Siavash K. Kurdistani, and Michael Grunstein. 2002. ‘Requirement of Hos2 Histone Deacetylase for Gene Activity in Yeast’. Science (New York, N.Y.) 298 (5597): 1412–14. 10.1126/science.1077790.

Wang, Chengshu, and Raymond J. St Leger. 2007. ‘The MAD1 Adhesin of Metarhizium Anisopliae Links Adhesion with Blastospore Production and Virulence to Insects, and the MAD2 Adhesin Enables Attachment to Plants’. Eukaryotic Cell 6 (5): 808–16. 10.1128/EC.00409-06.

Wang, Peng, Jia Xu, Perng-Kuang Chang, Zhemin Liu, and Qing Kong. 2022. ‘New Insights of Transcriptional Regulator AflR in *Aspergillus flavus* Physiology’. Microbiology Spectrum 10 (1): e00791–21. 10.1128/spectrum.00791-21.

Wang, Zheng, Aditya Gudibanda, Ugochukwu Ugwuowo, Frances Trail, and Jeffrey P. Townsend. 2018. ‘Using Evolutionary Genomics, Transcriptomics, and Systems Biology to Reveal Gene Networks Underlying Fungal Development’. *Fungal Biology Reviews*, Complex multicellularity in Fungi, 32 (4): 249–64. 10.1016/j.fbr.2018.02.001.

Wang, Zheng, Francesc Lopez-Giraldez, Nina Lehr, Marta Farré, Ralph Common, Frances Trail, and Jeffrey P Townsend. 2014. ‘Global Gene Expression and Focused Knockout Analysis Reveals Genes Associated with Fungal Fruiting Body Development in *Neurospora crassa*’. Eukaryotic Cell 13 (1): 154–69. 10.1128/EC.00248-13.

Wang, Zheng, Francesc Lopez-Giraldez, Jason Slot, Oded Yarden, Frances Trail, and Jeffrey P. Townsend. 2022. ‘Secondary Metabolism Gene Clusters Exhibit Increasingly Dynamic and Differential Expression during Asexual Growth, Conidiation, and Sexual Development in *Neurospora crassa*’. mSystems 7 (3): e00232–22. 10.1128/msystems.00232-22.

Wang, Zheng, Francesc López-Giráldez, Junrui Wang, Frances Trail, and Jeffrey P. Townsend. 2019. ‘Integrative Activity of Mating Loci, Environmentally Responsive Genes, and Secondary Metabolism Pathways during Sexual Development of *Chaetomium globosum*’. mBio 10 (6): 10.1128/mbio.02119-19. https://doi.org/10.1128/mbio.02119-19.

White, B. T., and C. Yanofsky. 1993. ‘Structural Characterization and Expression Analysis of the Neurospora Conidiation Gene Con-6’. Developmental Biology 160 (1): 254–64. 10.1006/dbio.1993.1303.

Wilson, Andi M., P. Markus Wilken, Magriet A. van der Nest, Michael J. Wingfield, and Brenda D. Wingfield. 2019. ‘It’s All in the Genes: The Regulatory Pathways of Sexual Reproduction in Filamentous Ascomycetes’. Genes 10 (5): 330. 10.3390/genes10050330.

Wilson, Andi M., P. Markus Wilken, Michael J. Wingfield, and Brenda D. Wingfield. 2021. ‘Genetic Networks That Govern Sexual Reproduction in the Pezizomycotina’. Microbiology and Molecular Biology Reviews 85 (4): e00020–21. 10.1128/MMBR.00020-21.

Wolfe, Cecily J., Isaac S. Kohane, and Atul J. Butte. 2005. ‘Systematic Survey Reveals General Applicability of “Guilt-by-Association” within Gene Coexpression Networks’. BMC Bioinformatics 6 (1): 227. 10.1186/1471-2105-6-227.

Yang, Qiuying, Qiaohong Anne Ye, and Yi Liu. 2015. ‘Mechanism of siRNA Production from Repetitive DNA’. Genes & Development 29 (5): 526–37. 10.1101/gad.255828.114.

Young, B. P., R. A. Craven, P. J. Reid, M. Willer, and C. J. Stirling. 2001. ‘Sec63p and Kar2p Are Required for the Translocation of SRP-Dependent Precursors into the Yeast Endoplasmic Reticulum in Vivo’. The EMBO Journal 20 (1–2): 262–71. 10.1093/emboj/20.1.262.

Zhao, Yanxia, Jianing Ding, Weihua Yuan, Jinjin Huang, Wenxiu Huang, Yan Wang, and Weifa Zheng. 2017. ‘Production of a Fungal Furocoumarin by a Polyketide Synthase Gene Cluster Confers the Chemo-Resistance of *Neurospora crassa* to the Predation by Fungivorous Arthropods’. Environmental Microbiology 19 (10): 3920–29. 10.1111/1462-2920.13791.

Zheng, Qian, Rui Hou, null Juanyu, null Zhang, Jiwen Ma, Zhongshou Wu, Guanghui Wang, Chenfang Wang, and Jin-Rong Xu. 2013. ‘The MAT Locus Genes Play Different Roles in Sexual Reproduction and Pathogenesis in *Fusarium graminearum*’. PloS One 8 (6): e66980. 10.1371/journal.pone.0066980.

Zickler, D., S. Arnaise, E. Coppin, R. Debuchy, and M. Picard. 1995. ‘Altered Mating-Type Identity in the Fungus *Podospora anserina* Leads to Selfish Nuclei, Uniparental Progeny, and Haploid Meiosis’. Genetics 140 (2): 493–503.

