## Supplemental figure file for "Sexual reproduction is controlled by successive transcriptomic waves in *Podospora anserina*"

### Graphical summary of data analysis

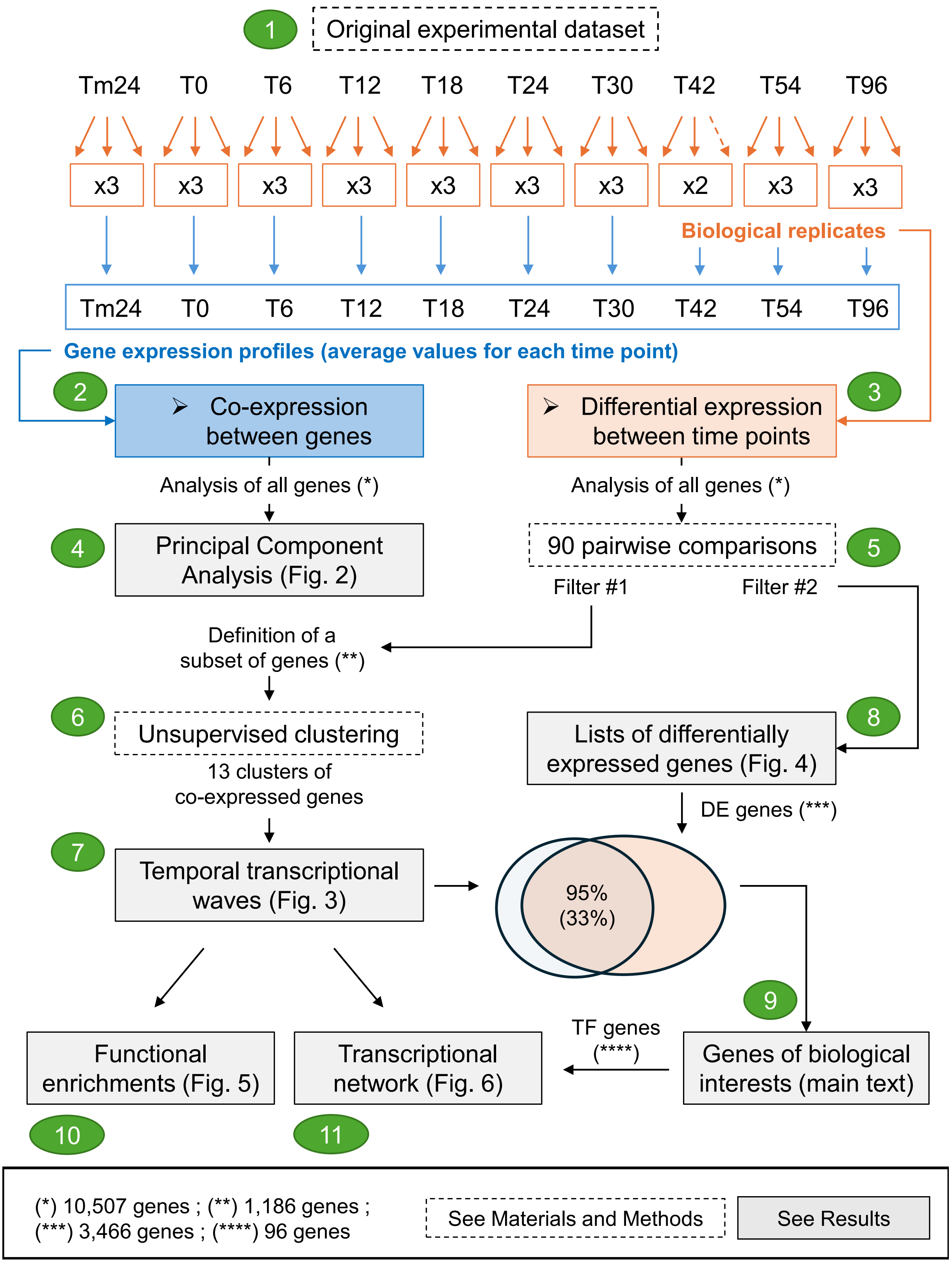

**Supplementary Figure 1: Simplified representation of the data analyses performed in this study**. **(1)** The dataset consists of 10 time points named Tm24, T0, T6, T12, T18, T24, T30, T42, T54 and T96 (see the main text). For each time point, gene expression levels were measured for all genes, in triplicate (orange boxes), except for time point T42, for which the third replicate was of insufficient quality. At each time point, triplicate values were averaged to define expression profiles for each gene (blue box). Thus, a gene expression profile consists of ten different values (Tm24, T0, T6, T12, T18, T24, T30, T42, T54 and T96) that represent the expression fluctuations of the gene over the time course. Expression profiles were used to study co-expression between genes, that is, to identify genes for which the fluctuations during the time course exhibit similar patterns **(2)**, and biological replicates were used to search for genes that were differentially expressed between any two time points in the time course **(3)**. (2) and (3) represent two different strategies for analyzing the original experimental dataset. Both analyses started by examining all genes (*i.e.* 10,507 genes) for which expression levels were available in (1). The expression profiles were used to perform principal component analysis, the results of which are shown in Figure 2 (main text), and the biological replicates were used to perform 90 pairwise comparisons to search for differentially expressed genes (see Methods). Two filters were used to select genes of interest for further investigation. The first filter (see Methods) allowed the definition of a subset of genes (*i.e.* 1,186 genes) for which significant fluctuations in expression were observed between the different stages of *P. anserina* sexual reproduction. The co-expression between them was analyzed using unsupervised clustering **(6)**. After several attempts and careful manual inspection of the results (see Methods), 13 clusters appeared to be a good compromise between minimizing intra-cluster variability between expression profiles and maximizing inter-cluster variability. Most of the exploration of the data was done with these 13 clusters. The 5 temporal waves shown in Figure 3 were created (see Methods) to summarize the main characteristics of the clusters **(7)**. A second filter (see Methods) allowed the generation of lists of differentially expressed genes **(8)**. Compared to the first filter, it relies on a selection of differentially expressed genes based on a stricter threshold on the adjusted p-value (whereas the first filter relied on a stricter threshold on the overall amplitude of fluctuations over time). In total, the lists of differentially expressed genes contained 3,466 genes, of which 33% also belonged to the temporal waves (and 95% of the genes in the temporal waves belonged to differentially expressed genes). Such an overlap underlines the interest of both data analysis strategies (**(2)** and **(3)**), each of which has specificities but remains relevant to the other. On this basis, a careful inspection of genes of biological interest was carried out **(9)**, allowing the interpretation (see the main text) of the functional enrichments observed in the temporal waves **(10)** and the construction of a transcriptional network **(11)**.

**A**

| Experiment N° | T0 | T0 | T0.5h | T1h | T2h | T3h | T6h | T10h |
|---|---|---|---|---|---|---|---|---|
| 1 | WT | 136 | | | | | | |
| 2 | WT | - | 136 | | | | | |
| 3 | WT | - | - | 136 | | | | |
| 4 | WT | - | - | - | 136 | | | |
| 5 | WT | - | - | - | - | 136 | | |
| 6 | WT | - | - | - | - | - | 136 | |
| 7 | WT | - | - | - | - | - | - | 136 |

**B**

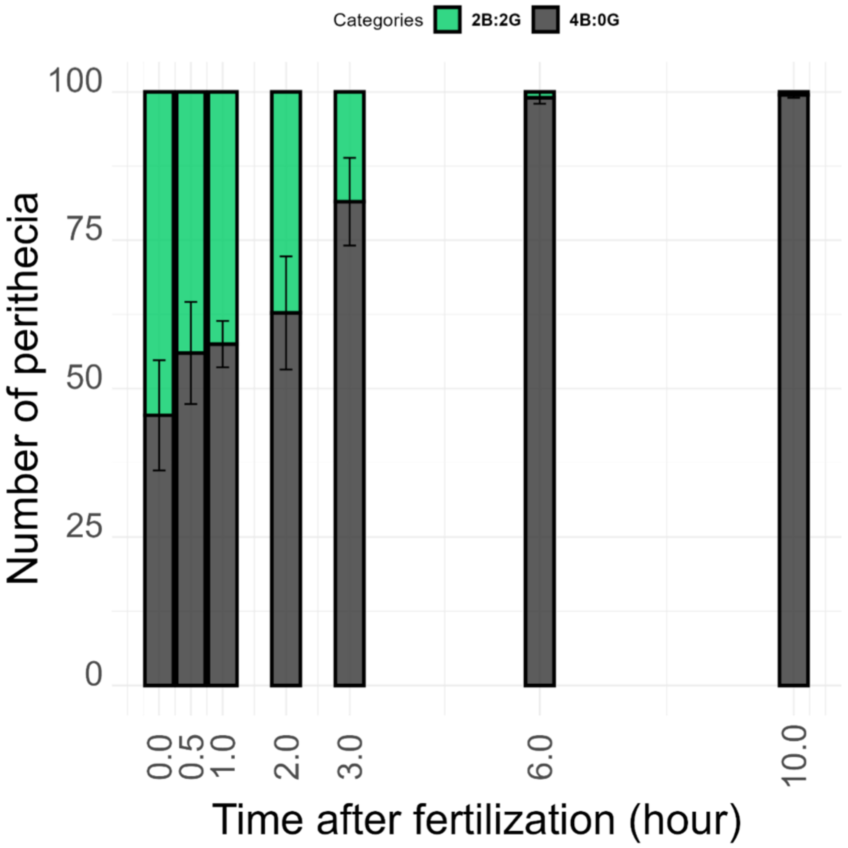


**Supplementary Figure 2: Fertilization kinetics and features.**

We tested whether a given female gamete can be fertilized multiple times. To this hand, we set up three separate fertilization experiments. The first one consisted in a control experiment where wild-type female gametes were fertilized by wild-type male gametes. Five days later, it gave rise to fructifications (perithecia, Fig. 1) each filled with nearly 200 asci containing four black ascospores (4B:0W). The second control experiment consisted in fertilizing wild-type female gametes by male gametes issued from a mutant strain that produces green ascospores (the "*136*" strain, mutated in the *pks1* gene). This latter setting gave rise to fructifications each filled with nearly 200 asci containing 2 green and 2 black ascospores (2B:2G, as well as rare 4B:0W corresponding to the 2% second division segregation of the *pks1* locus). We then proceed to the test experiment where wild-type female gametes were fertilized by a mix of wild-type and *136* mutant spermatia in equal proportion (~ 2 x $1.5.10^5$ spermatia/cross). Five days post-fertilization, we characterized the ascus segregation patterns of over 600 fructifications. None of them showed mixed progeny within a single fructification, *i.e.* both 4B:0W and 2B:2G asci. We then performed kinetics experiments to ask whether after an initial massive fertilization event (T0, Fig. S1A), some of the remaining unfertilized female gametes could still be competent for fertilization. We first performed control experiments where the initial fertilization was made using both wild-type and *136* mutant male gametes in equal proportion (Fig. S1A). Five days later, we examined the content of 60 to 110 fructifications per cross (N=5 x 2 independent experiments, Fig. S1B). When the initial fertilization (T0) was made using both wild-type and *136* mutant male gametes in equal proportion, about half of the fructifications produced 4B:0W asci while the other half produced 2B:2G asci. We then performed a series of crosses for which the initial fertilization even (T0) was made using wild-type male gametes only. Then for each of the 6 time points of the kinetics, subsequent additions of *136* mutant male gametes were made from T0.5 (30 minutes) up to T10 (ten hours) (Fig. S1A). Under such conditions (Fig. S1A), about 80% of the fructifications showed 4B:0W asci at T3 (3 hours), reaching 100% at T6 (6 hours). Accordingly, we showed that fructifications generating 2B:2G asci accounted for 20% only at T3 and were not present anymore at T6 (Fig. S1B).

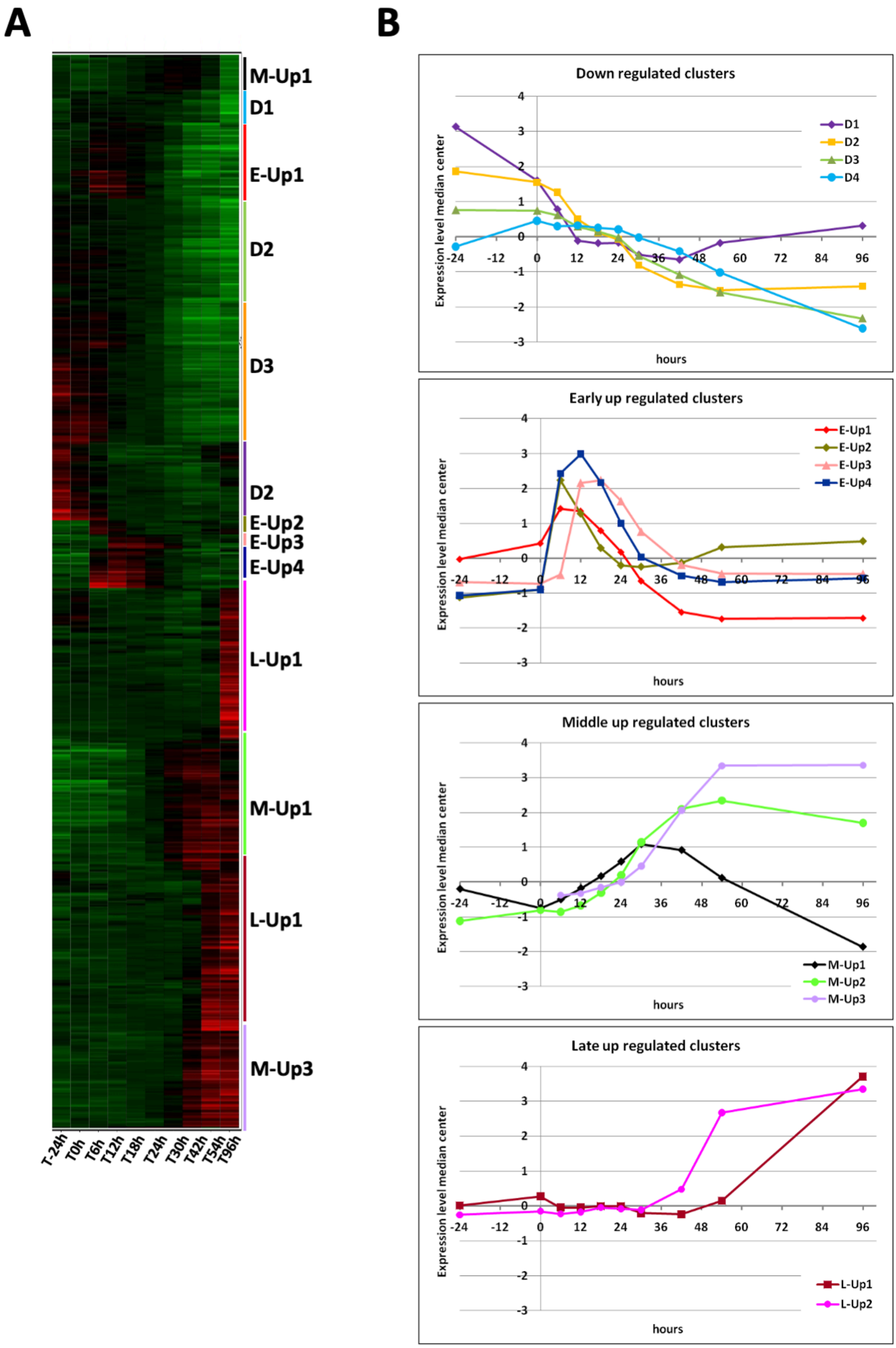
A
B
M-Up1
D1
E-Up1
D2
D3
D2
E-Up2
E-Up3
E-Up4
L-Up1
M-Up1
L-Up1
M-Up3
T-24h
T0h
T6h
T12h
T18h
T24h
T30h
T42h
T54h
T96h
Down regulated clusters
Expression level median center
hours
D1
D2
D3
D4
Early up regulated clusters
E-Up1
E-Up2
E-Up3
E-Up4
Middle up regulated clusters
M-Up1
M-Up2
M-Up3
Late up regulated clusters
L-Up1
L-Up2

**Supplementary Figure 3: Clustered patterns of gene expression during sexual development.**

**A.** Expression patterns of down regulated (green) or up regulated (red) with columns representing experimental time points and rows representing genes. The brightness of the colors is proportional to the level of regulation. The bars on the right side show the approximate position of the expression clusters colored as in B.

**B.** Average expression profiles of the 13 clusters classified in four groups Down (D1, D2, D3, D4), Early (E-Up1, E-Up2, E-Up3, E-Up4), Middle (M-Up1, M-Up2, M-Up3) and Late (L-Up1, L-Up2).

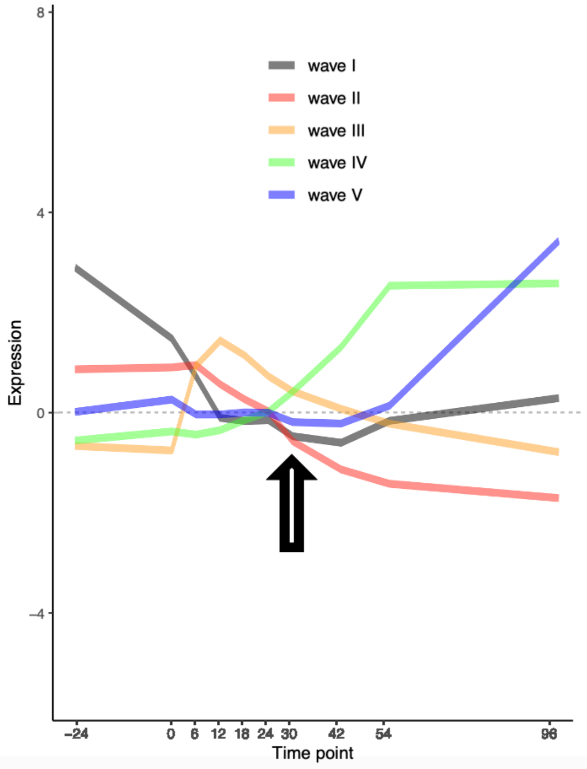


**Supplementary Figure 4: Superimposition of the five waves.**

X-axis: sexual reproduction timeline in hours, where 0 is the original time of fertilization. Y-axis, normalized expression profiles of genes (See Methods). Arrow: common pivot point.

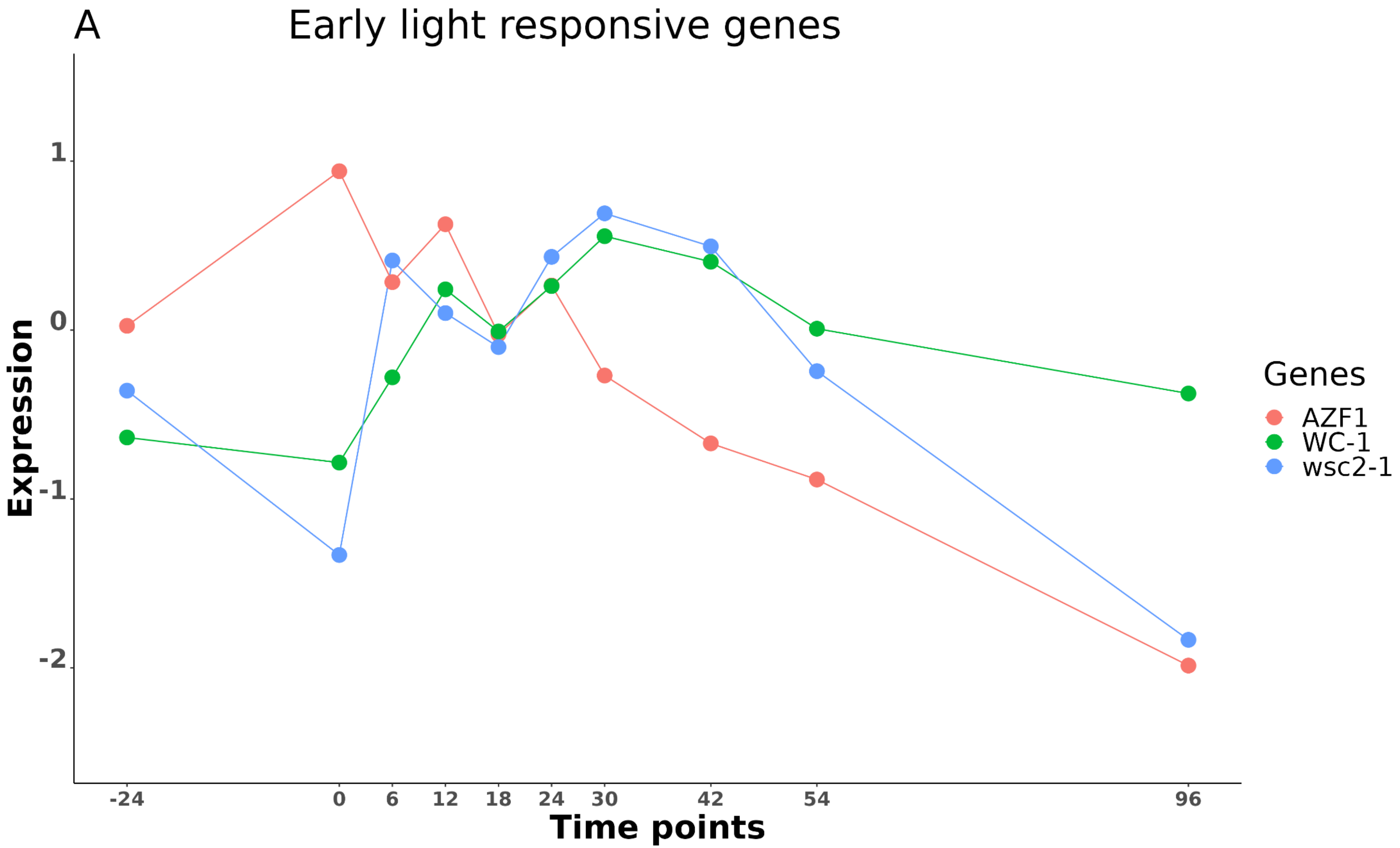

A
Early light responsive genes
Expression
1
0
-1
-2
-24
0
6
12
18
24
30
42
54
96
Time points
Genes
AZF1
WC-1
wsc2-1

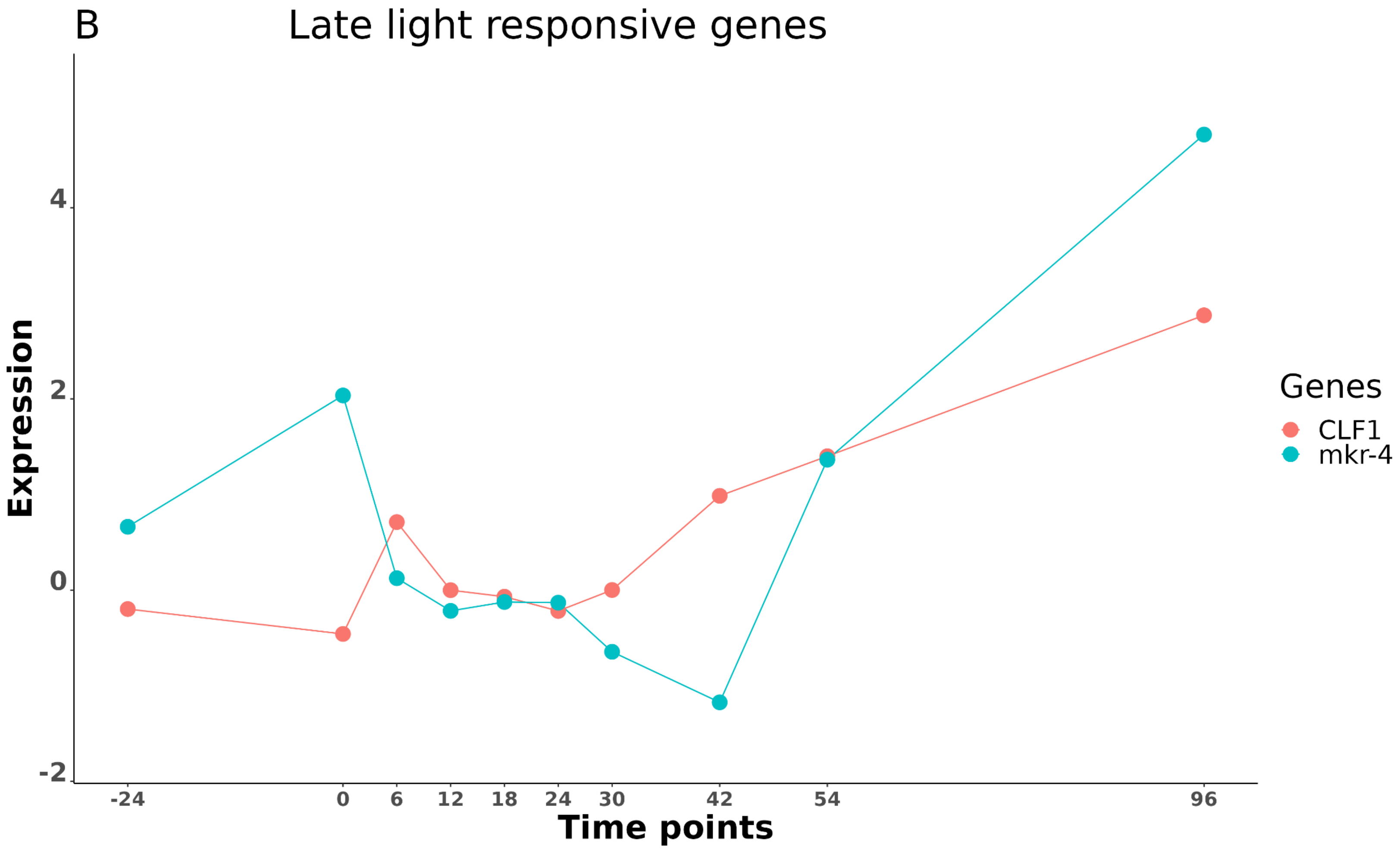

B
Late light responsive genes
Expression
4
2
0
-2
-24
0
6
12
18
24
30
42
54
96
Time points
Genes
CLF1
mkr-4

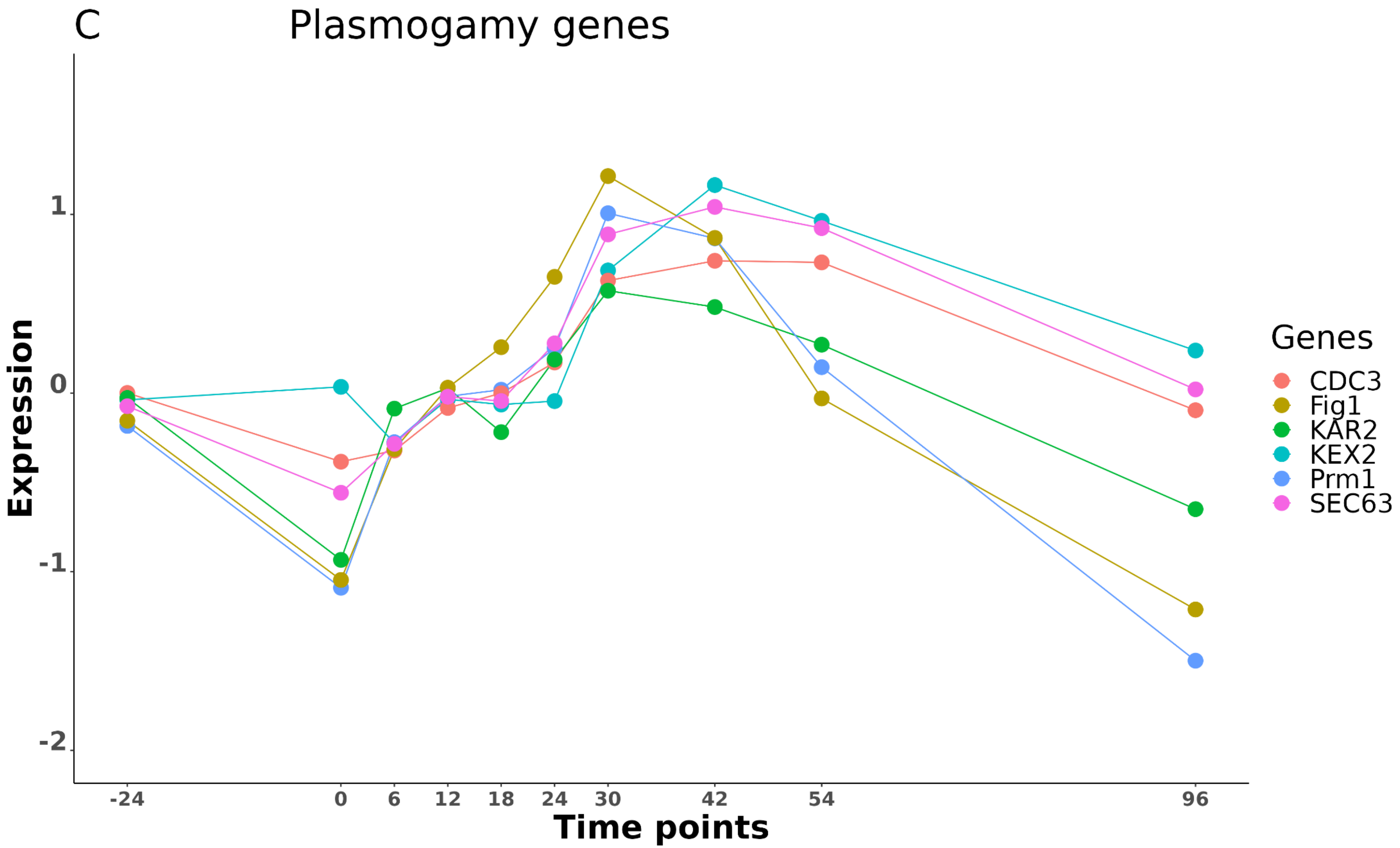
C
Plasmogamy genes
Expression
1
0
-1
-2
-24
0
6
12
18
24
30
42
54
96
Time points
Genes
CDC3
Fig1
KAR2
KEX2
Prm1
SEC63

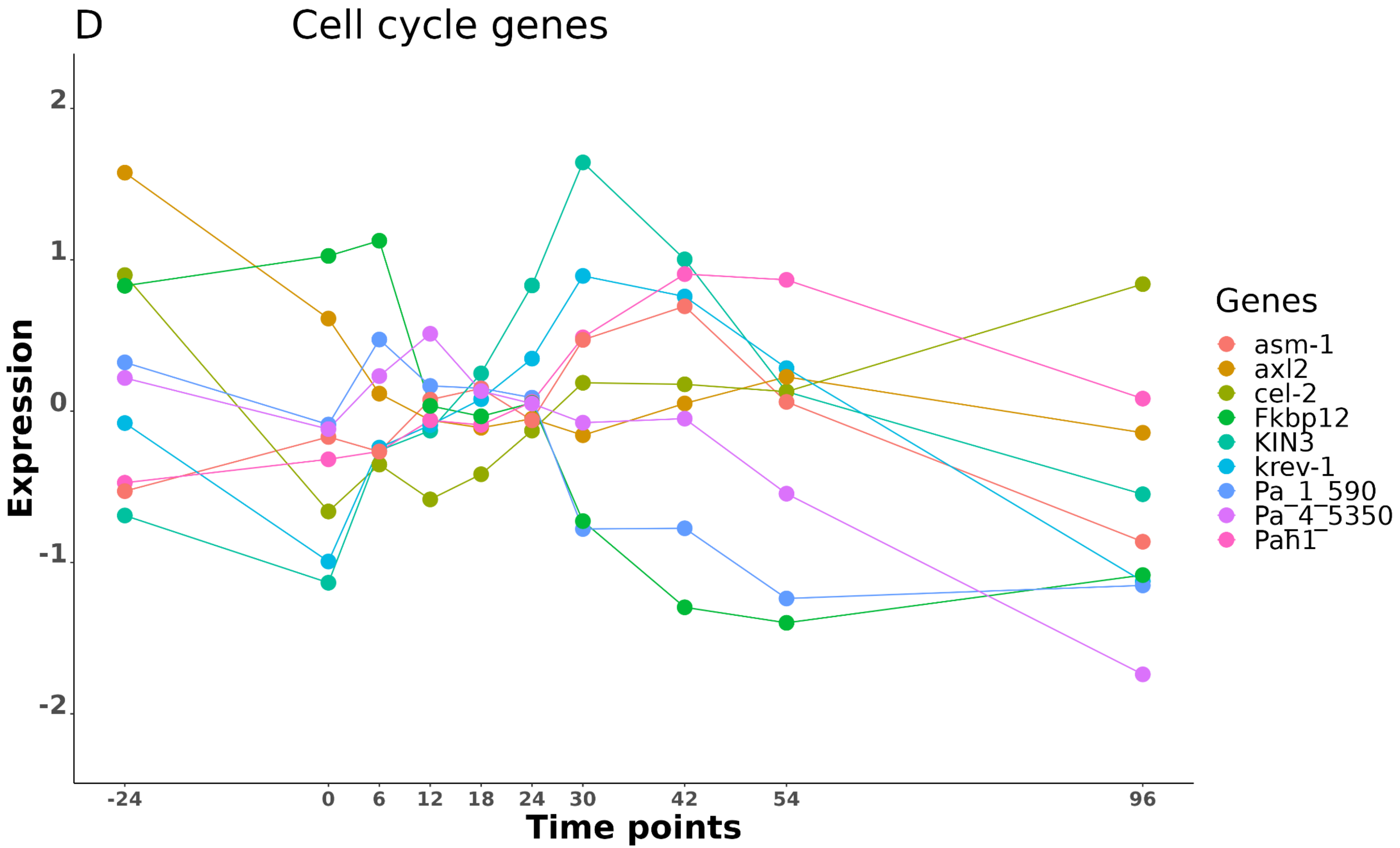

D
Cell cycle genes
Expression
Time points
2
1
0
-1
-2
-24
0
6
12
18
24
30
42
54
96
Genes
asm-1
axl2
cel-2
Fkbp12
KIN3
krev-1
Pa_1_590
Pa_4_5350
Pah1

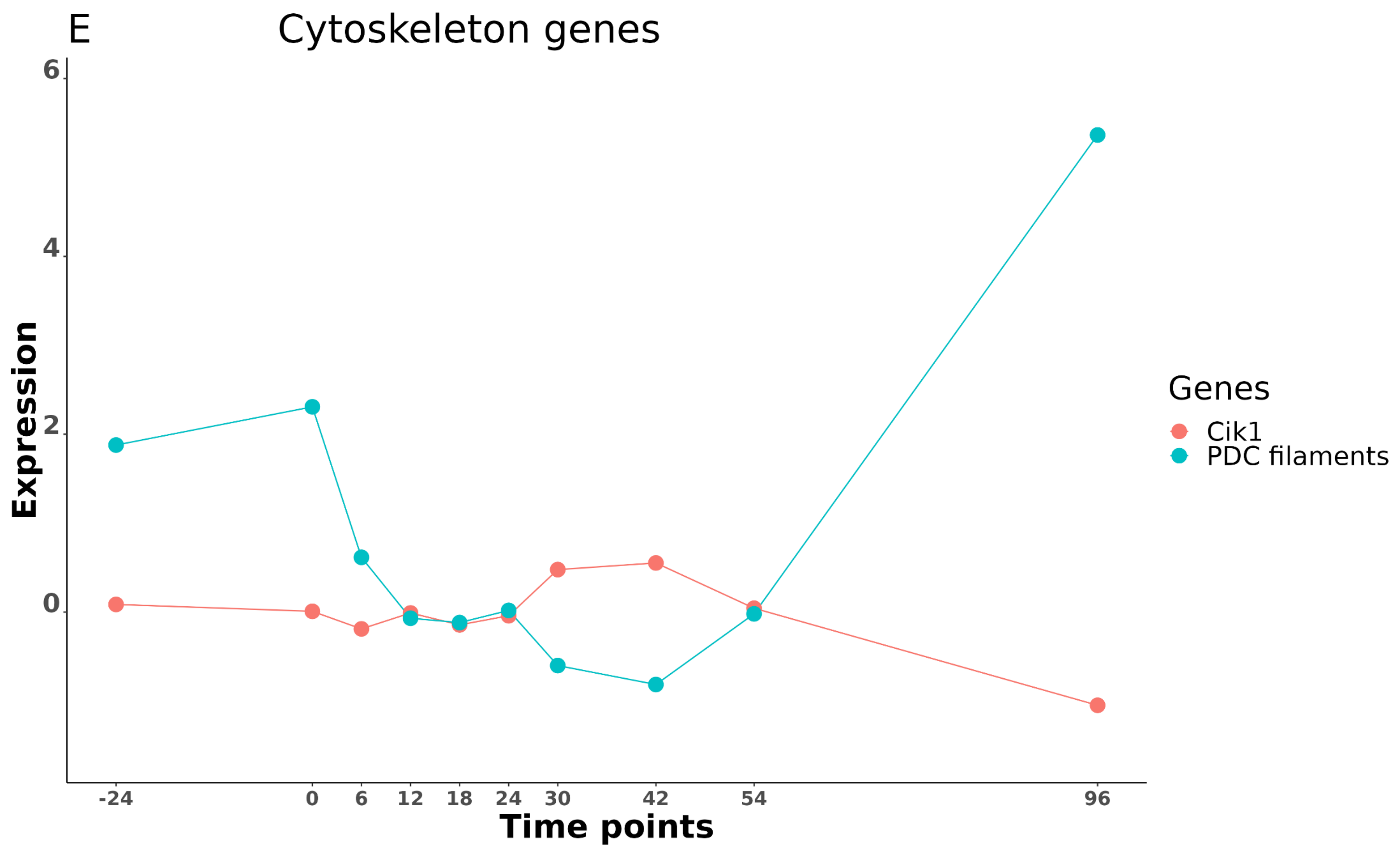
E
Cytoskeleton genes
Expression
6
4
2
0
-24
0
6
12
18
24
30
42
54
96
Time points
Genes
Cik1
PDC filaments

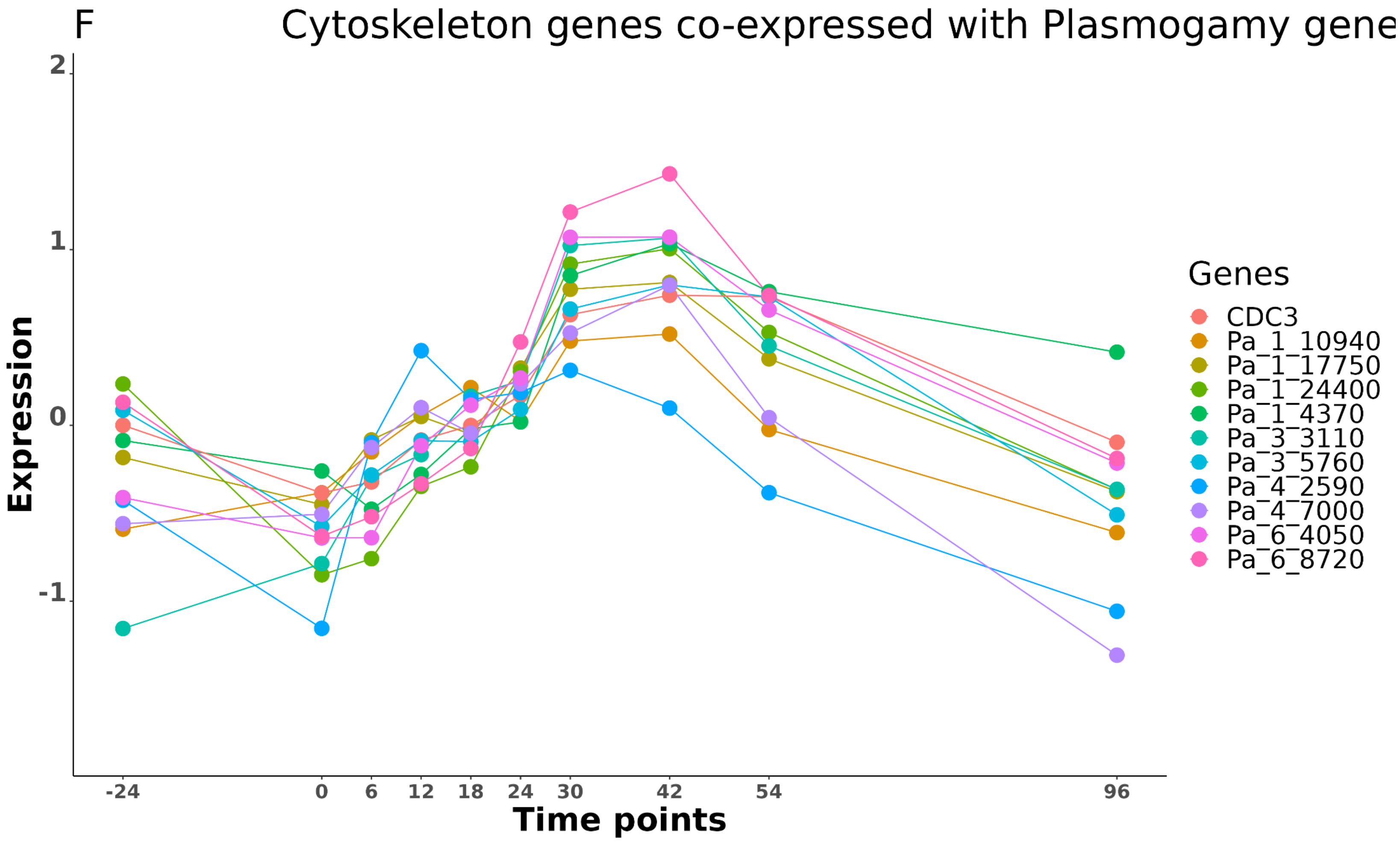

F
Cytoskeleton genes co-expressed with Plasmogamy gene
Expression
2
1
0
-1
-24
0
6
12
18
24
30
42
54
96
Time points
Genes
CDC3
Pa_1_10940
Pa_1_17750
Pa_1_24400
Pa_1_4370
Pa_3_3110
Pa_3_5760
Pa_4_2590
Pa_4_7000
Pa_6_4050
Pa_6_8720

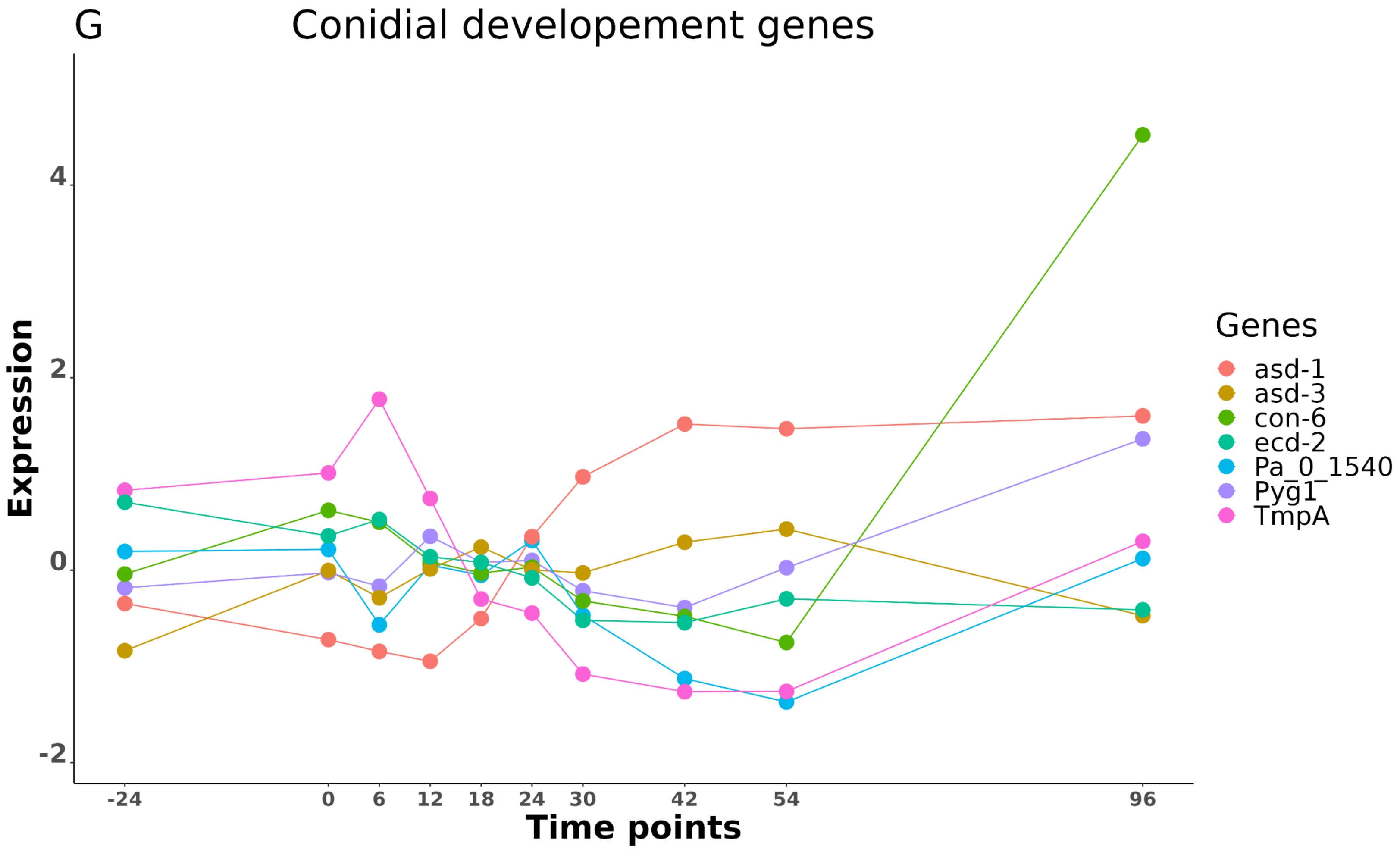
G
Conidial developement genes
Expression
4
2
0
-2
-24
0
6
12
18
24
30
42
54
96
Time points
Genes
asd-1
asd-3
con-6
ecd-2
Pa_0_1540
Pyg1
TmpA

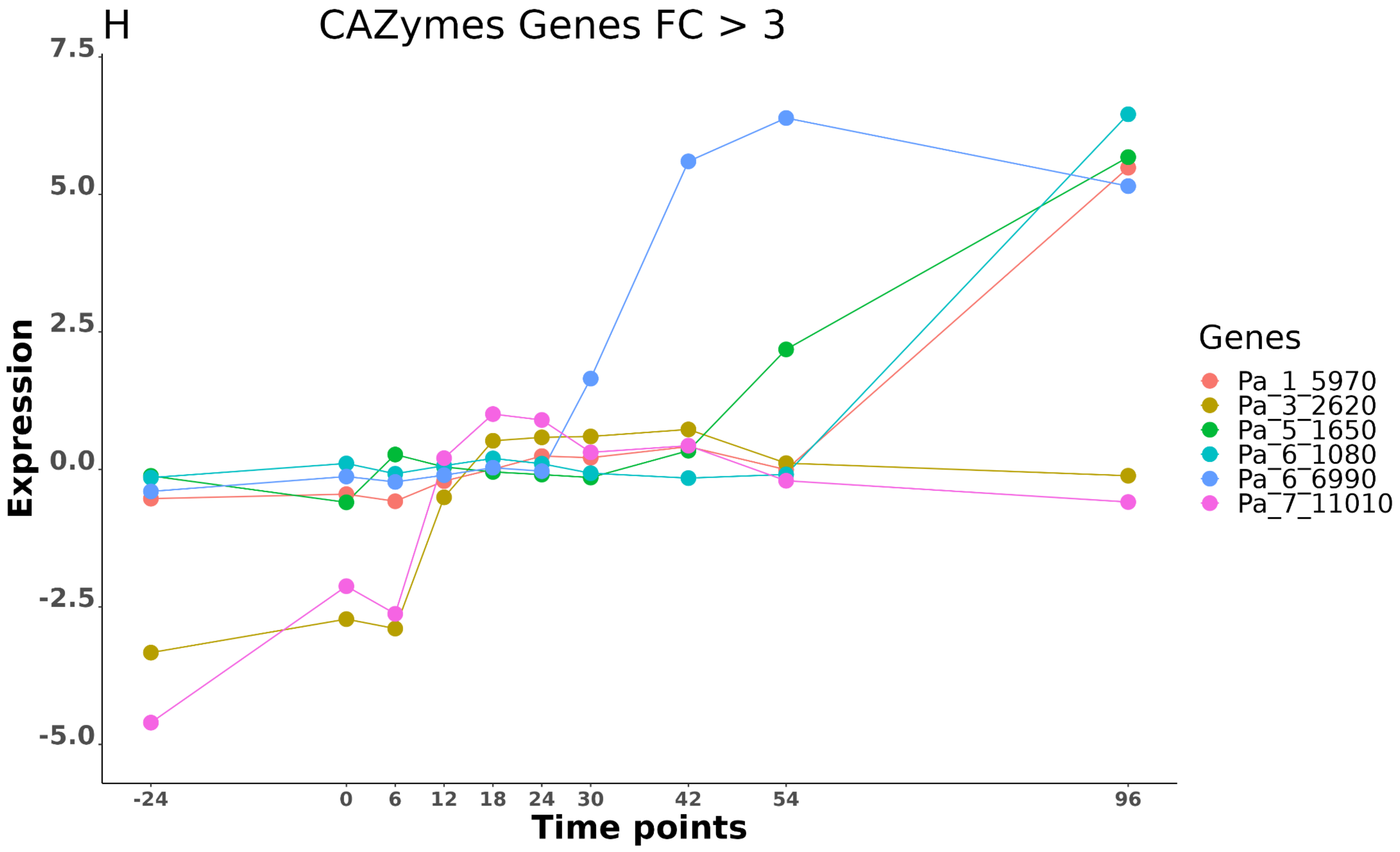

H
CAZymes Genes FC > 3
7.5
5.0
2.5
0.0
-2.5
-5.0
Expression
-24
0
6
12
18
24
30
42
54
96
Time points
Genes
Pa_1_5970
Pa_3_2620
Pa_5_1650
Pa_6_1080
Pa_6_6990
Pa_7_11010

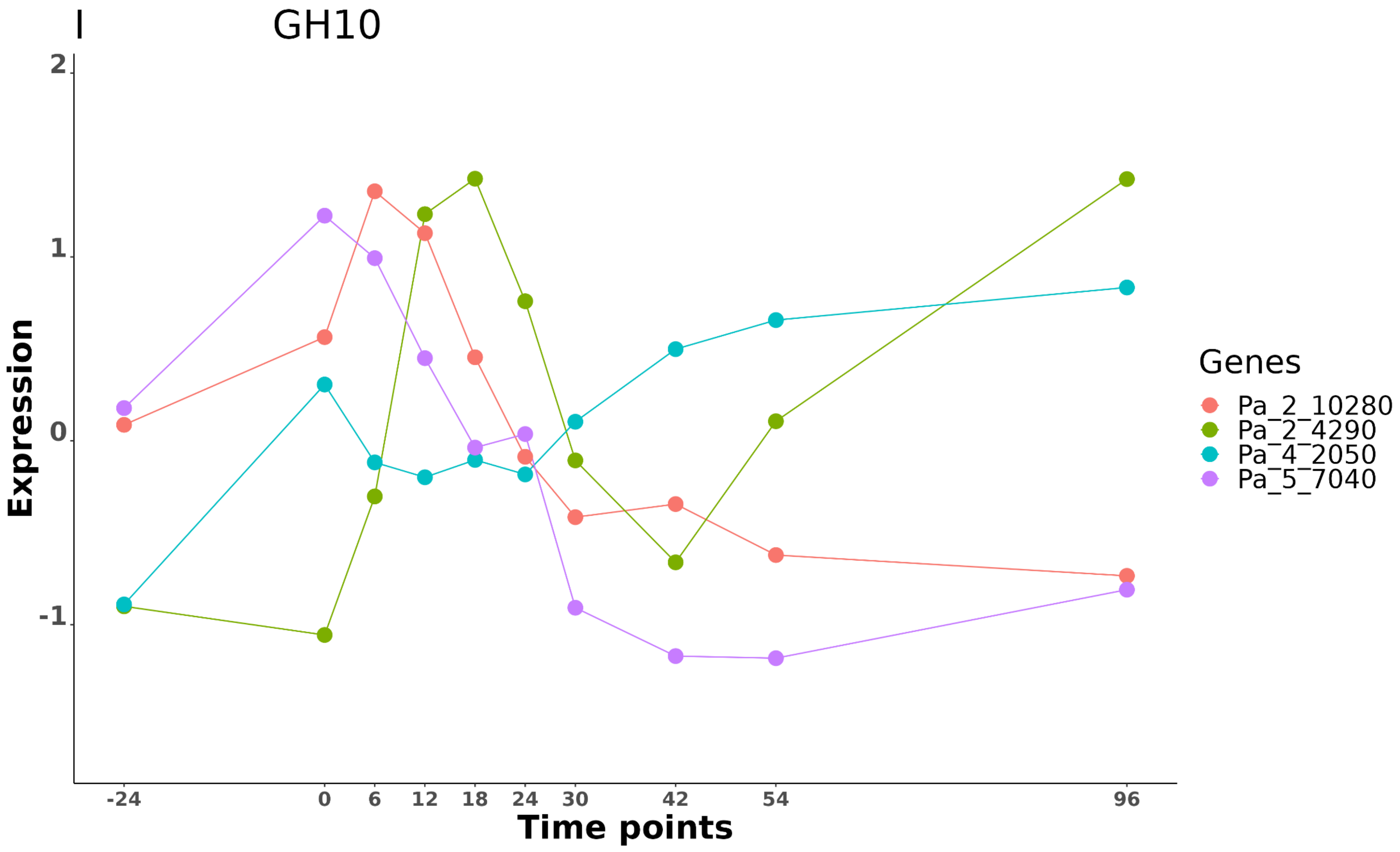
I
GH10
Expression
Time points
2
1
0
-1
-24
0
6
12
18
24
30
42
54
96
Genes
Pa_2_10280
Pa_2_4290
Pa_4_2050
Pa_5_7040

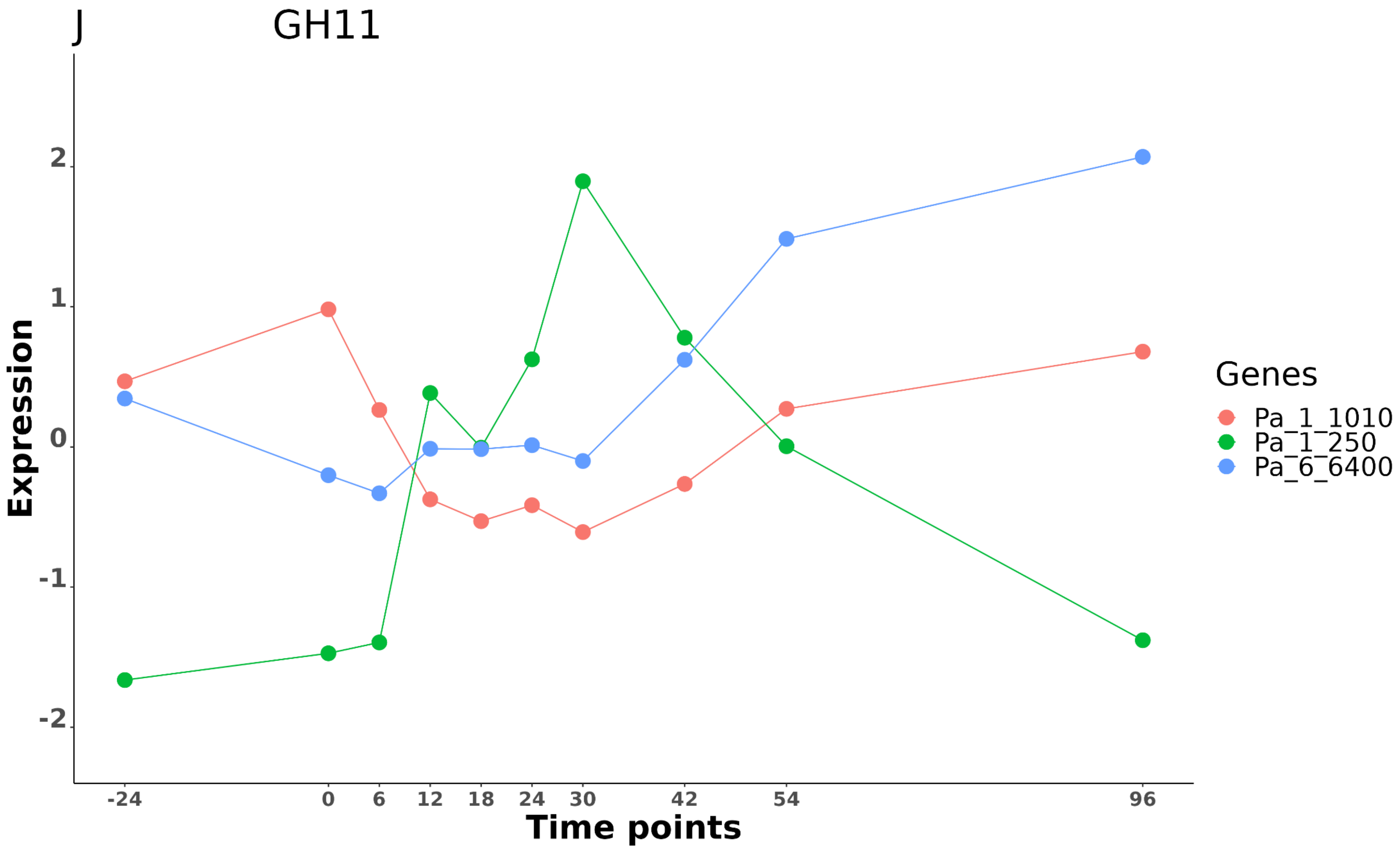

J
GH11
Expression
2
1
0
-1
-2
-24
0
6
12
18
24
30
42
54
96
Time points
Genes
Pa_1_1010
Pa_1_250
Pa_6_6400

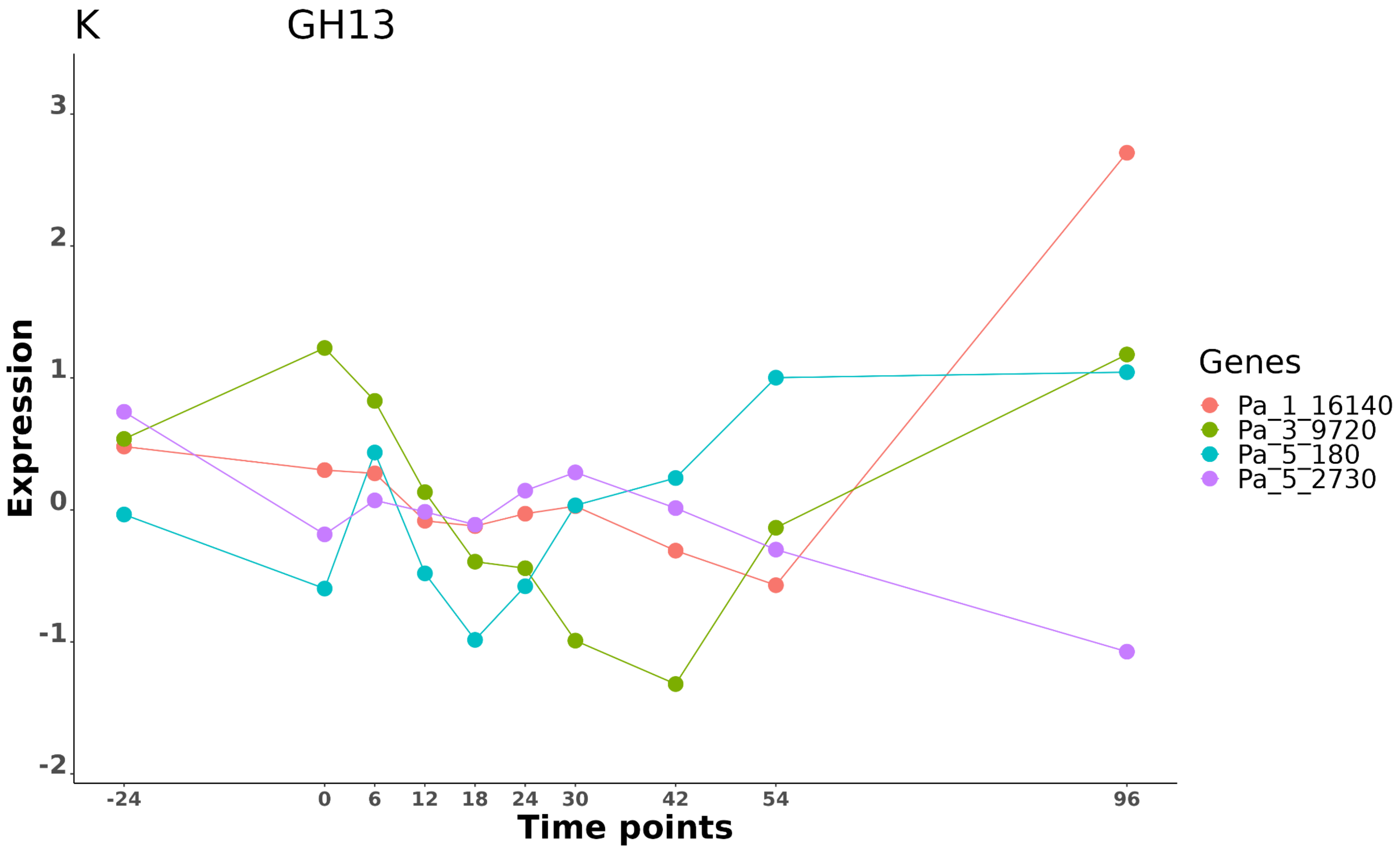

K
GH13
Expression
3
2
1
0
-1
-2
-24
0
6
12
18
24
30
42
54
96
Time points
Genes
Pa_1_16140
Pa_3_9720
Pa_5_180
Pa_5_2730

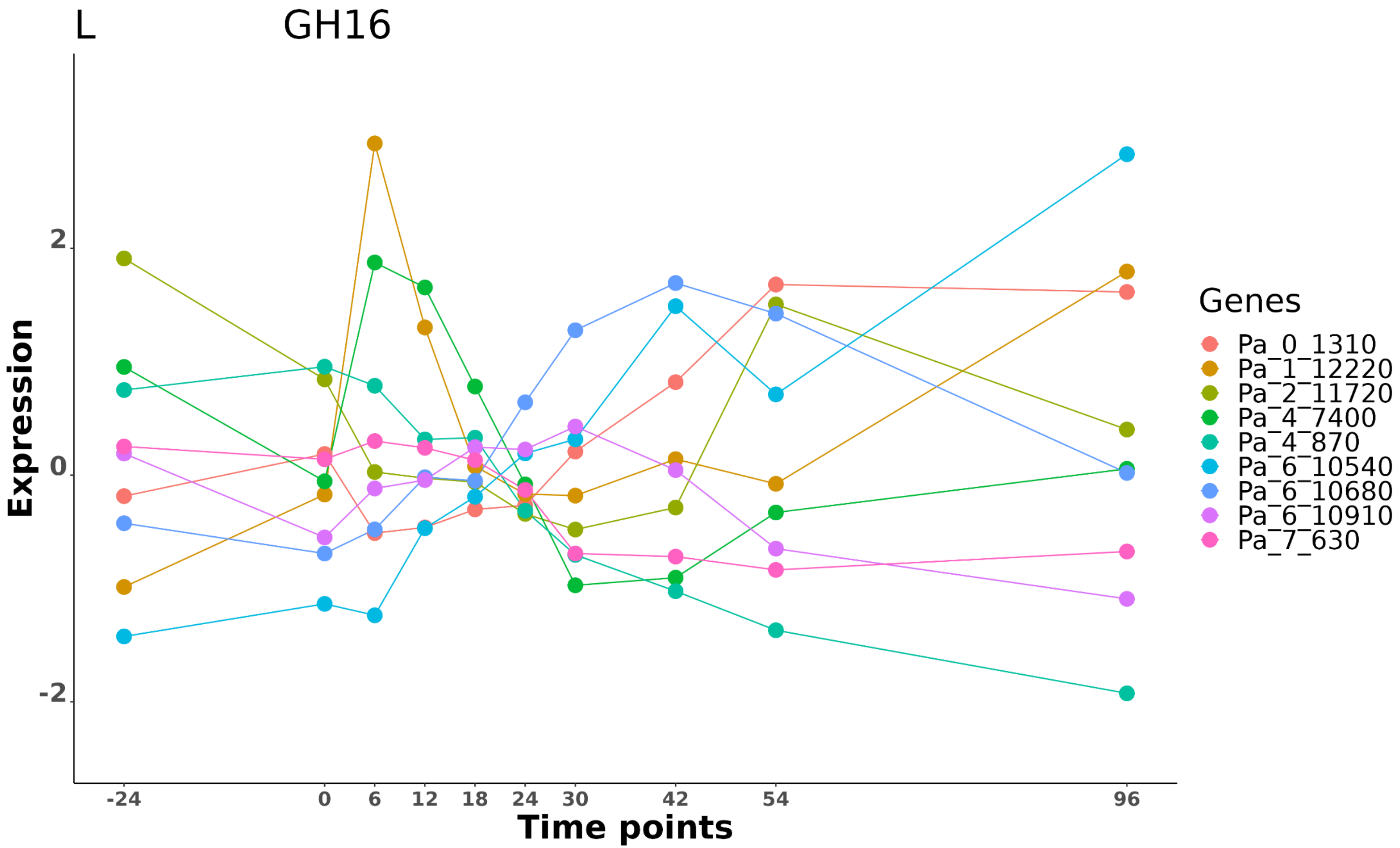

L
GH16
Expression
2
0
-2
-24
0
6
12
18
24
30
42
54
96
Time points
Genes
Pa_0_1310
Pa_1_12220
Pa_2_11720
Pa_4_7400
Pa_4_870
Pa_6_10540
Pa_6_10680
Pa_6_10910
Pa_7_630

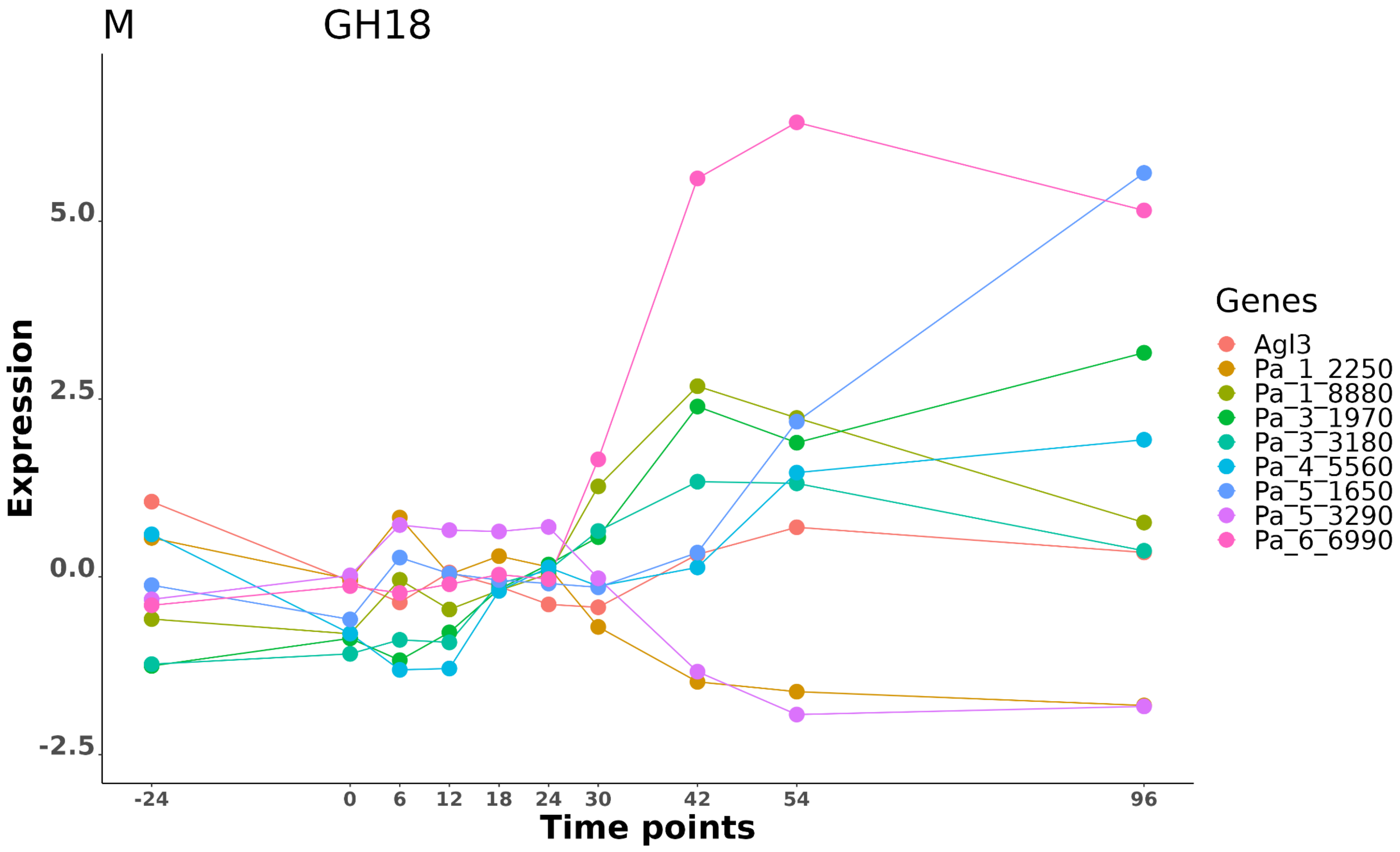

M
GH18
Expression
5.0
2.5
0.0
-2.5
-24
0
6
12
18
24
30
42
54
96
Time points
Genes
Agl3
Pa_1_2250
Pa_1_8880
Pa_3_1970
Pa_3_3180
Pa_4_5560
Pa_5_1650
Pa_5_3290
Pa_6_6990

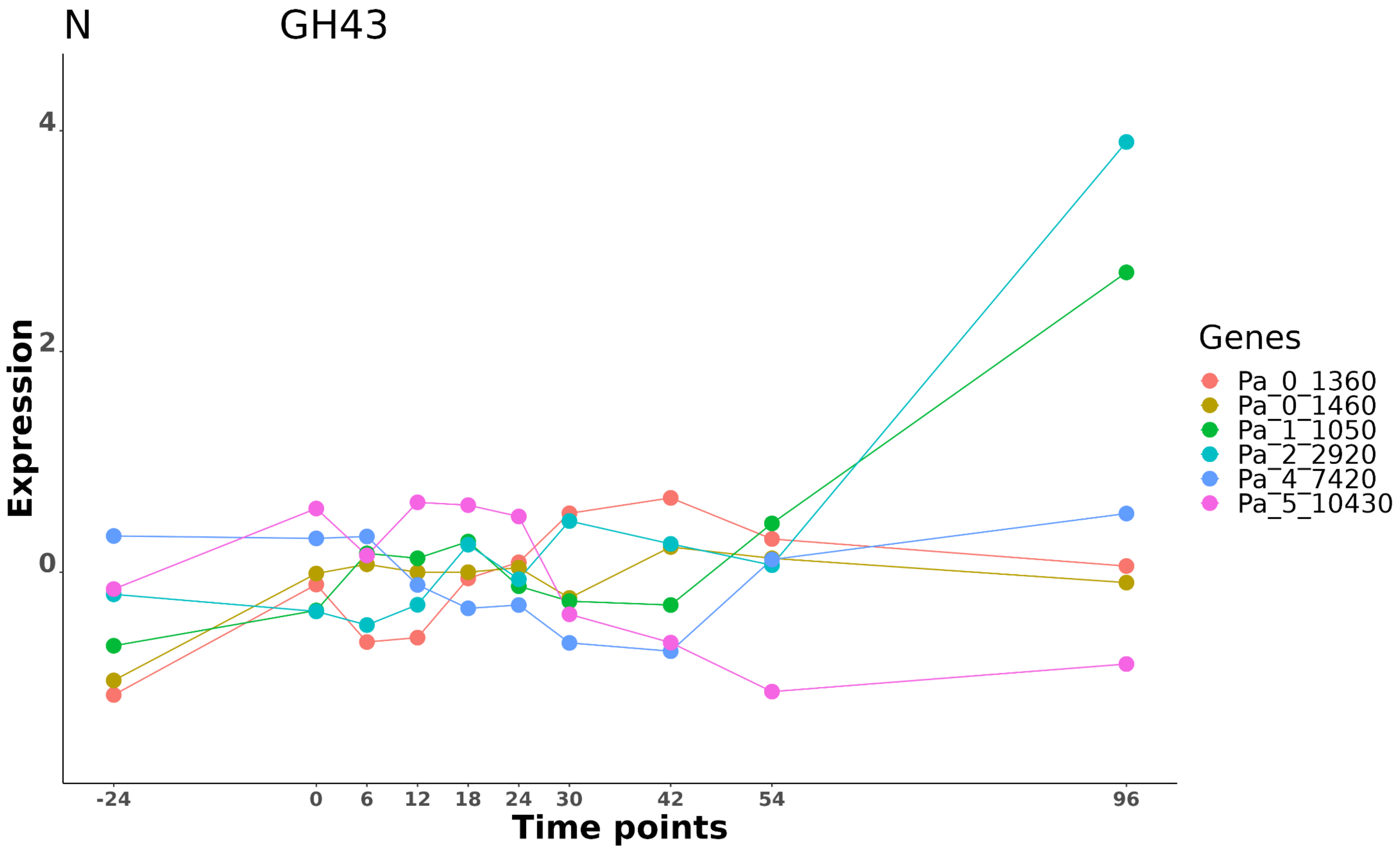

N
GH43
Expression
4
2
0
-24
0
6
12
18
24
30
42
54
96
Time points
Genes
Pa_0_1360
Pa_0_1460
Pa_1_1050
Pa_2_2920
Pa_4_7420
Pa_5_10430

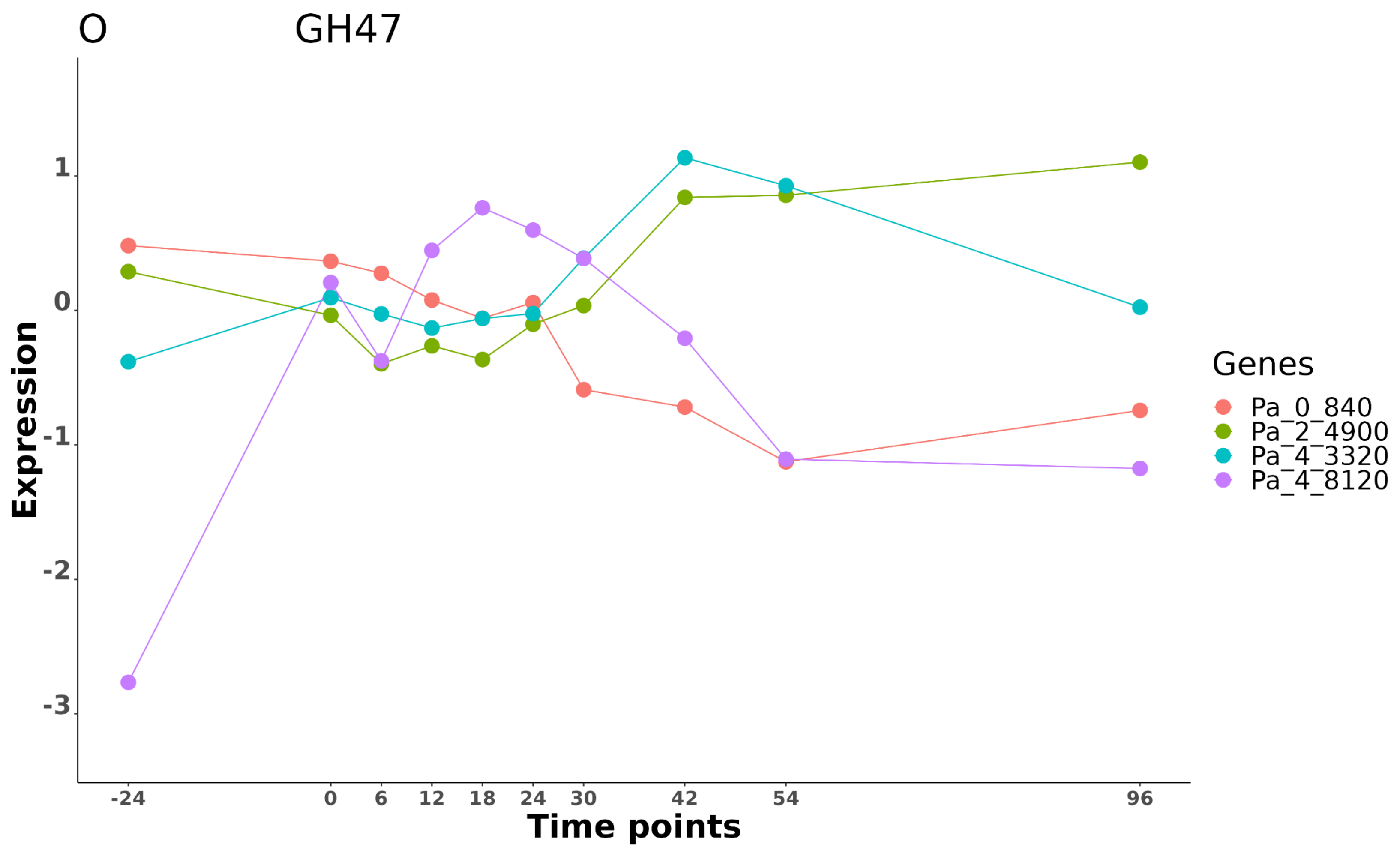

O
GH47
Expression
1
0
-1
-2
-3
-24
0
6
12
18
24
30
42
54
96
Time points
Genes
Pa_0_840
Pa_2_4900
Pa_4_3320
Pa_4_8120

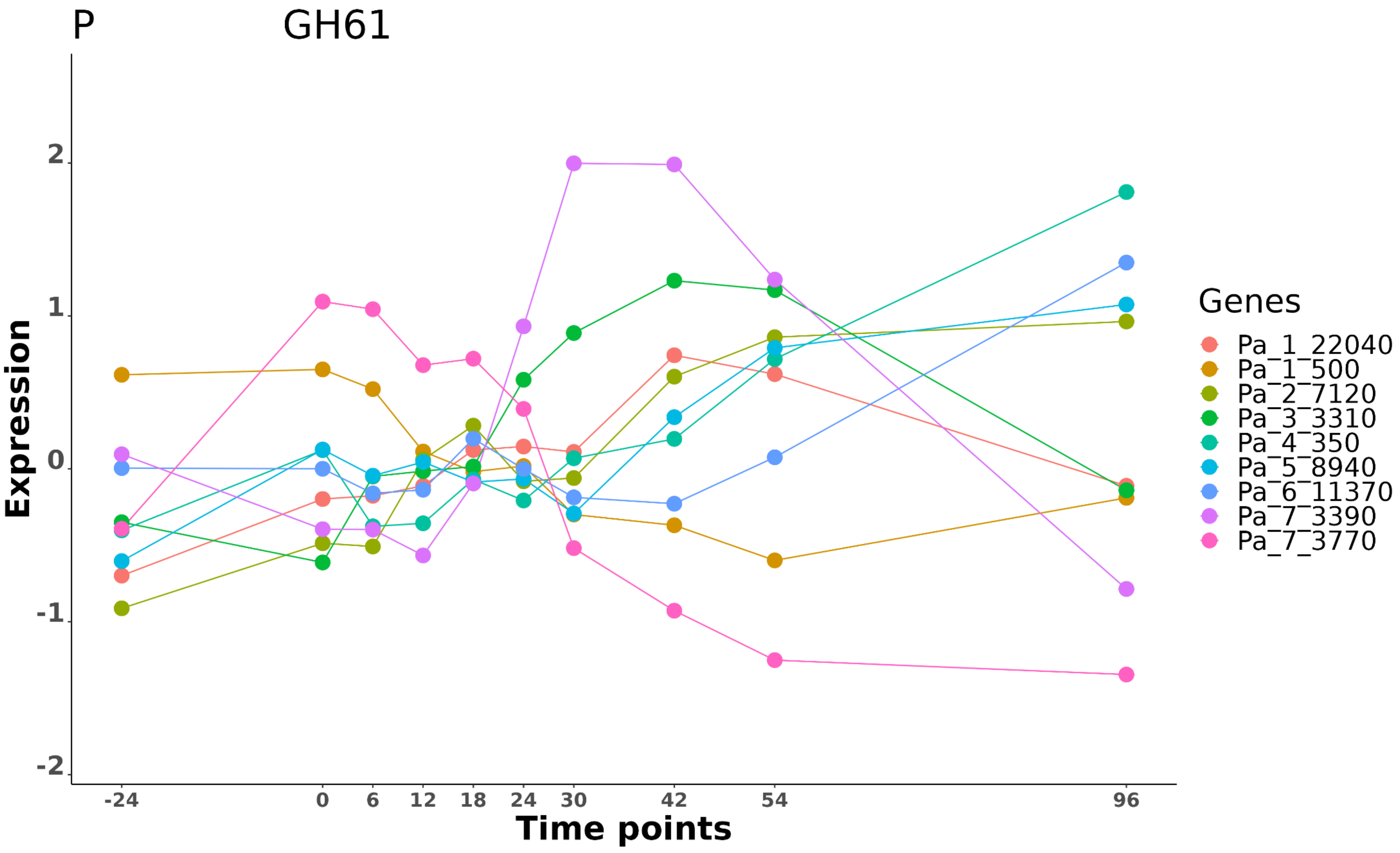
P
GH61
Expression
2
1
0
-1
-2
-24
0
6
12
18
24
30
42
54
96
Time points
Genes
Pa_1_22040
Pa_1_500
Pa_2_7120
Pa_3_3310
Pa_4_350
Pa_5_8940
Pa_6_11370
Pa_7_3390
Pa_7_3770

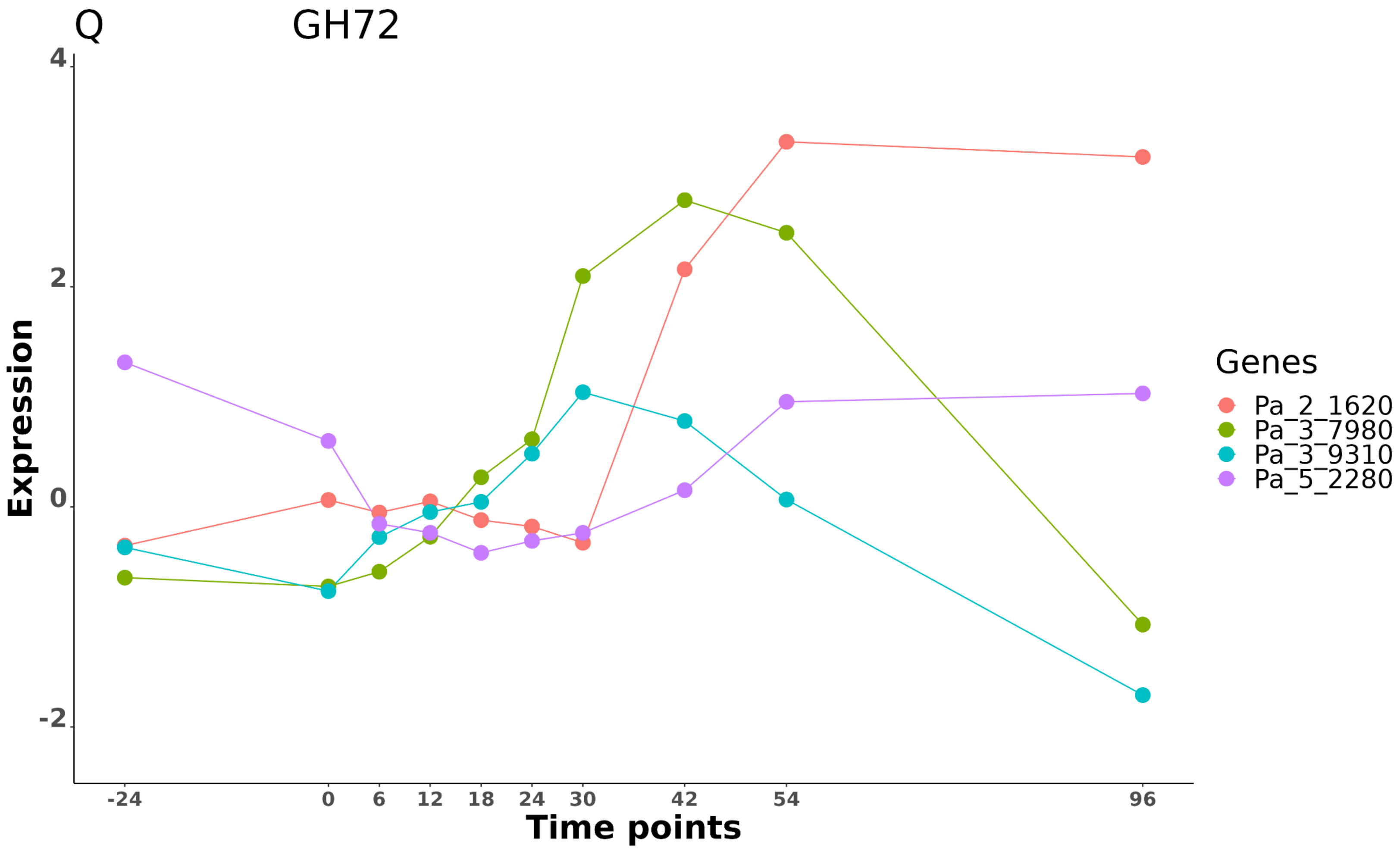
Q
GH72
Expression
4
2
0
-2
Time points
-24
0
6
12
18
24
30
42
54
96
Genes
Pa_2_1620
Pa_3_7980
Pa_3_9310
Pa_5_2280

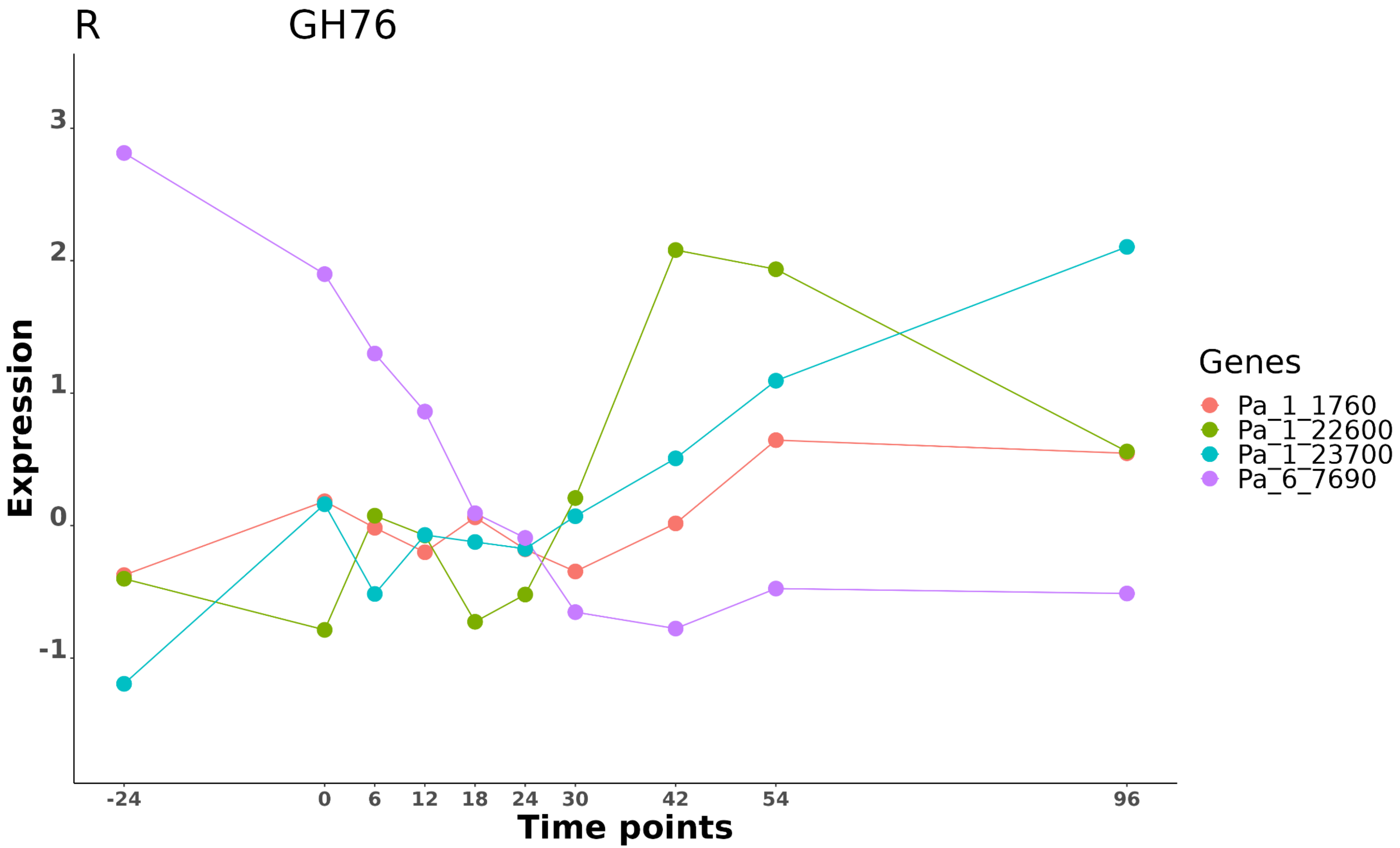

R
GH76
Expression
3
2
1
0
-1
-24
0
6
12
18
24
30
42
54
96
Time points
Genes
Pa_1_1760
Pa_1_22600
Pa_1_23700
Pa_6_7690

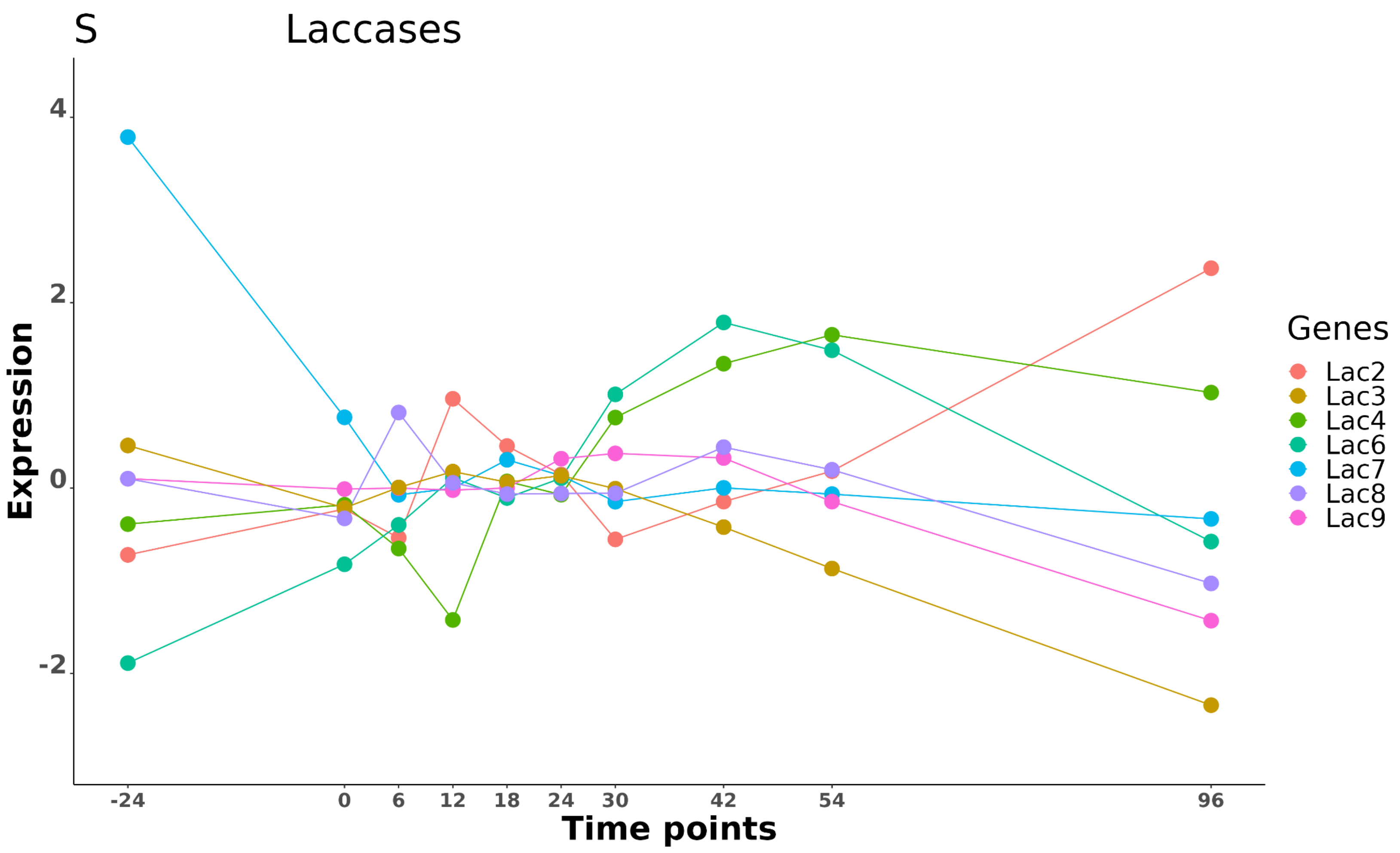
S
Laccases
Expression
4
2
0
-2
-24
0
6
12
18
24
30
42
54
96
Time points
Genes
Lac2
Lac3
Lac4
Lac6
Lac7
Lac8
Lac9

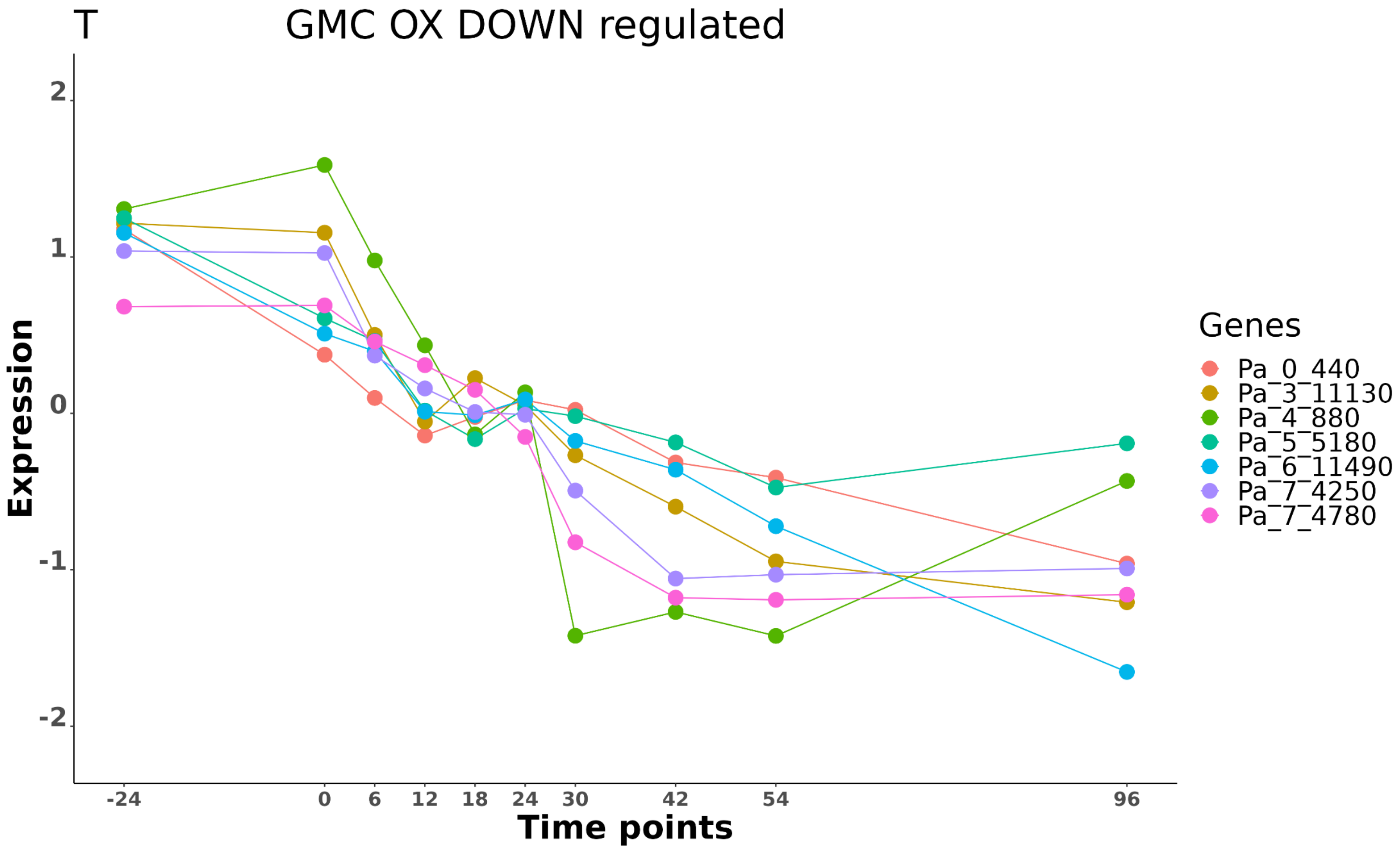
T
GMC OX DOWN regulated
Expression
2
1
0
-1
-2
-24
0
6
12
18
24
30
42
54
96
Time points
Genes
Pa_0_440
Pa_3_11130
Pa_4_880
Pa_5_5180
Pa_6_11490
Pa_7_4250
Pa_7_4780

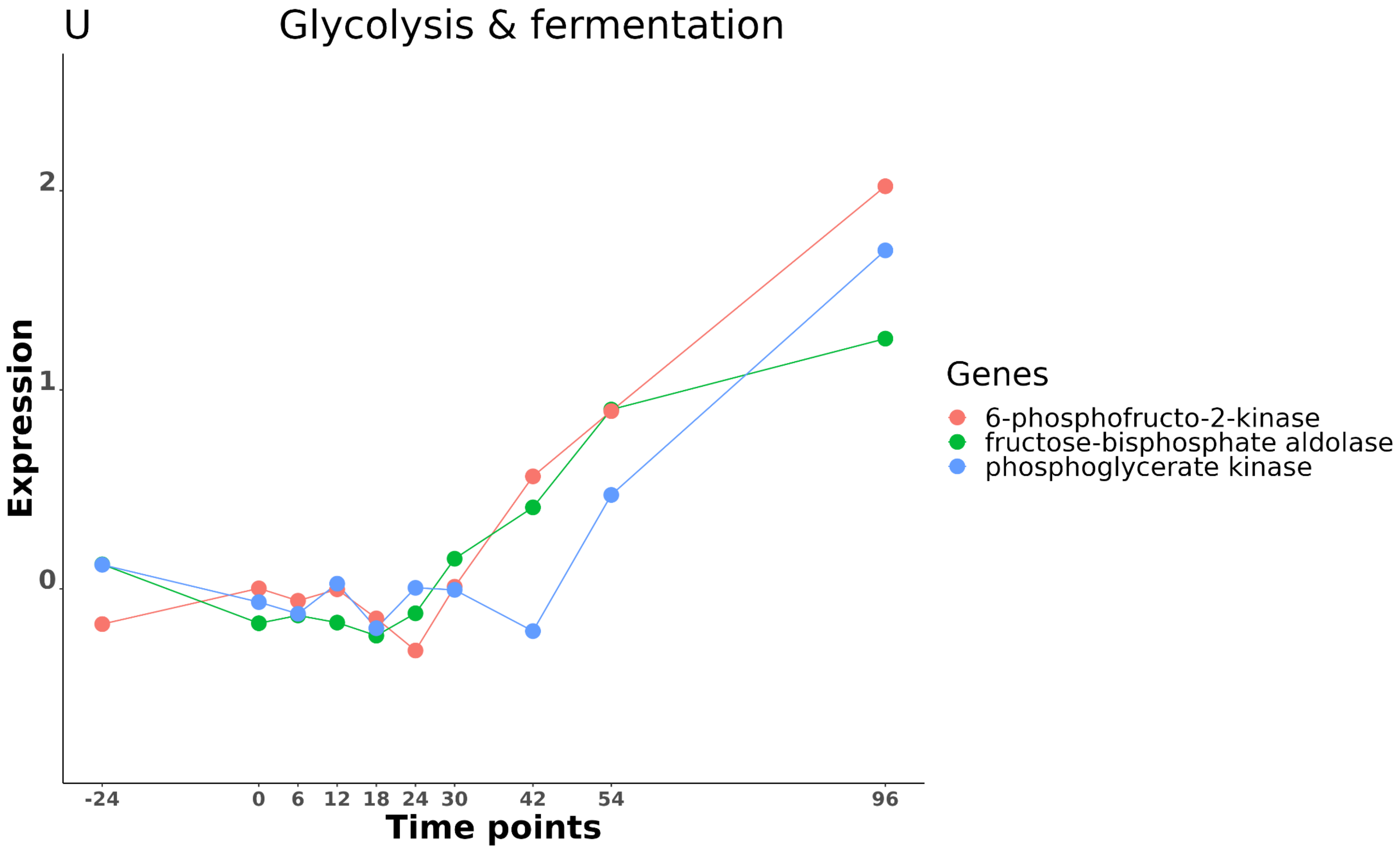
U
Glycolysis & fermentation
Expression
2
1
0
-24
0
6
12
18
24
30
42
54
96
Time points
Genes
6-phosphofructo-2-kinase
fructose-bisphosphate aldolase
phosphoglycerate kinase

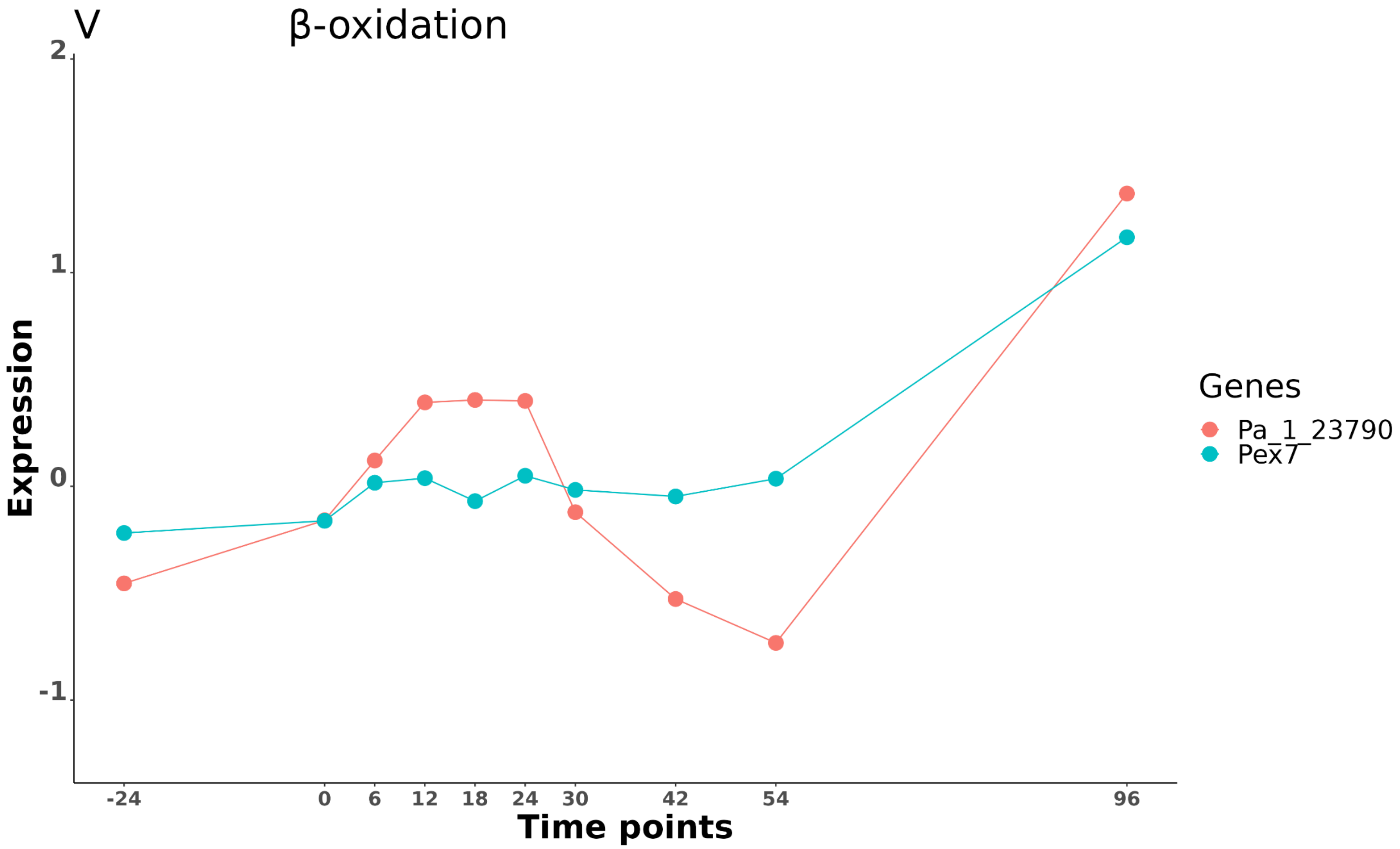
V
β-oxidation
Expression
2
1
0
-1
-24
0
6
12
18
24
30
42
54
96
Time points
Genes
Pa_1_23790
Pex7

W Expression profile of differentially expressed genes of the 36 secondary metabolite clusters present in *P. anserina* genome

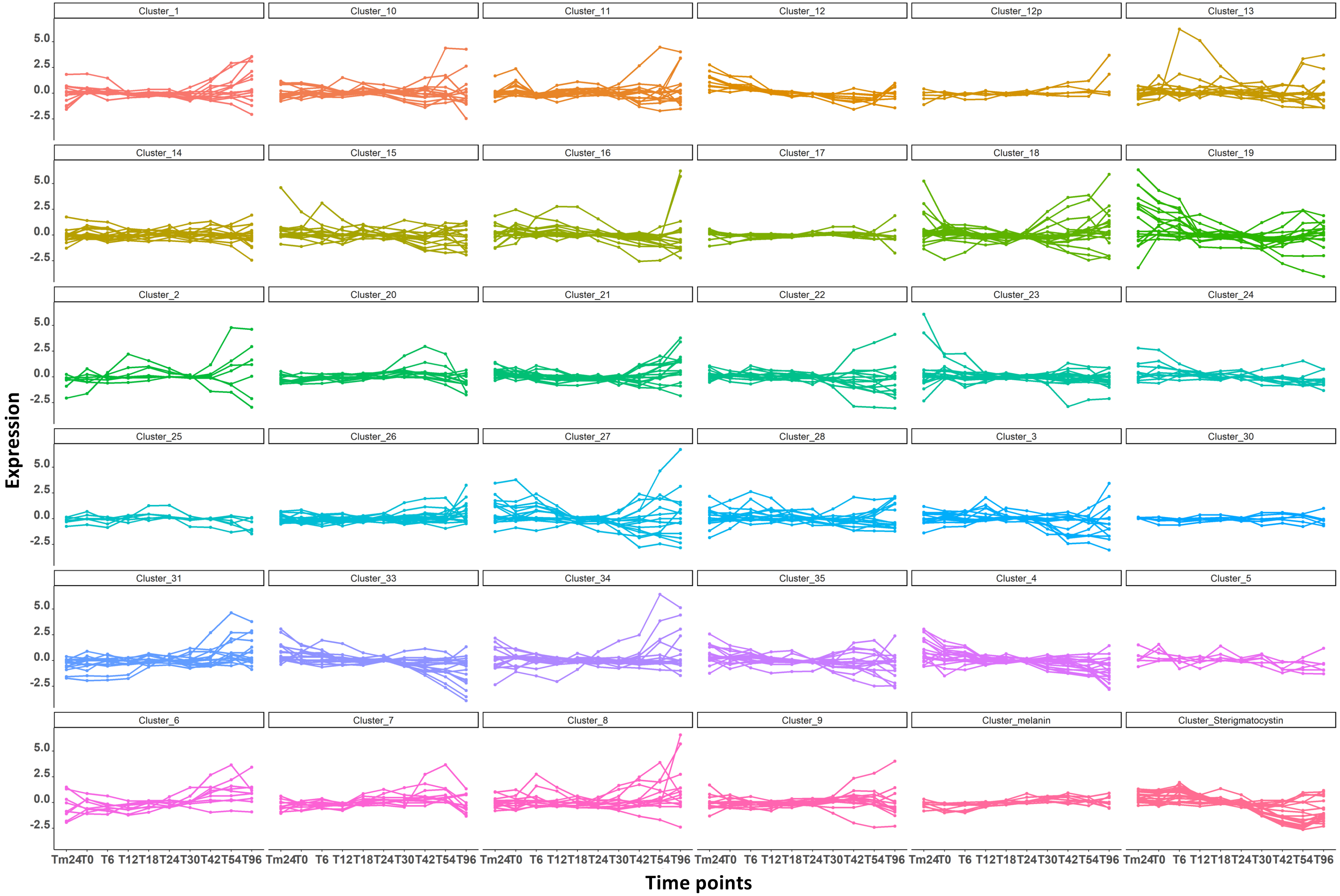

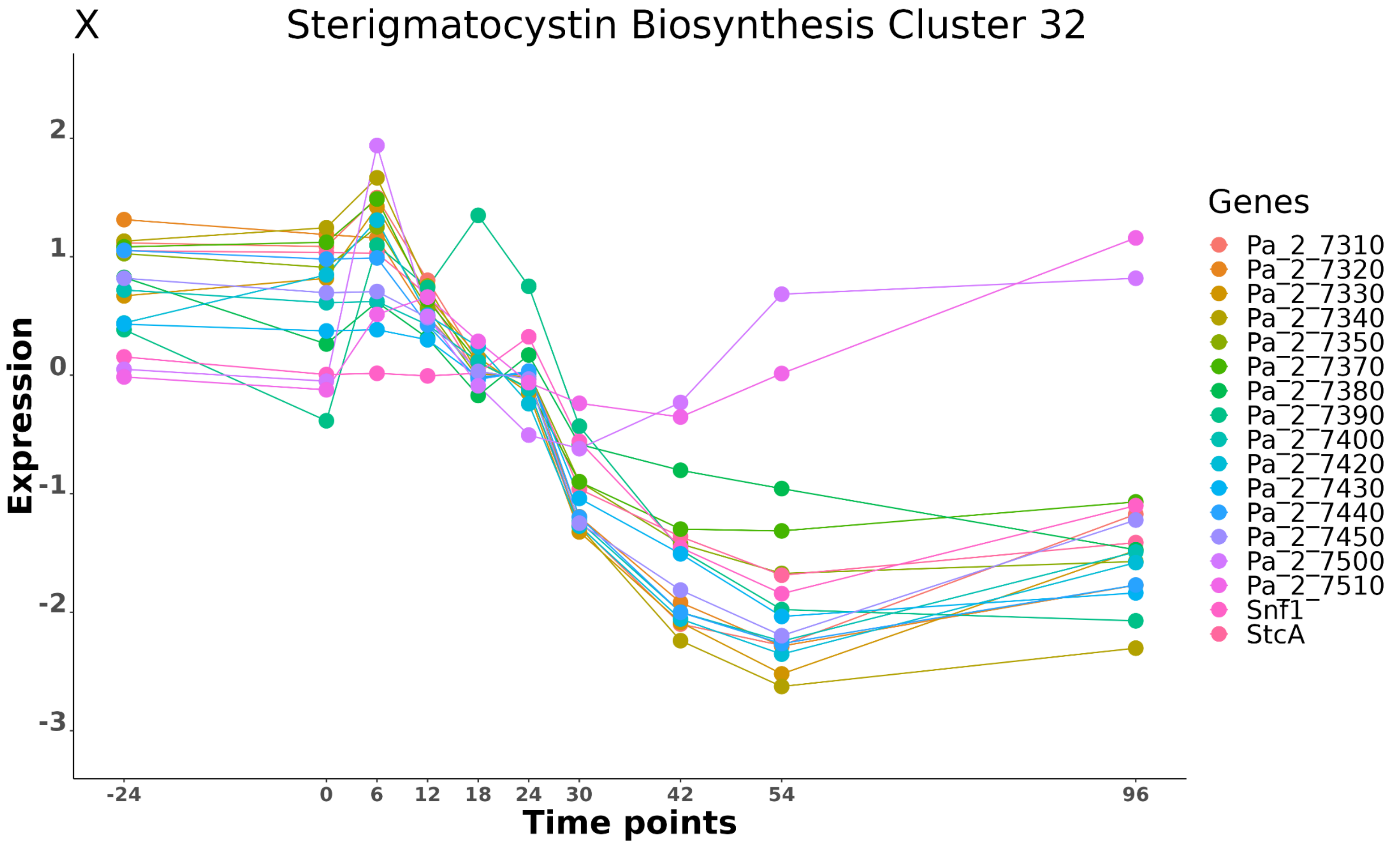

X
Sterigmatocystin Biosynthesis Cluster 32
Expression
2
1
0
-1
-2
-3
-24
0
6
12
18
24
30
42
54
96
Time points
Genes
Pa_2_7310
Pa_2_7320
Pa_2_7330
Pa_2_7340
Pa_2_7350
Pa_2_7370
Pa_2_7380
Pa_2_7390
Pa_2_7400
Pa_2_7420
Pa_2_7430
Pa_2_7440
Pa_2_7450
Pa_2_7500
Pa_2_7510
Snf1
StcA

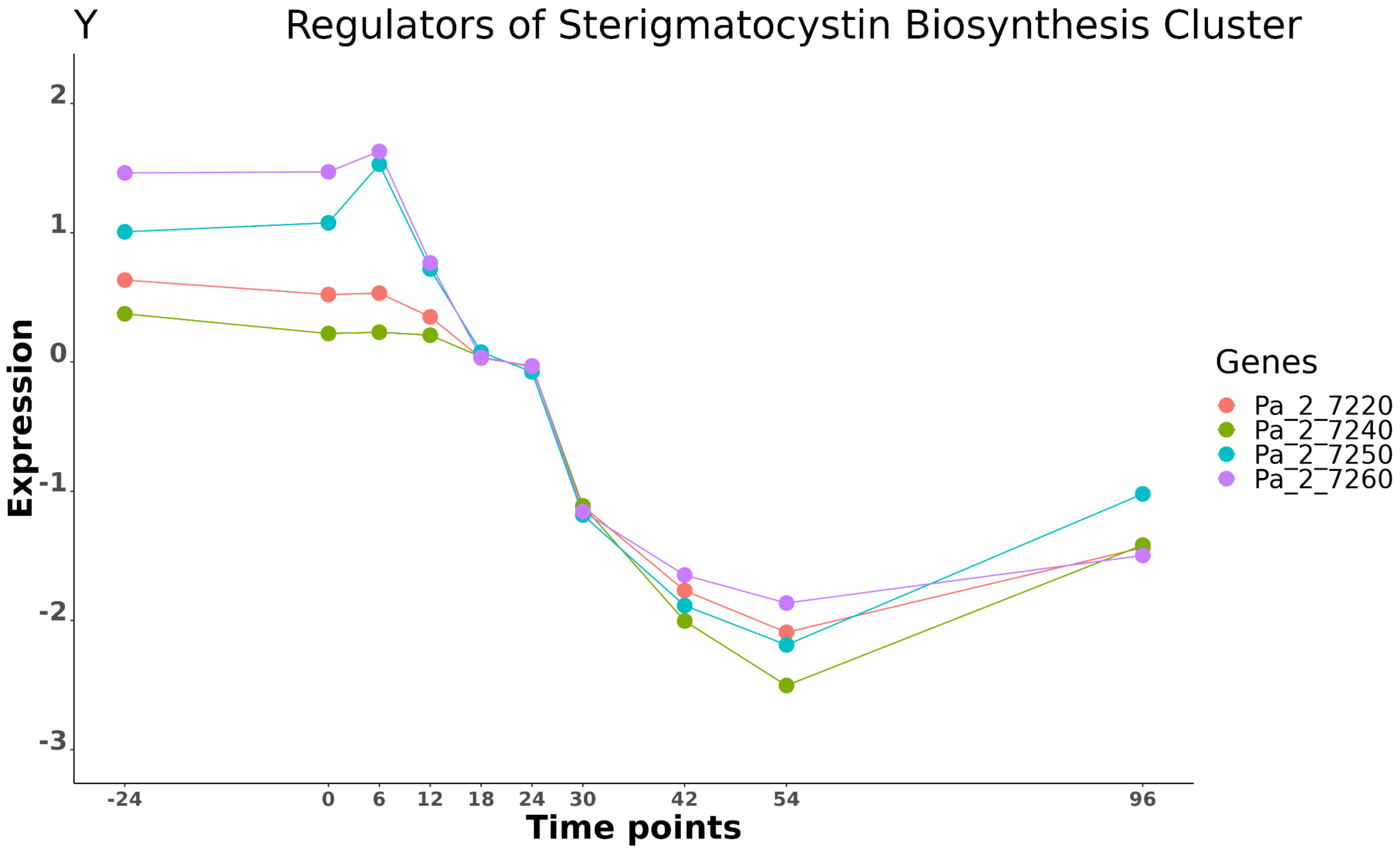
Y
Regulators of Sterigmatocystin Biosynthesis Cluster
Expression
2
1
0
-1
-2
-3
-24
0
6
12
18
24
30
42
54
96
Time points
Genes
Pa_2_7220
Pa_2_7240
Pa_2_7250
Pa_2_7260

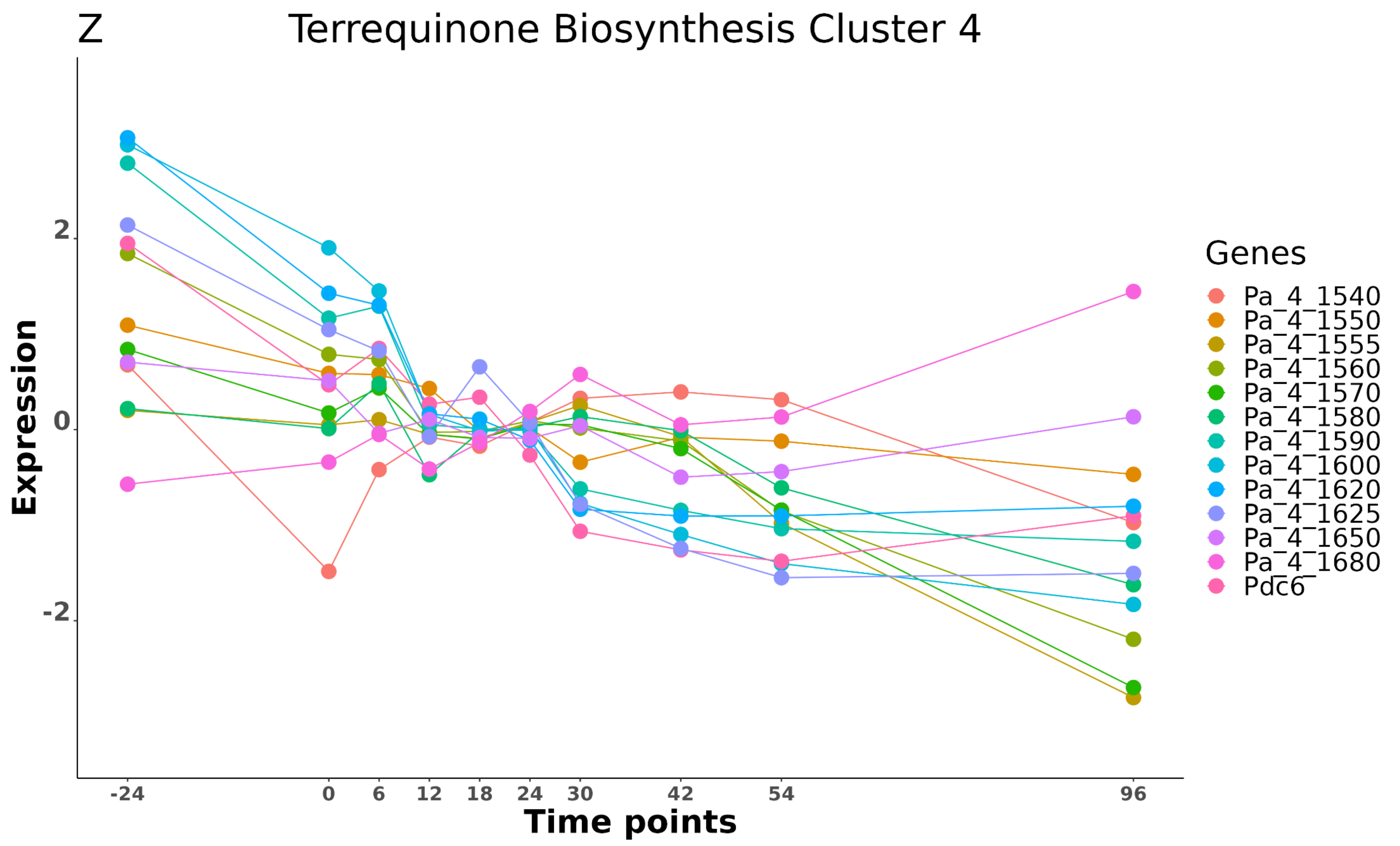

Z
Terrequinone Biosynthesis Cluster 4
Expression
2
0
-2
-24
0
6
12
18
24
30
42
54
96
Time points
Genes
Pa_4_1540
Pa_4_1550
Pa_4_1555
Pa_4_1560
Pa_4_1570
Pa_4_1580
Pa_4_1590
Pa_4_1600
Pa_4_1620
Pa_4_1625
Pa_4_1650
Pa_4_1680
Pdc6

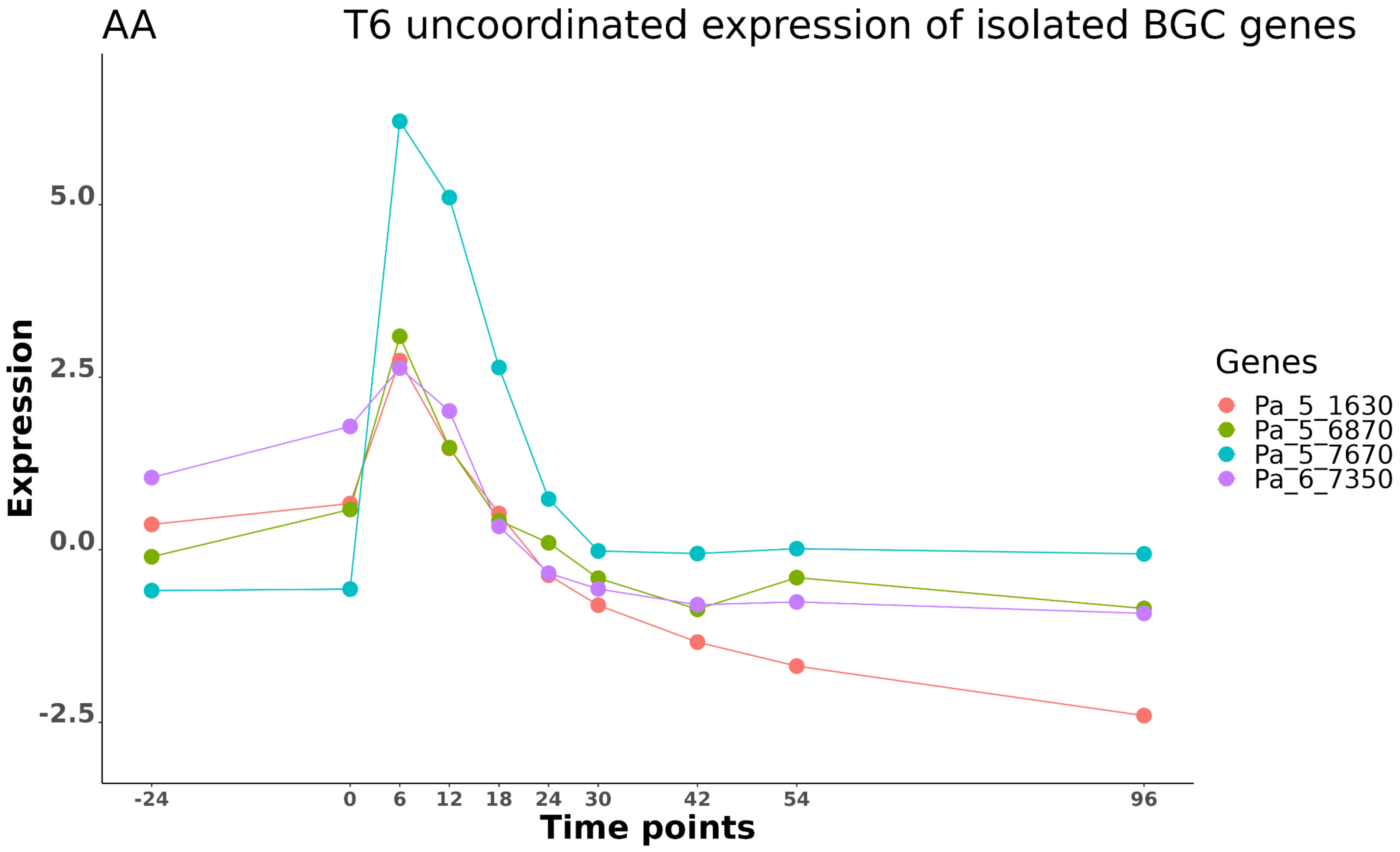

AA
T6 uncoordinated expression of isolated BGC genes
Expression
5.0
2.5
0.0
-2.5
-24
0
6
12
18
24
30
42
54
96
Time points
Genes
Pa_5_1630
Pa_5_6870
Pa_5_7670
Pa_6_7350

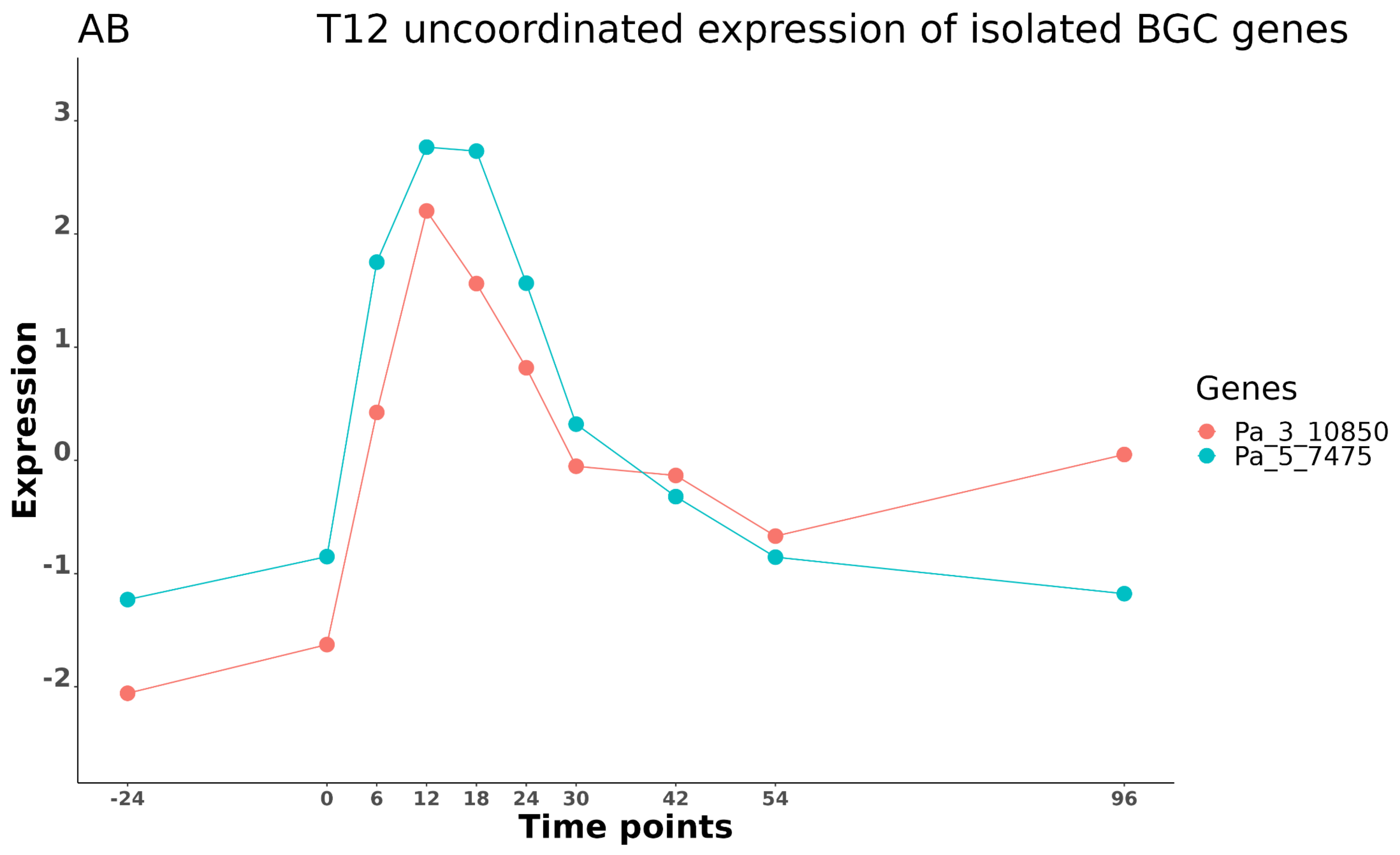
AB
T12 uncoordinated expression of isolated BGC genes
Expression
3
2
1
0
-1
-2
Time points
-24
0
6
12
18
24
30
42
54
96
Genes
Pa_3_10850
Pa_5_7475

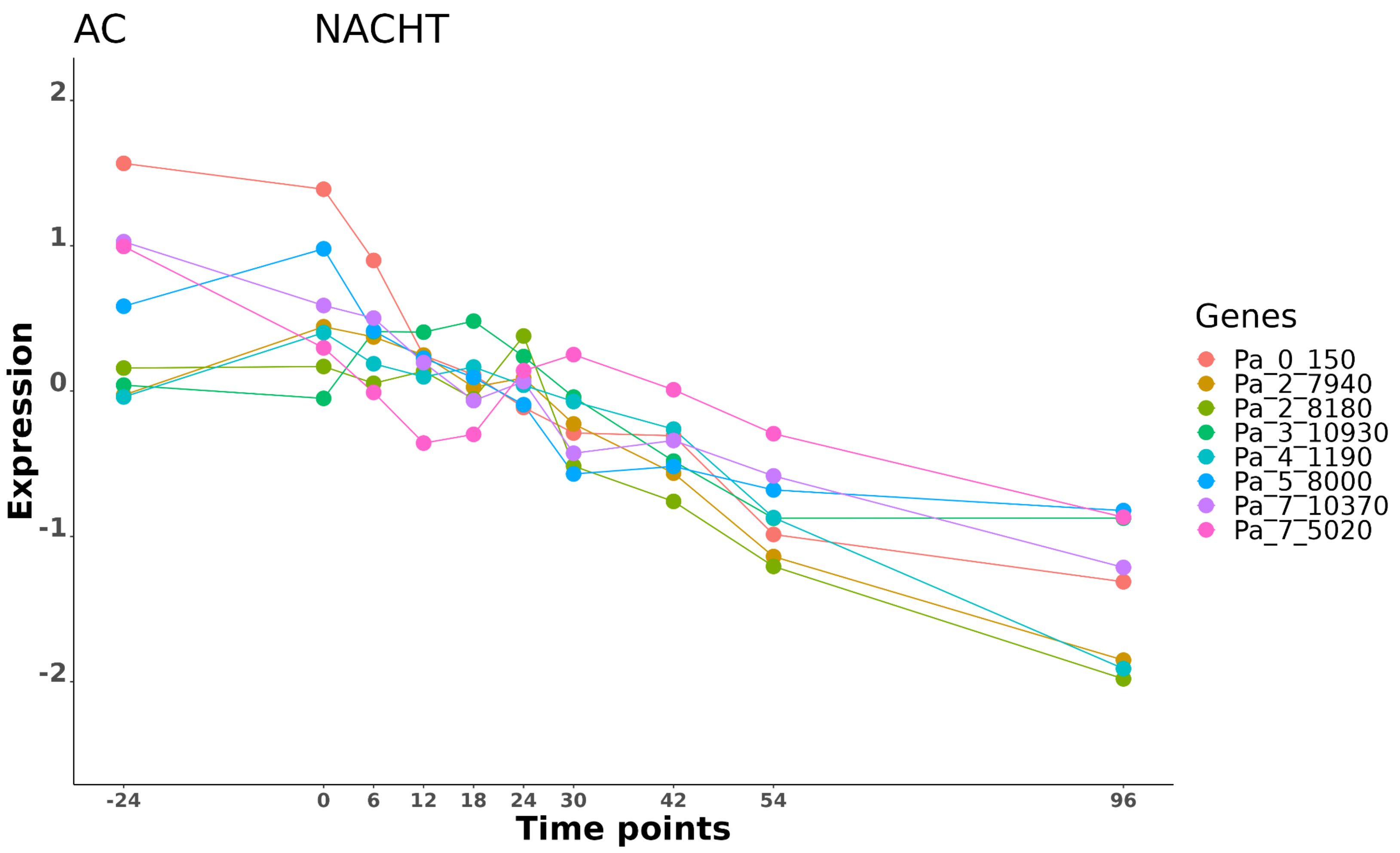

AC
NACHT
Expression
2
1
0
-1
-2
-24
0
6
12
18
24
30
42
54
96
Time points
Genes
Pa_0_150
Pa_2_7940
Pa_2_8180
Pa_3_10930
Pa_4_1190
Pa_5_8000
Pa_7_10370
Pa_7_5020

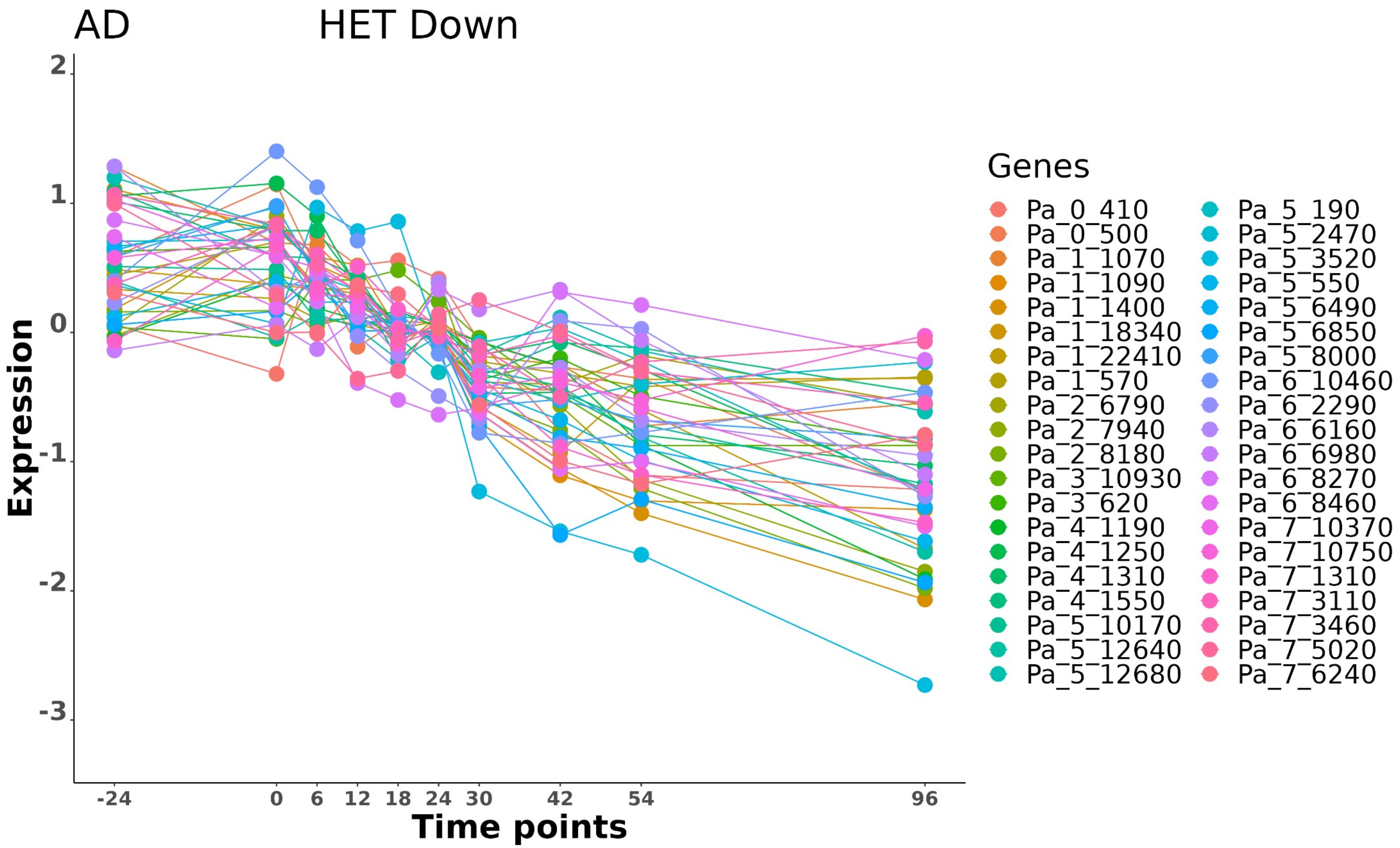

AD
HET Down
Expression
Time points
2
1
0
-1
-2
-3
-24
0
6
12
18
24
30
42
54
96
Genes
Pa_0_410
Pa_0_500
Pa_1_1070
Pa_1_1090
Pa_1_1400
Pa_1_18340
Pa_1_22410
Pa_1_570
Pa_2_6790
Pa_2_7940
Pa_2_8180
Pa_3_10930
Pa_3_620
Pa_4_1190
Pa_4_1250
Pa_4_1310
Pa_4_1550
Pa_5_10170
Pa_5_12640
Pa_5_12680
Pa_5_190
Pa_5_2470
Pa_5_3520
Pa_5_550
Pa_5_6490
Pa_5_6850
Pa_5_8000
Pa_6_10460
Pa_6_2290
Pa_6_6160
Pa_6_6980
Pa_6_8270
Pa_6_8460
Pa_7_10370
Pa_7_10750
Pa_7_1310
Pa_7_3110
Pa_7_3460
Pa_7_5020
Pa_7_6240

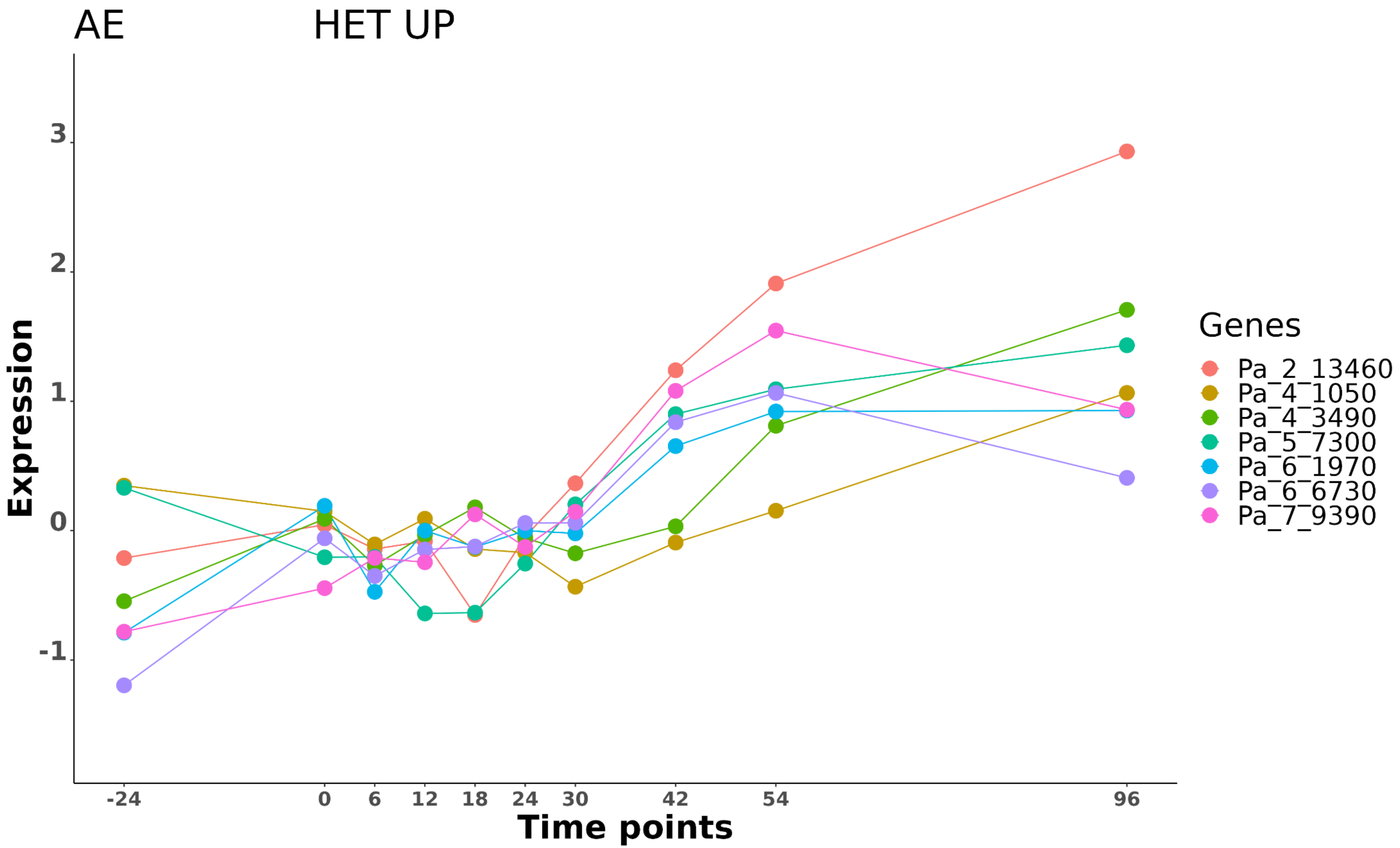
AE
HET UP
Expression
Time points
3
2
1
0
-1
-24
0
6
12
18
24
30
42
54
96
Genes
Pa_2_13460
Pa_4_1050
Pa_4_3490
Pa_5_7300
Pa_6_1970
Pa_6_6730
Pa_7_9390

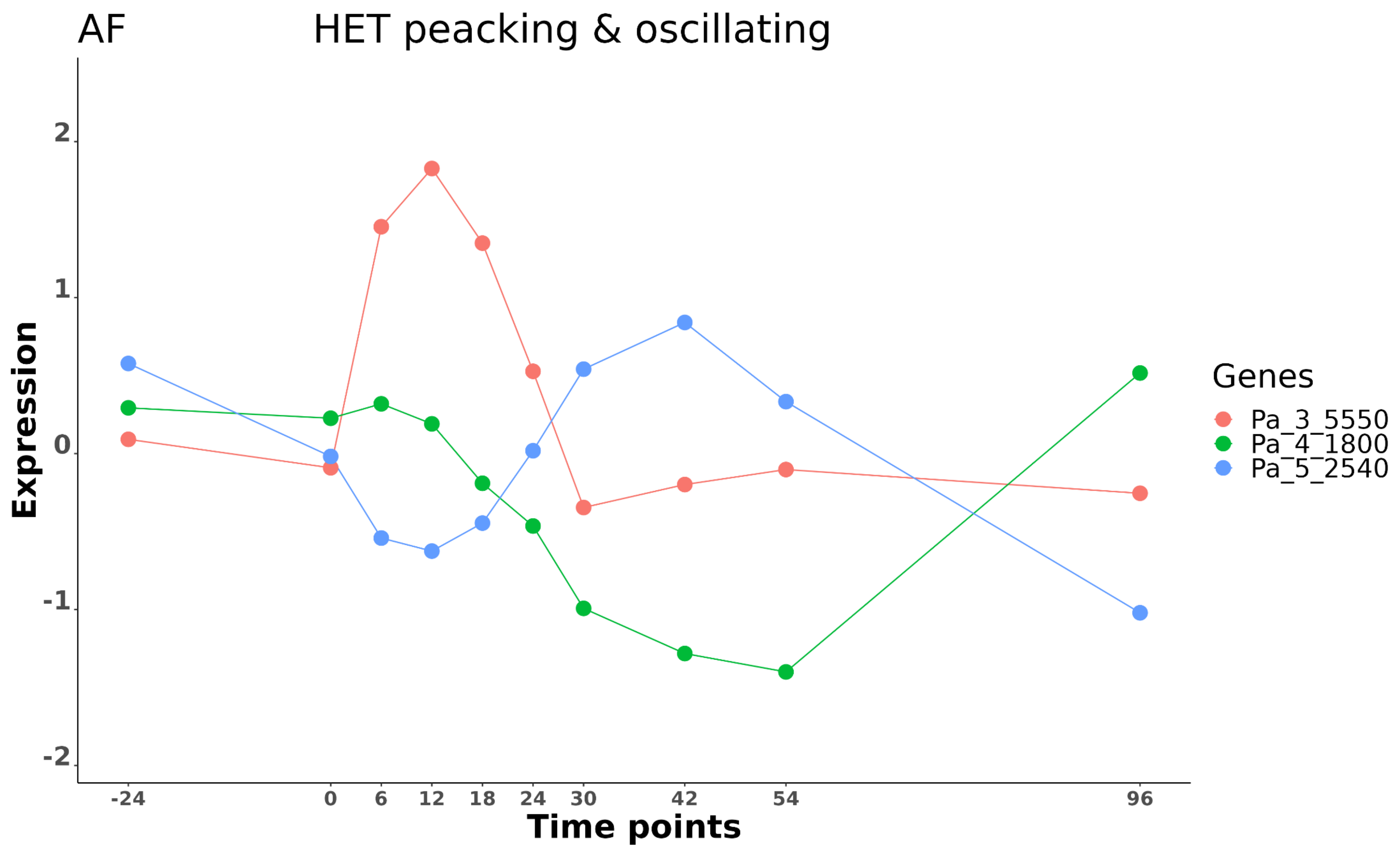

AF
HET peacking & oscillating
Expression
2
1
0
-1
-2
-24
0
6
12
18
24
30
42
54
96
Time points
Genes
Pa_3_5550
Pa_4_1800
Pa_5_2540

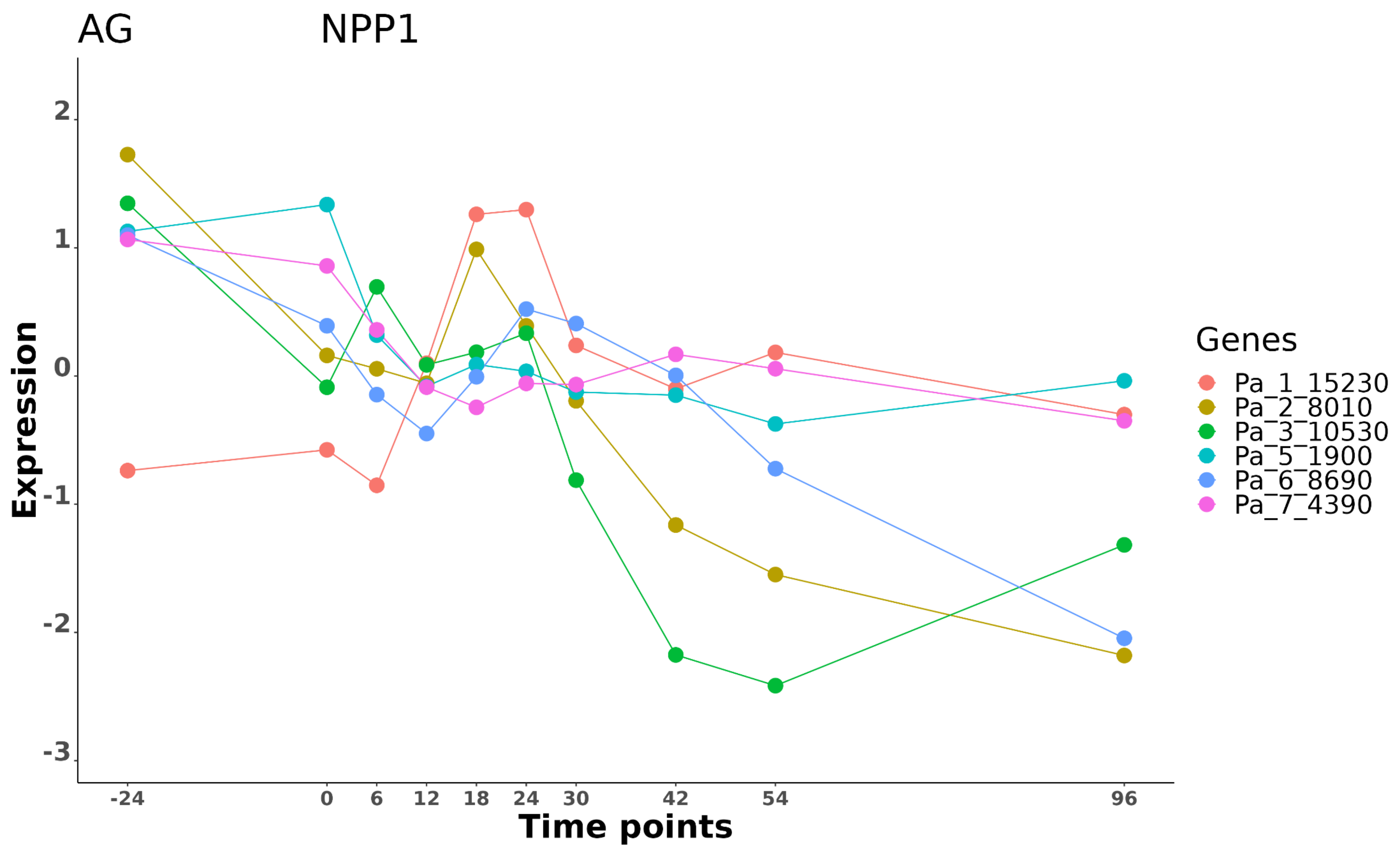
AG
NPP1
Expression
2
1
0
-1
-2
-3
-24
0
6
12
18
24
30
42
54
96
Time points
Genes
Pa_1_15230
Pa_2_8010
Pa_3_10530
Pa_5_1900
Pa_6_8690
Pa_7_4390

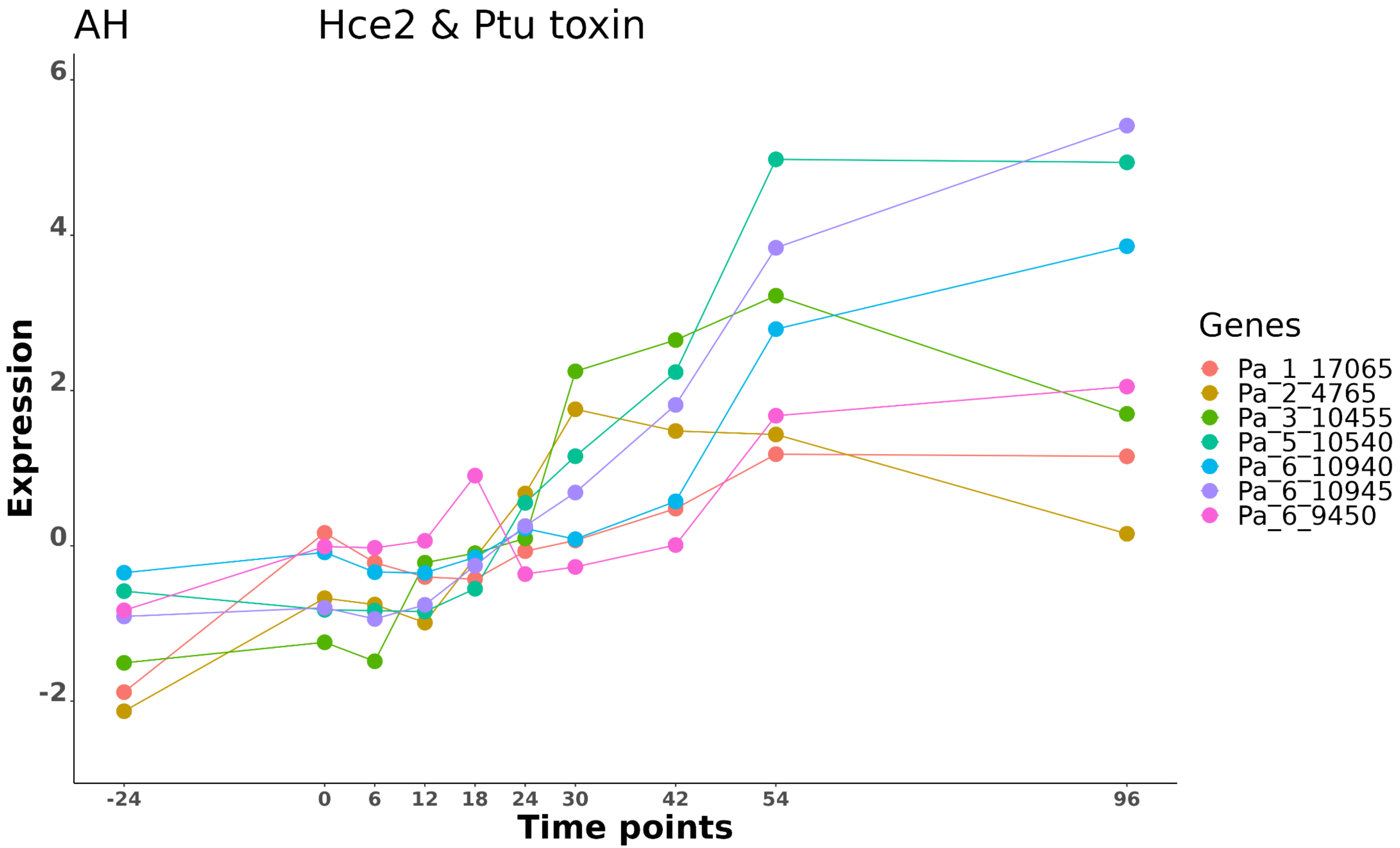
AH
Hce2 & Ptu toxin
Expression
6
4
2
0
-2
-24
0
6
12
18
24
30
42
54
96
Time points
Genes
Pa_1_17065
Pa_2_4765
Pa_3_10455
Pa_5_10540
Pa_6_10940
Pa_6_10945
Pa_6_9450

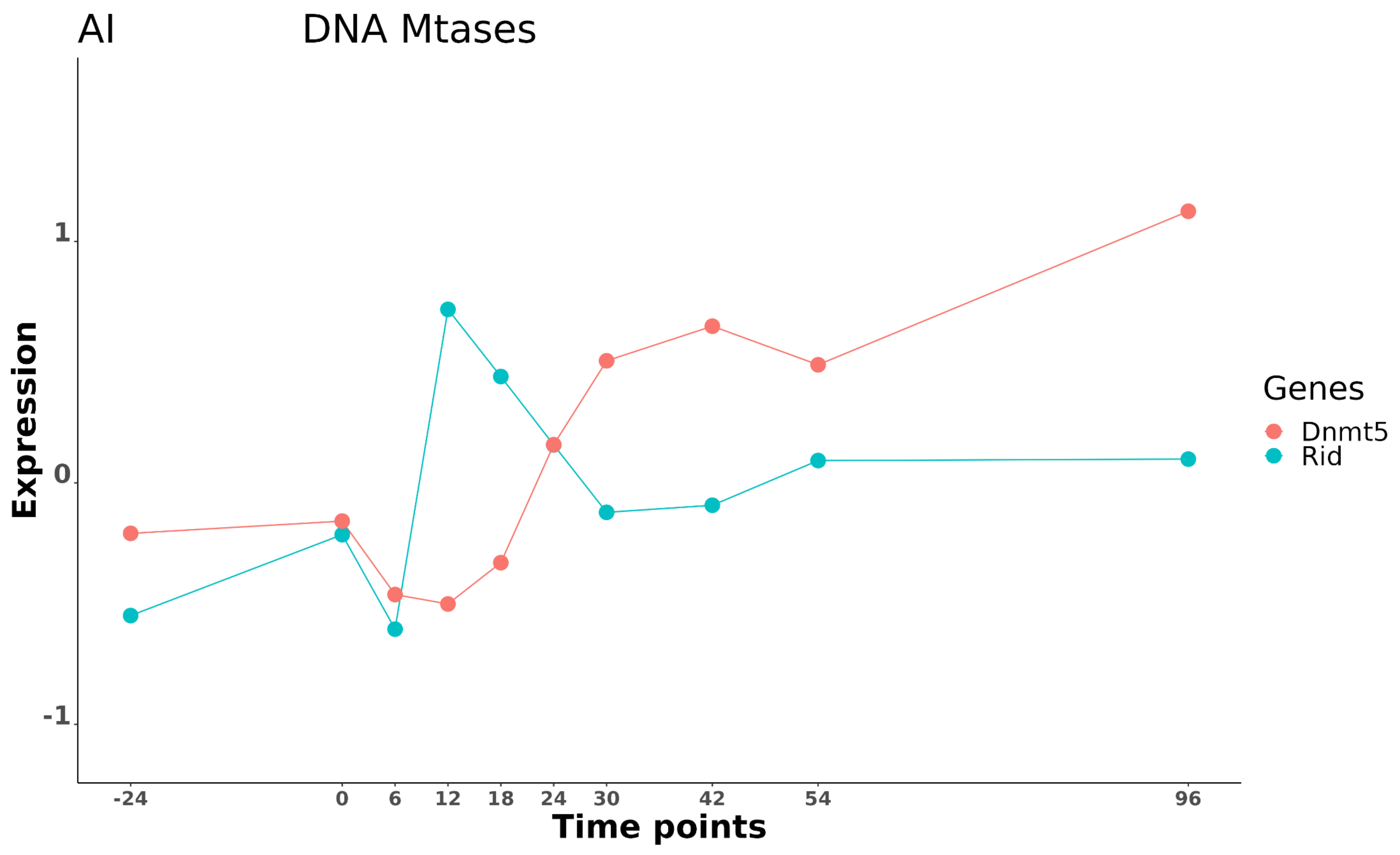
AI
DNA Mtases
Expression
1
0
-1
-24
0
6
12
18
24
30
42
54
96
Time points
Genes
Dnmt5
Rid

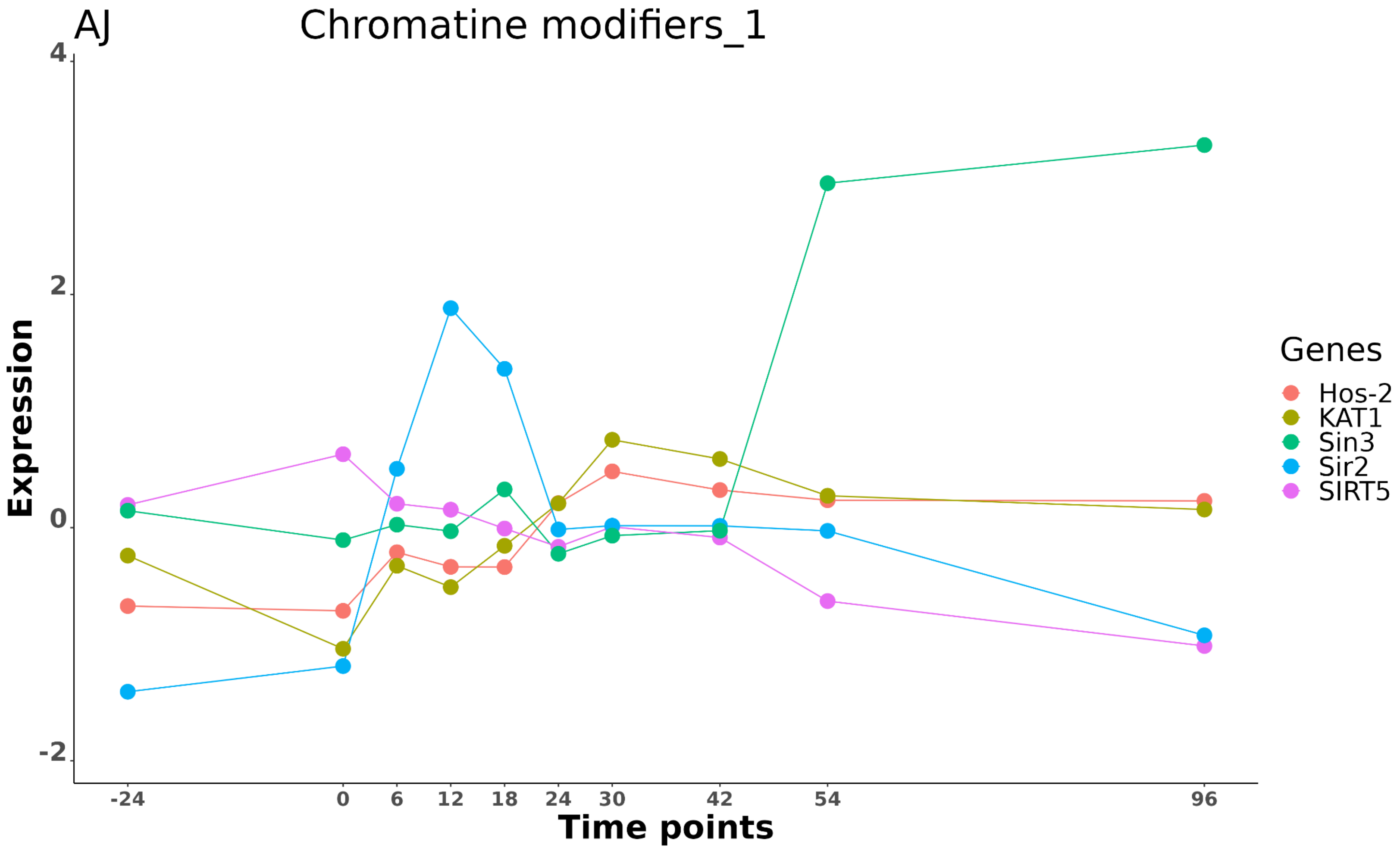
AJ
Chromatine modifiers_1
Expression
4
2
0
-2
-24
0
6
12
18
24
30
42
54
96
Time points
Genes
Hos-2
KAT1
Sin3
Sir2
SIRT5

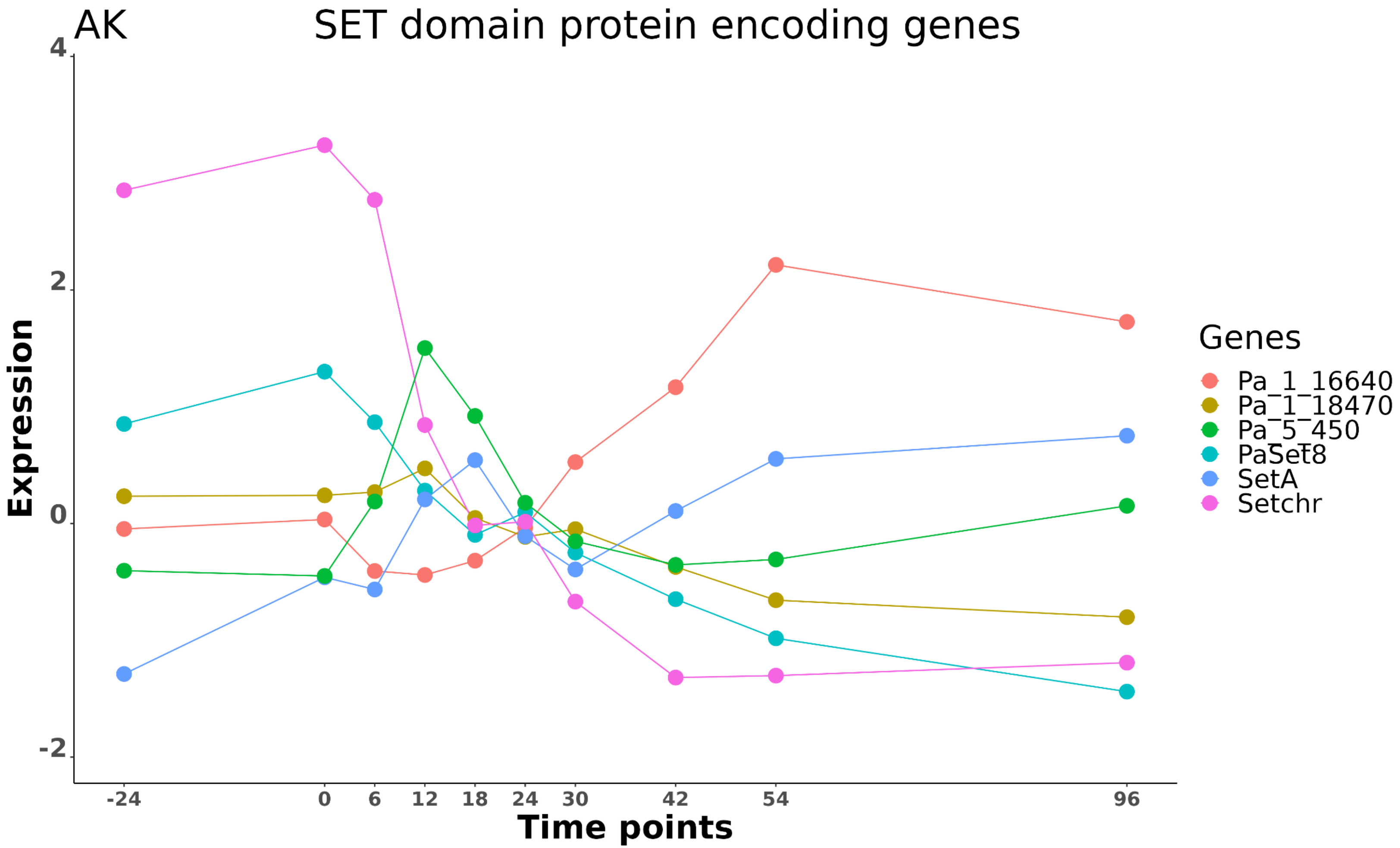
AK
SET domain protein encoding genes
4
2
0
-2
Expression
-24
0
6
12
18
24
30
42
54
96
Time points
Genes
Pa_1_16640
Pa_1_18470
Pa_5_450
PaSet8
SetA
Setchr

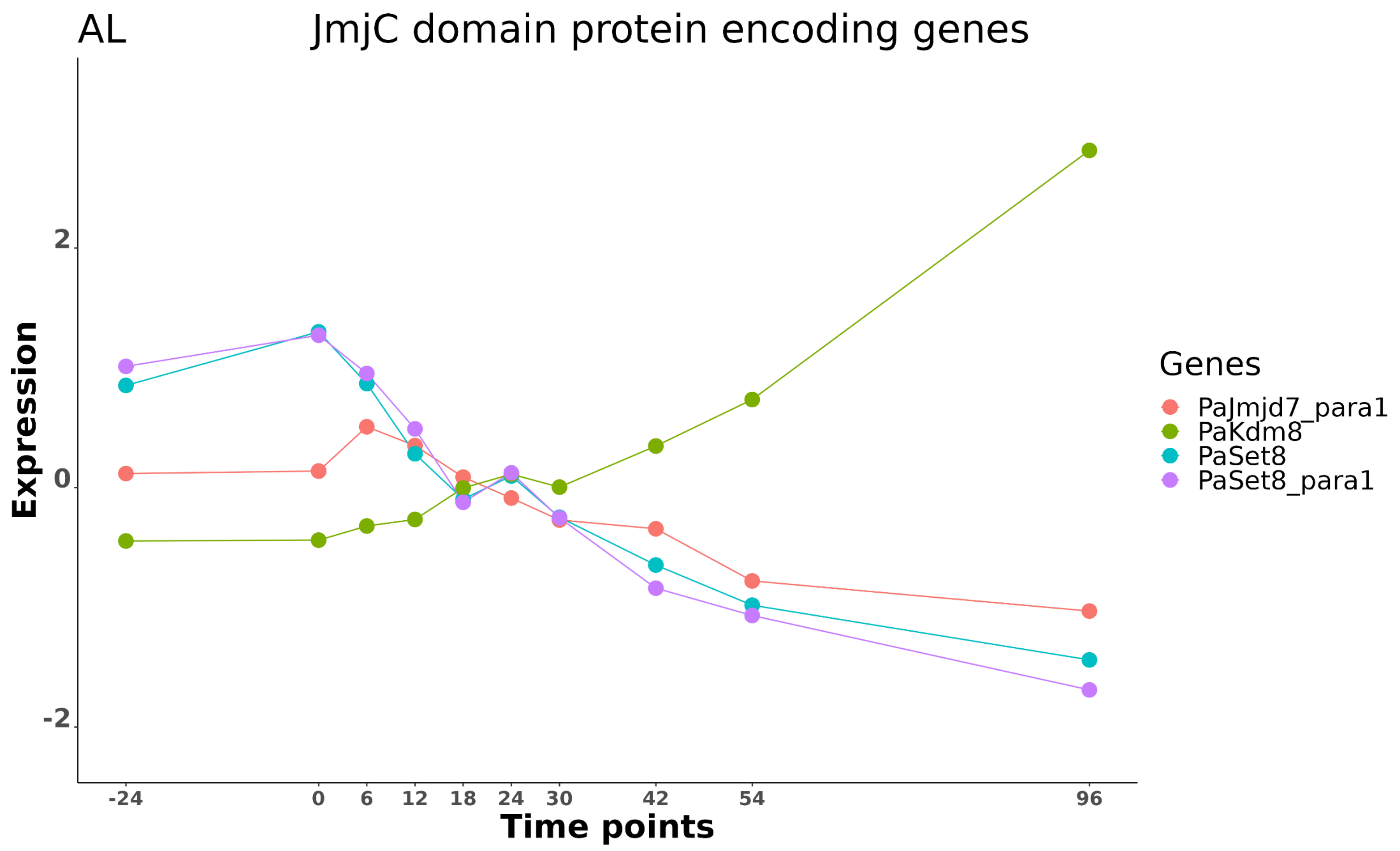

AL
JmjC domain protein encoding genes
Expression
2
0
-2
-24
0
6
12
18
24
30
42
54
96
Time points
Genes
PaJmjd7_para1
PaKdm8
PaSet8
PaSet8_para1

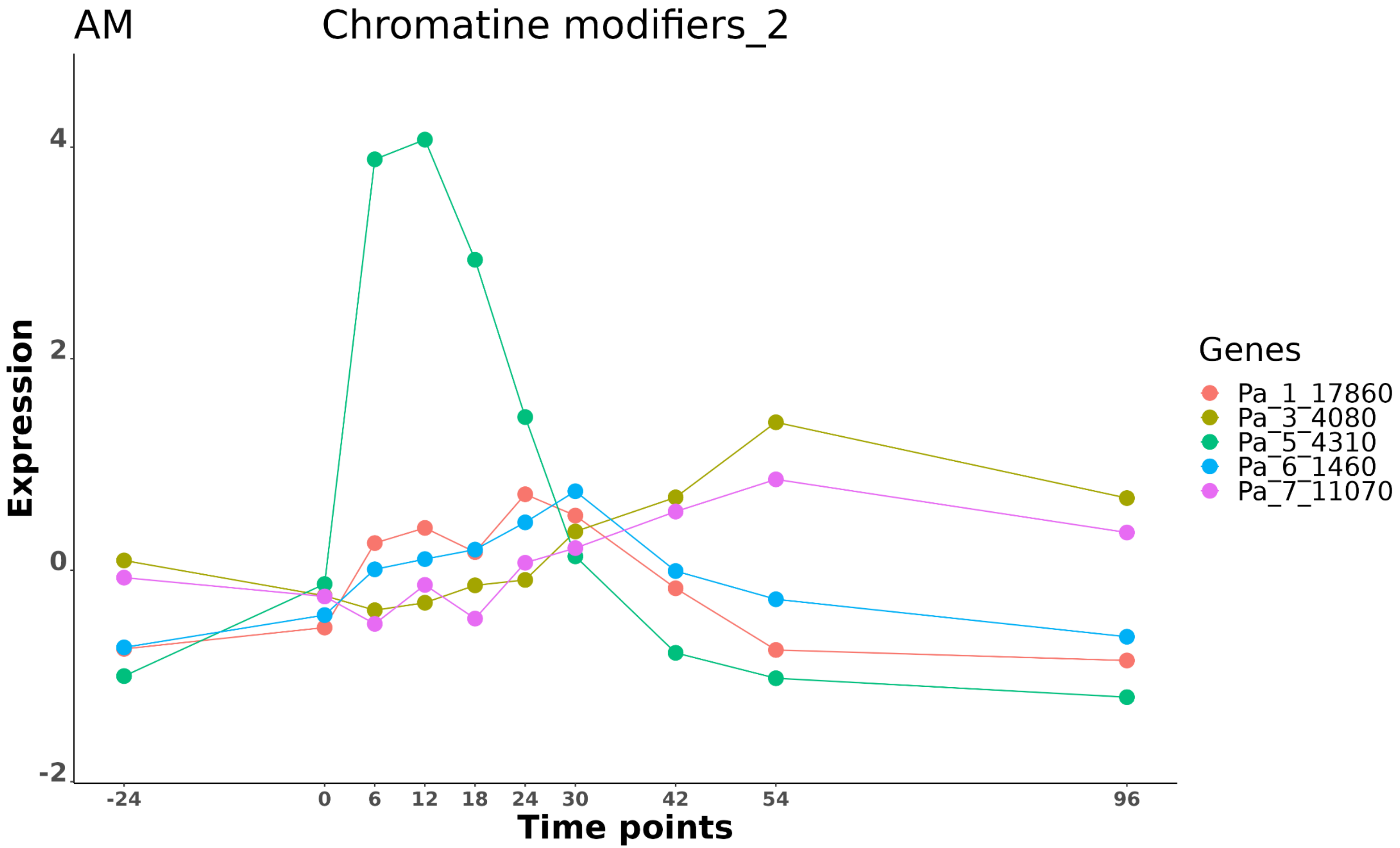

AM
Chromatine modifiers_2
Expression
4
2
0
-2
-24
0
6
12
18
24
30
42
54
96
Time points
Genes
Pa_1_17860
Pa_3_4080
Pa_5_4310
Pa_6_1460
Pa_7_11070

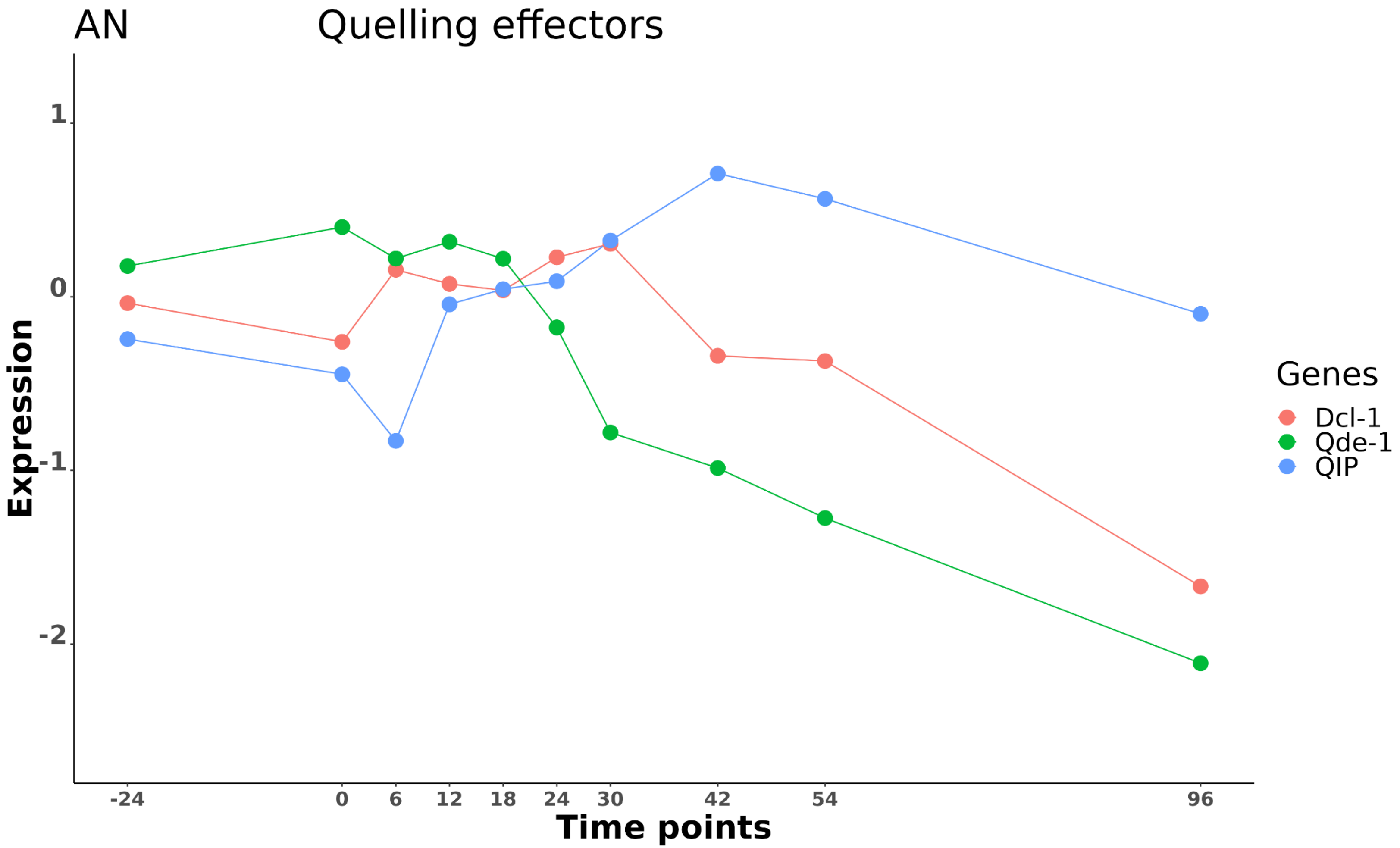
AN
Quelling effectors
1
0
-1
-2
Expression
-24
0
6
12
18
24
30
42
54
96
Time points
Genes
Dcl-1
Qde-1
QIP

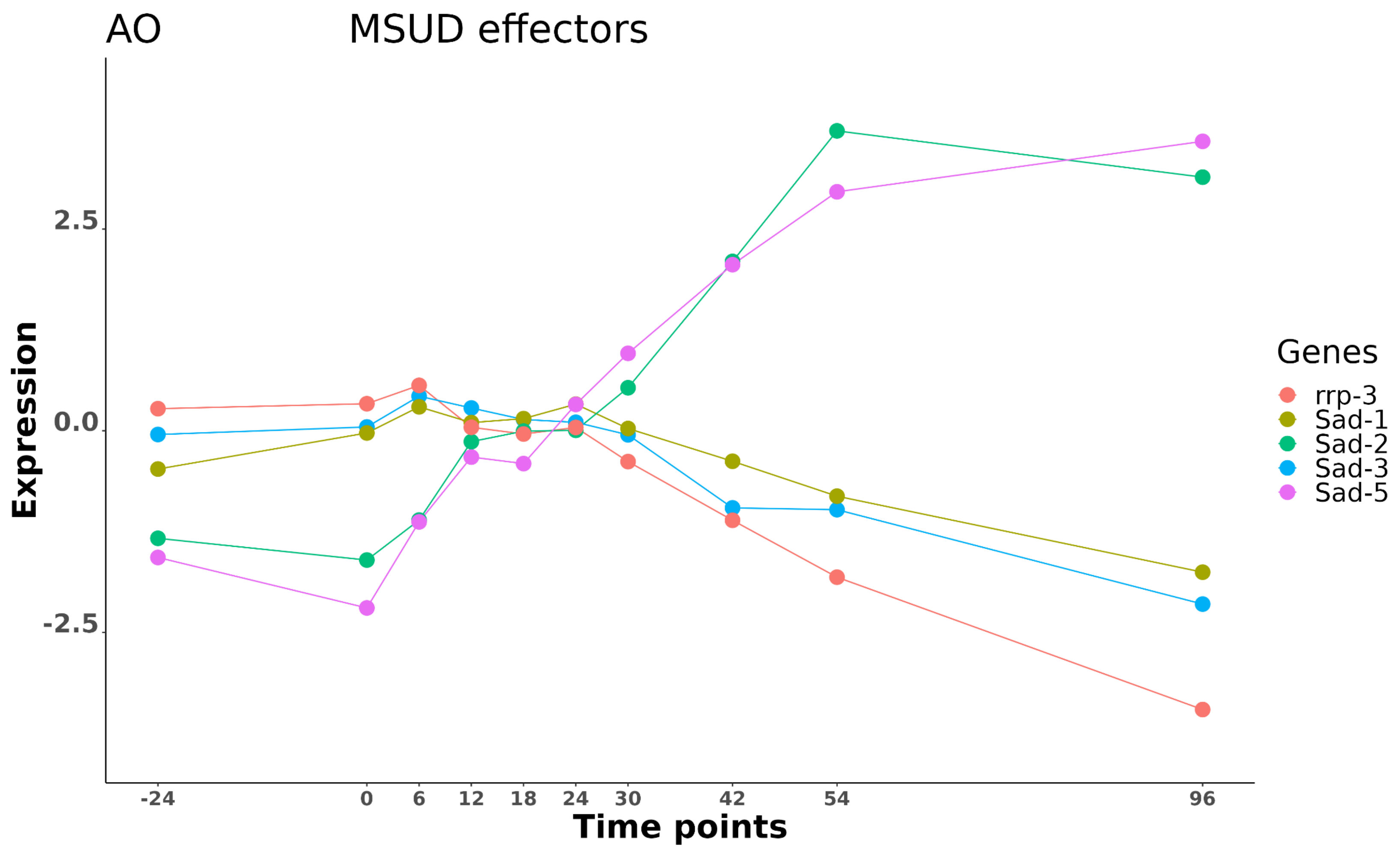

AO
MSUD effectors
Expression
2.5
0.0
-2.5
-24
0
6
12
18
24
30
42
54
96
Time points
Genes
rrp-3
Sad-1
Sad-2
Sad-3
Sad-5

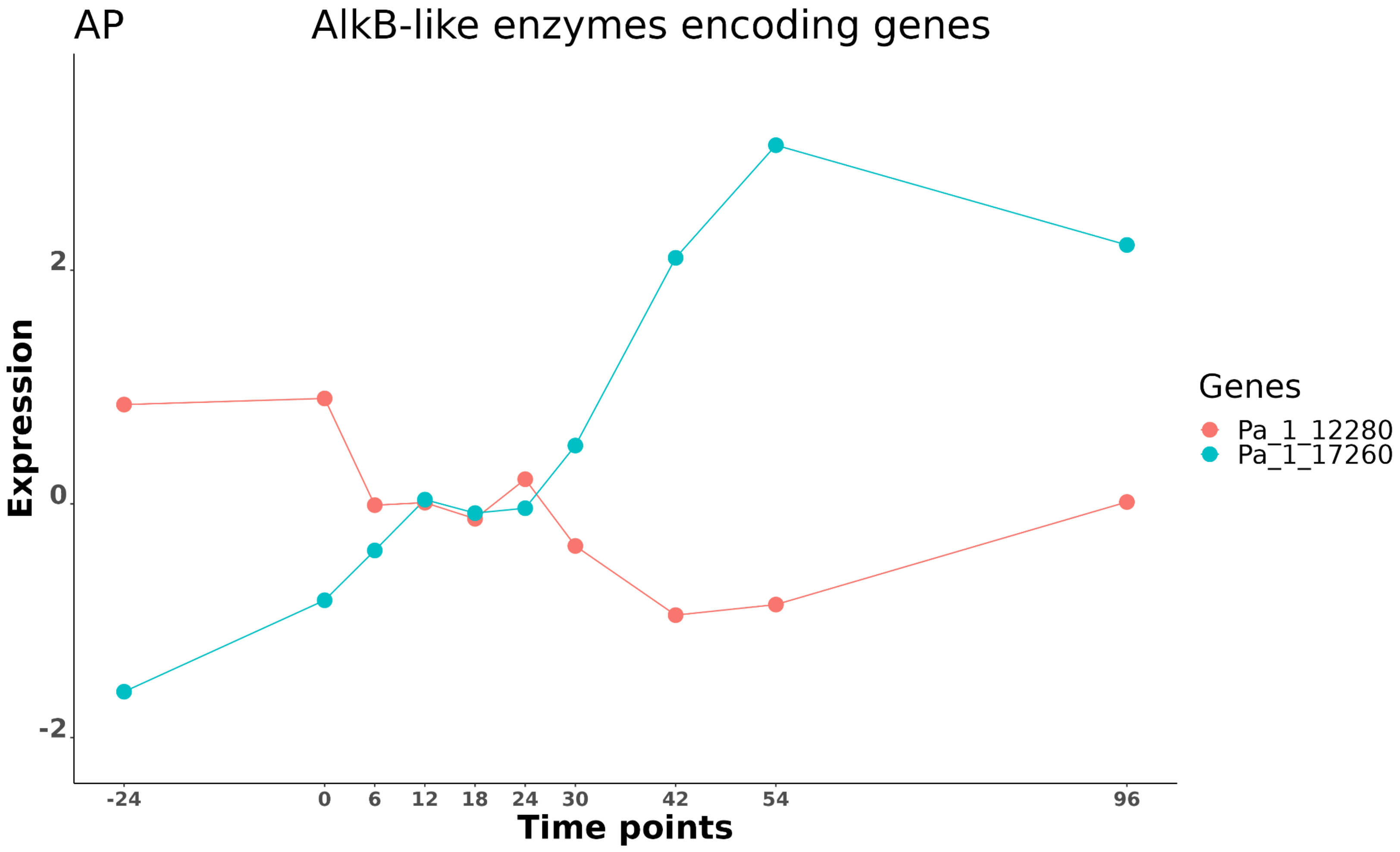

AP
AlkB-like enzymes encoding genes
Expression
2
0
-2
-24
0
6
12
18
24
30
42
54
96
Time points
Genes
Pa_1_12280
Pa_1_17260

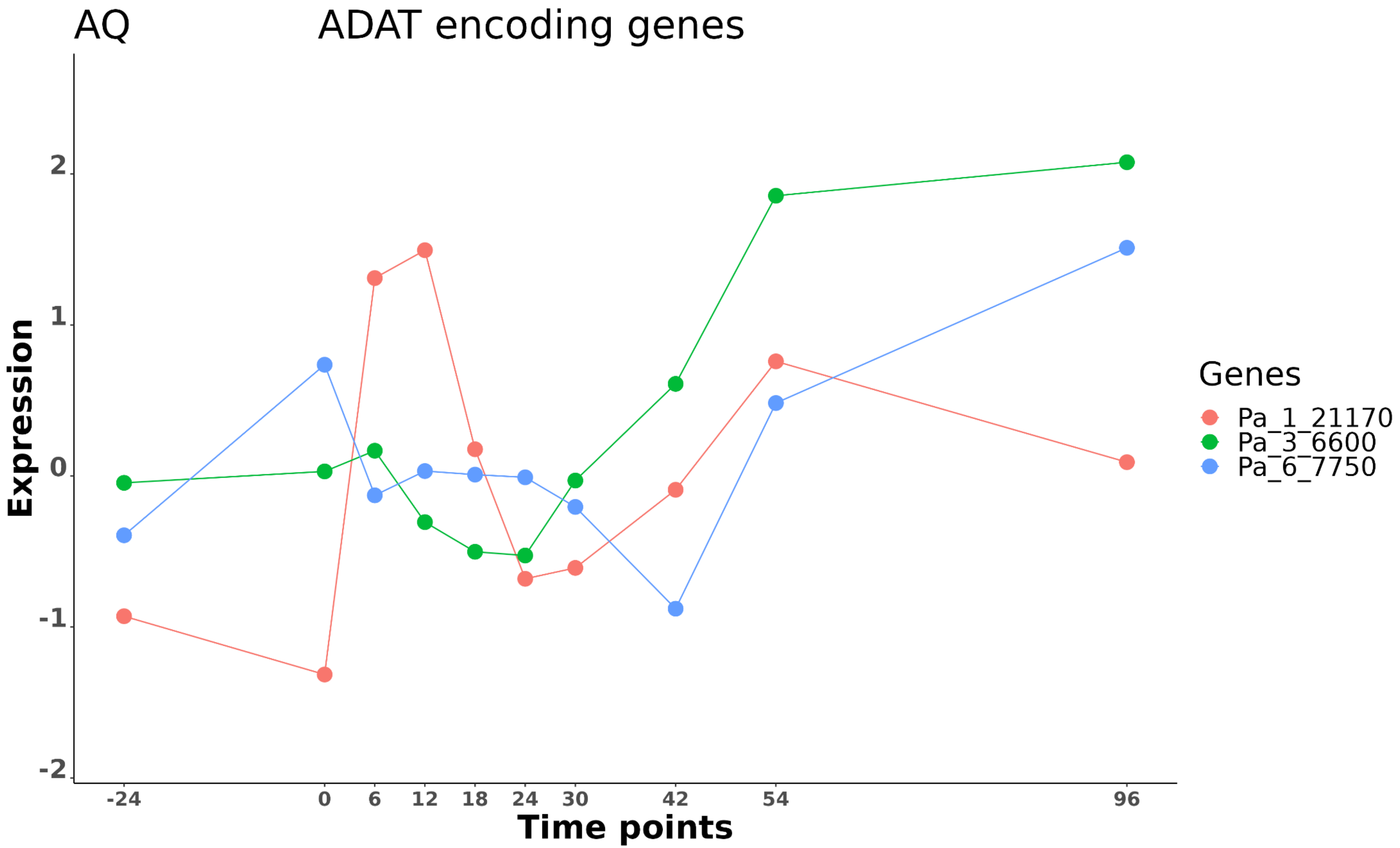

AQ
ADAT encoding genes
Expression
2
1
0
-1
-2
-24
0
6
12
18
24
30
42
54
96
Time points
Genes
Pa_1_21170
Pa_3_6600
Pa_6_7750

**Supplementary Figure 5: Expression profile graphs of genes of interest.**

All graphs presented in this figure were generated using the tool we developed for this study. The WEB application is available at: https://pierregrognet.shinyapps.io/cinetique_app/ (see Methods). X axis: Time point expressed in hours, where 0 is the original time of fertilization. Y axis: Normalized expression profiles of genes (See Methods).

**Supplementary Table 1: List of DE genes.**

**Supplementary Table 2: PCA analysis list of DE genes.** PCA1: first axis. PCA2: second axis. PCA3: third axis. Loadings indicate how a variable contributes to the principal component.

**Supplementary Table 3: Groups of orthologs among DE genes.**

**Supplementary Table 4: multi-class GO-Term analysis. All DE genes.** Class 1 and class 2 GO-term analysis for all of the 3,466 DE genes**. Waves I to V.** Class 1, class 2 and Class 3 GO-term analysis for all of the 1,133 wave DE genes.

**Supplementary Table 5: Pfam analysis and category enrichment. All DE genes.** Pfam search in all of the putative proteins encoded by the 3,466 DE genes. **Waves I to V.** Pfam search in all of the putative proteins encoded by the 1,133 wave DE genes. k: Number of proteins encoded by the DE genes of the considered set that contain this Pfam domain. M: Number of all proteins in the complete set of *P. anserina* CDS that contain this Pfam domain. The categories highlighted in yellow are significantly enriched (adjusted p-value < 0.05%).

**Supplementary Table 6: List of DE genes sorted by functional category.** Annotation column : see FungiDB for name's reference (https://fungidb.org/). F-Secondary Metabolism sheet : the yellow highlighted genes are DE.

**Supplementary Table 7: DE genes involved in the co-expression network centered on transcription factors. A.** List of interactions, cor: coefficient of correlation. **B.** Targets in waves. Numbers indicate the number of putative targets included in each wave. Purple bars provide a graphical visualization of these numbers.

**Supplementary Table 8: Main features of genes accumulating transcripts in *mat-* strain.**

**Supplementary Table 9: Main features of genes accumulating transcripts in *mat+* strain.**

**Supplementary Table 10: DE genes investigated in previous studies in *P. anserina*.**
