## Supplemental Table 8 for "Sexual reproduction is controlled by successive transcriptomic waves in *Podospora anserina*"

**Table S8.** Main features of genes accumulating transcripts in the *mat-* strain.

| **Gene number** | **FC** | **Gene name or function** | **FPR1^a^** | **FMR1^b^** | **DE** | **Wave** |
| --- | --- | --- | --- | --- | --- | --- |
| Pa_1_8290 | -51,54 | *MFM* | 0 | A | - | - |
| FMR1 | -13,91 | *FMR1* | 0 | 0 | - | IV |
| Pa_6_7350 | -6,56 | protease | R | A | YES | II |
| Pa_7_60 | -5,97 | unknown function | 0 | A | YES | - |
| Pa_7_9070 | -5,62 | *PRE1* | 0 | A | - | - |
| Pa_4_1290 | -5,05 | carbohydrate esterase | 0 | A | - | - |
| Pa_5_10930 | -4,72 | unknown function | R | A | - | - |
| nad1 | -4,25 | *nad1* | 0 | 0 | - | - |
| Pa_5_10935 | -4,21 | unknown function | 0 | A | - | - |
| Pa_6_10330 | -3,91 | polyketide synthase | 0 | A | YES | - |
| Pa_2_13500 | -3,72 | unknown function | R | 0 | - | - |
| Pa_6_8610 | -3,62 | unknown function | R | A | YES | IV |
| Pa_3_10350 | -3,61 | unknown function | 0 | 0 | - | - |
| Pa_0_850 | -3,45 | unknown function | 0 | A | YES | IV |
| Pa_5_10945 | -3,27 | unknown function | R | A | - | - |
| Pa_7_875 | -3,11 | unknown function | 0 | A | - | - |
| Pa_5_6620 | -3,08 | cytochrome P450 | R | 0 | YES | - |
| Pa_0_1140 | -3,05 | hydroxyisobutyrate dehydrogenase | 0 | A | - | - |
| nad3 | -3 | *nad3* | 0 | 0 | - | - |
| Pa_1_1310 | -2,93 | unknown function | R | 0 | - | - |
| Pa_6_6400 | -2,89 | glycoside hydrolase | 0 | A | YES | - |
| Pa_7_4140 | -2,88 | gibberellin dioxygenase | 0 | A | - | - |
| Pa_5_6640 | -2,85 | cytochrome P450 | R | A | YES | - |
| Pa_2_1200 | -2,81 | agmatinase 1 precursor | R | A | - | - |
| Pa_1_21320 | -2,69 | unknown function | R | 0 | - | - |
| Pa_3_7460 | -2,67 | alcohol dehydrogenase | R | A | YES | - |
| Pa_1_21830 | -2,66 | biphenyl dioxygenase | R | 0 | - | - |
| Pa_4_1433 | -2,64 | unknown function | 0 | 0 | - | - |
| Pa_5_7120 | -2,62 | unknown function | R | 0 | YES | - |
| Pa_1_6014 | -2,62 | unknown function | 0 | 0 | - | - |
| Pa_5_10940 | -2,61 | unknown function | 0 | A | - | - |
| Pa_1_21970 | -2,6 | peroxisomal alcohol oxidase | 0 | A | - | - |
| Pa_1_21840 | -2,55 | unknown function | R | 0 | YES | - |
| nad2 | -2,53 | *nad2* | 0 | 0 | - | - |
| Pa_5_2930 | -2,52 | glucose transporter | R | 0 | - | - |
| Pa_2_80 | -2,52 | transporter protein | 0 | A | - | - |
| Pa_5_1230 | -2,49 | unknown function | 0 | A | - | - |
| Pa_7_15 | -2,46 | unknown function | 0 | 0 | - | - |
| Pa_1_5700 | -2,4 | potassium transporter | R | 0 | YES | - |
| Pa_1_14680 | -2,37 | 3-ketoacyl-(acyl-carrier-protein) reductase | R | 0 | - | - |
| Pa_5_10190 | -2,36 | glycosyl transferase | R | 0 | YES | - |
| Pa_3_830 | -2,25 | carbohydrate esterase | 0 | 0 | - | - |
| Pa_5_1890 | -2,24 | unknown function | R | 0 | YES | - |
| Pa_2_14160 | -2,2 | lactose permease | 0 | A | - | - |
| Pa_6_6730 | -2,17 | unknown function | R | A | YES | - |
| Pa_1_3100 | -2,16 | unknown function | 0 | A | YES | - |
| Pa_2_8150 | -2,15 | unknown function | 0 | A | - | - |
| Pa_4_4560 | -2,15 | zeaxanthin epoxidase | R | A | YES | - |
| Pa_1_1490 | -2,13 | sorbitol dehydrogenase | R | 0 | - | - |
| Pa_6_11500 | -2,13 | polysaccharide lyase | 0 | 0 | YES | III |
| Pa_7_140 | -2,12 | uracil phosphoribosyltransferase | 0 | 0 | YES | - |
| Pa_1_13200 | -2,12 | unknown function | 0 | 0 | YES | - |
| Pa_3_130 | -2,11 | unknown function | R | 0 | YES | - |
| Pa_5_1620 | -2,1 | NADH oxidase | 0 | A | YES | - |
| Pa_2_4220 | -2,09 | unknown function | R | 0 | YES | II |
| Pa_4_9880 | -2,09 | unknown function | R | 0 | - | - |
| Pa_4_5280 | -2,09 | unknown function | R | 0 | YES | - |
| Pa_1_11070 | -2,08 | glycoside hydrolase | 0 | 0 | YES | - |
| Pa_2_13480 | -2,08 | unknown function | 0 | 0 | YES | IV |
| Pa_3_10430 | -2,08 | unknown function | 0 | 0 | - | - |
| Pa_5_7010 | -2,07 | unknown function | 0 | 0 | YES | II |
| Pa_4_9550 | -2,07 | unknown function | 0 | A | - | - |
| Pa_3_5600 | -2,05 | unknown function | 0 | 0 | - | - |
| Pa_2_13420 | -2,05 | unknown function | 0 | 0 | YES | IV |
| Pa_1_17920 | -2,03 | phosphate transporter | 0 | A | YES | IV |
| Pa_6_10 | -2,02 | unknown function | R | 0 | YES | II |
| Pa_0_1080 | -2,01 | unknown function | 0 | 0 | YES | - |
| Pa_4_110 | -2 | unknown function | 0 | A | YES | - |
| Pa_5_1550 | -2 | unknown function | R | 0 | - | - |

^a^ A: gene activated by FPR1; R: gene repressed by FPR1; 0: gene not controlled by FPR1.

^b^ A: gene activated by FMR1; R: gene repressed by FMR1; 0: gene not controlled by FMR1.

FC = Fold Change

DE = Differentially Expressed
