## Supplemental Table 9 for "Sexual reproduction is controlled by successive transcriptomic waves in *Podospora anserina*"

**Table S9.** Main features of genes accumulating transcripts in the *mat+* strain.

| **Gene number** | **FC** | **Gene name or function** | **FPR1^a^** | **FMR1^b^** | **DE** | **Wave** |
| --- | --- | --- | --- | --- | --- | --- |
| Pa_2_2310 | 144.05 | *MFP* | A | 0 | - | - |
| Pa_4_3858 | 48.16 | unknown function | A | 0 | YES | - |
| Pa_1_24410 | 27.87 | SAM-dependent methyltransferase | A | 0 | - | II |
| Pa_4_1380 | 11.23 | *PRE2* | A | 0 | - | - |
| Pa_5_3435 | 7.76 | unknown function | A | 0 | - | II |
| Pa_7_4100 | 7.5 | unknown function | A | 0 | YES | II |
| Pa_3_3210 | 7.2 | unknown function | A | 0 | - | - |
| Pa_5_9770 | 5.96 | Asp protease | A | 0 | YES | - |
| Pa_5_6960 | 5.94 | unknown function | A | 0 | - | - |
| Pa_1_9625 | 5.08 | unknown function | A | 0 | YES | - |
| Pa_3_1710 | 4.58 | *AOX* | A | 0 | - | - |
| Pa_5_12980 | 3.64 | unknown function | A | 0 | YES | - |
| Pa_1_540 | 3.6 | unknown function | A | 0 | YES | - |
| Pa_4_3160 | 3.54 | phosphoenolpyruvate carboxykinase | A | 0 | YES | IV |
| Pa_6_515 | 3.52 | unknown function | A | 0 | YES | II |
| Pa_1_18670 | 3.43 | unknown function | 0 | 0 | - | - |
| Pa_2_13290 | 3.42 | unknown function | A | 0 | - | - |
| Pa_1_16335 | 3.28 | unknown function | 0 | R | YES | - |
| Pa_4_1370 | 3.06 | amidohydrolase | A | 0 | YES | - |
| Pa_4_7760 | 3,03 | farnesyltransferase subunit beta | A | 0 | YES | II |
| Pa_7_9690 | 2,95 | protein-S-isoprenylcysteine O-methyltransferase | A | R | - | - |
| Pa_7_1820 | 2,79 | mitochondrial NADH-ubiquinone oxidoreductase 1 | A | 0 | - | - |
| Pa_1_20140 | 2,77 | unknown function | A | 0 | - | - |
| Pa_5_4585 | 2,73 | unknown function | A | 0 | - | - |
| Pa_4_9360 | 2,67 | fructose-1,6-bisphosphatase | 0 | 0 | YES | - |
| Pa_2_8800 | 2,64 | unknown function | A | 0 | YES | V |
| Pa_4_7450 | 2,62 | ketopantoate hydroxymethyltransferase | A | 0 | YES | II |
| Pa_6_540 | 2,61 | unknown function | 0 | 0 | YES | V |
| Pa_6_9760 | 2,56 | nonribosomal peptide synthetase | A | 0 | - | - |
| Pa_1_19170 | 2,54 | unknown function | 0 | 0 | - | - |
| Pa_6_9980 | 2,53 | unknown function | A | 0 | YES | II |
| Pa_7_2860 | 2,5 | cyclic-nucleotide phosphodiesterase | A | 0 | - | - |
| Pa_5_820 | 2,49 | unknown function | 0 | 0 | - | - |
| Pa_6_350 | 2,48 | plasma membrane proteolipid 3 | A | 0 | YES | - |
| Pa_2_5500 | 2,48 | cation-transporting ATPase | A | R | - | - |
| Pa_2_9145 | 2,46 | unknown function | A | 0 | YES | - |
| Pa_5_7250 | 2,45 | unknown function | A | 0 | YES | V |
| Pa_0_690 | 2,42 | acyl CoA thioesterase | A | 0 | YES | - |
| Pa_2_7170 | 2,4 | glycosyltransferase | A | 0 | - | - |
| Pa_6_11620 | 2,4 | methionine permease | A | 0 | YES | - |
| Pa_2_4810 | 2,4 | unknown function | A | 0 | YES | - |
| Pa_2_7180 | 2,39 | unknown function | A | 0 | - | - |
| Pa_5_9600 | 2,39 | isovaleryl-CoA dehydrogenase | A | 0 | YES | - |
| Pa_4_80 | 2,37 | unknown function | 0 | 0 | - | - |
| Pa_5_12470 | 2,36 | unknown function | 0 | 0 | YES | I |
| Pa_3_8590 | 2,36 | cytochrome c | A | 0 | - | - |
| Pa_2_9500 | 2,36 | unknown function | A | 0 | - | - |
| Pa_1_30 | 2,35 | unknown function | 0 | 0 | YES | - |
| Pa_5_9790 | 2,35 | unknown function | A | 0 | YES | IV |
| Pa_5_11640 | 2,34 | ABC transporter | 0 | 0 | YES | - |
| Pa_5_11460 | 2,32 | abhydrolase | A | 0 | - | - |
| Pa_7_2870 | 2,31 | unknown function | A | 0 | - | II |
| Pa_0_1270 | 2,29 | MSF superfamily | A | A | YES | II |
| Pa_1_20590 | 2,27 | *FPR1* | 0 | 0 | - | - |
| Pa_4_860 | 2,27 | unknown function | A | 0 | YES | - |
| Pa_1_5530 | 2,27 | unknown function | A | 0 | YES | - |
| Pa_2_6010 | 2,26 | cholesterol oxidase | A | 0 | - | - |
| Pa_2_6830 | 2,26 | C6 transcription factor | A | 0 | YES | - |
| Pa_5_7750 | 2,25 | polyketide synthase | A | 0 | - | - |
| Pa_1_10600 | 2,25 | mitochondrial deoxynucleotide carrier | A | 0 | - | - |
| Pa_1_12800 | 2,24 | unknown function | A | 0 | - | - |
| Pa_7_950 | 2,24 | unknown function | 0 | 0 | - | - |
| Pa_1_8280 | 2,23 | unknown function | 0 | 0 | YES | - |
| Pa_1_15470 | 2,23 | laccase | A | 0 | - | - |
| Pa_3_1990 | 2,22 | polyketide synthase | 0 | 0 | YES | - |
| Pa_2_3690 | 2,22 | unknown function | A | R | - | - |
| Pa_0_1190 | 2,2 | molybdenum cofactor sulfurase | 0 | A | YES | II |
| Pa_2_10220 | 2,19 | Glutamine synthetase | A | 0 | YES | - |
| Pa_7_3250 | 2,18 | unknown function | A | 0 | - | - |
| Pa_4_5450 | 2,17 | unknown function | A | R | YES | - |
| Pa_5_3130 | 2,16 | esterase/lipase | A | 0 | YES | - |
| Pa_4_9520 | 2,16 | copper fist DNA binding domain protein | A | 0 | - | - |
| Pa_1_1230 | 2,14 | unknown function | 0 | 0 | YES | V |
| Pa_4_3860 | 2,09 | esterase/lipase | A | 0 | YES | - |
| Pa_2_1170 | 2,09 | unknown function | A | 0 | - | - |
| Pa_2_11120 | 2,08 | malic enzyme | A | 0 | - | - |
| Pa_1_15590 | 2,08 | unknown function | A | 0 | - | - |
| Pa_6_3770 | 2,06 | C6 transcription factor | A | 0 | YES | - |
| Pa_1_15330 | 2,06 | membrane transport protein | A | 0 | - | - |
| Pa_2_50 | 2,05 | mannose-6-phosphate isomerase | A | 0 | YES | - |
| Pa_1_20510 | 2,05 | sugar transporter | A | R | YES | IV |
| Pa_4_230 | 2,04 | unknown function | 0 | 0 | YES | II |
| Pa_5_570 | 2,04 | unknown function | 0 | 0 | YES | - |
| Pa_1_22300 | 2,03 | glycine dehydrogenase | A | 0 | - | - |
| Pa_4_3210 | 2,03 | unknown function | A | 0 | - | - |
| Pa_2_7590 | 2,02 | unknown function | 0 | 0 | YES | - |
| Pa_3_720 | 2,01 | unknown function | A | 0 | - | - |
| Pa_2_5340 | 2,01 | esterase/lipase | A | 0 | - | - |

^a^ A: gene induced by FPR1; R: gene repressed by FPR1; 0: gene not controlled by FPR1.

^b^ A: gene induced by FMR1; R: gene repressed by FMR1; 0: gene not controlled by FMR1.

FC = Fold Change

DE = Differentially Expressed
